# HIV-exposure related disruptions in functional and structural connectivity in the central auditory system in adolescence

**DOI:** 10.64898/2026.04.06.716813

**Authors:** Joanah Madzime, Marcin Jankiewicz, Ernesta M. Meintjes, Peter Torre, Barbara Laughton, Martha J. Holmes

## Abstract

**Background:** Children who are HIV-exposed but uninfected (CHEU) face elevated risks of hearing loss and language deficits compared to HIV-unexposed peers. The central auditory system (CAS) undergoes substantial maturational changes during adolescence, yet no neuroimaging study has examined its structural or functional integrity in CHEU. Prior work in this cohort identified white matter (WM) alterations in regions adjacent to the CAS at age 7, and reduced auditory working memory in CHEU relative to unexposed children (CHUU).

**Aim:** To characterise WM integrity and functional connectivity (FC) of the CAS and related regions in CHEU at age 11, to investigate structural and functional network topology, and to examine associations between imaging outcomes and neurocognitive function.

**Methods:** Forty-eight children aged 11–12 (20 CHEU, 28 CHUU) from an ongoing longitudinal neurodevelopmental cohort underwent 3T MRI including diffusion tensor imaging (DTI) and resting-state fMRI (RS-fMRI). CAS regions (cochlear nucleus/superior olivary complex, inferior colliculus [IC], medial geniculate nucleus [MGN], and primary auditory cortex [PAC]) were manually segmented and combined with an automated atlas. DTI probabilistic tractography was performed, extracting FA, MD, AD, RD, fractional number of tracts, and tract volume. FC was computed using Pearson correlations between regional time series. Graph theory measures (degree, strength, transitivity, nodal and local efficiency) were derived for structural and functional networks. RS-fMRI group comparisons used Bayesian multilevel modelling (matrix-based and region-based analyses), while DTI comparisons used linear models with FDR correction. Neurocognitive testing employed the KABC-II.

**Results:** No significant group differences in DTI WM metrics (FA, MD, AD, RD) were observed after FDR correction. CHEU demonstrated higher structural nodal strength in the left IC (FDR-significant) and in the bilateral rostral middle frontal cortex (rMFC) and right cuneus. RS-fMRI revealed lower FC between the bilateral IC in CHEU, alongside reduced FC in the left caudate, left hippocampus CA3, left pericalcarine, and left lingual gyrus. CHEU showed higher FC between the left MGN and right precentral, left postcentral, and right rMFC; the right PAC also showed higher FC to the right rMFC and left postcentral gyrus. No significant group differences were observed in functional nodal measures. No significant associations were found between structural or functional imaging outcomes and neurocognitive scores after multiple comparison correction.

**Discussion:** Structural and functional alterations within the CAS were most prominent in the IC, with increased nodal strength in CHEU potentially reflecting compensatory structural connectivity, and reduced interhemispheric FC between the bilateral IC suggesting disrupted auditory integration. Altered FC between the MGN/PAC and cortical regions, including the rMFC and sensorimotor cortices, may reflect differences in top-down auditory processing. The absence of imaging-cognition associations at age 11 suggests that these connectivity differences do not, at this stage, translate into measurable deficits in auditory or language-related neurocognitive performance.

**Conclusion:** This is the first study to examine functional and structural connectivity of the CAS in CHEU children. HIV exposure is associated with subtle but discernible alterations in IC connectivity and in CAS links to cortical regions at age 11, without detectable neurocognitive correlates. Longitudinal follow-up and inclusion of audiological and ART exposure data are needed to clarify the developmental and functional consequences of these findings.

## Introduction

The central auditory system (CAS) is made up of structures and pathways involving the brainstem, midbrain, thalamus and cortex that allow for the understanding and interpretation of sounds. Adolescence, defined by the WHO as 10 – 19 years, is a period of substantial maturational changes for the central nervous system. During adolescence, the combination of increased myelination and synaptic pruning is thought to lead to more defined and complex neural networks (Asato et al., 2010; Casey et al., 2008; Paus, 1999). As a result, neural processing (including auditory) continues to mature during this period (Albrecht et al., 2000; Krizman et al., 2015; Ponton et al., 2000).

Accessibility to combination antiretroviral therapy (cART) has reduced the transmission of human immunodeficiency virus (HIV) from mother to child by 54% in the last decade (UNAIDS, 2021). As a result, there is an increasing number of children exposed to HIV in utero and uninfected. (CHEU). Infants and children exposed to HIV are at a higher risk of hearing loss (Torre et al., 2012) as well as language deficits (Redmond et al., 2016; Van Rie et al., 2008; Wedderburn et al., 2019), compared to their unexposed peers. Research into the peripheral auditory system (PAS) in CHEU has reported links between disruptions and hearing loss. However, these findings do not provide a full picture of HIV and cART exposure-related auditory and language abnormalities. A better understanding of HIV-exposure on the CAS may provide additional insight.

Diffusion tensor imaging (DTI) and resting-state functional MRI (RS-fMRI) allow for investigating white matter microstructure and function of brain regions, respectively (Basser et al., 1994; van den Heuvel & Hulshoff Pol, 2010). Outcomes of DTI include fractional anisotropy (FA), a measure related to axonal integrity, and mean diffusivity (MD), which is a measure of axonal maturation (Feldman et al., 2010; Ranzenberger & Snyder, 2022). RS-fMRI measures the extent of co-activation between anatomically separate gray matter regions (Biswal et al., 1997; Yang et al., 2020), referred to as functional connectivity (FC). Recently, both DTI and RS-fMRI have been combined with graph theoretical frameworks to investigate the brain as structural and functional networks, respectively (Sporns, 2014). In these networks we can investigate the integration and segregation between brain regions, so called nodes, ultimately revealing the arrangement or topology of the network and its regions (Bullmore et al., 2016).

DTI and RS-fMRI studies in CHEU are still accumulating and none have focused on the CAS. To date, imaging studies haven’t reported HIV-exposure-related alterations in CAS structures. However, there is evidence of changes in pathways and structures that work together with the CAS. In CHEU, reduced FA in the supramarginal gyrus, important for phonological processing, and the middle temporal gyrus was reported in comparison to children who are HIV uninfected and unexposed (CHUU) (Yadav et al., 2020). Tran and colleagues reported higher FA in the cerebellar peduncles of HEU infants and found associations between FA of the uncinate fasciculus, a white matter (WM) fiber connecting deep temporal regions to subcortical regions, and neurodevelopmental scores (Tran et al., 2016).

This work builds on an ongoing neurodevelopmental study of children infected with and exposed to HIV and followed since age 5. Within this cohort, a cross-sectional study that compared 7-year-old CHEU to their age-matched CHUU peers, found higher FA in CHEU in a cluster of the corona radiata, a WM fiber that is linked to the medial geniculate nucleus (MGN) (Jankiewicz et al., 2017). At the same age, higher FA was reported in the superior longitudinal fasciculus (SLF) in the motor resting state network of CHEU compared to CHUU (Madzime et al., 2021). Investigations into the neurodevelopmental outcomes of this cohort show comparable outcomes between CHEU and CHUU at age 5 and similar age-dependent improvements in most neurodevelopmental outcomes between 7 and 9 years old. However, CHUU had higher mean auditory working memory scores compared to CHEU at age 7 years old (Laughton et al., 2018; van Wyhe et al., 2021).

Motivated by the ongoing risks of hearing and language disruptions in CHEU, as well as exposure associated alterations to regions relating to the CAS, we used DTI and RS-fMRI to identify the potential effects of HIV exposure on WM integrity and FC of the CAS and related regions in CHEU at age 11. Further, we investigated the regional network properties of the CAS and related regions. Finally, we looked at associations between imaging outcomes and neurodevelopmental measures.

## Methods

### Study cohort

Participants were 48 11–12-year-old children (20 CHEU, 28 CHUU) who participated in an ongoing neurodevelopmental study that included neuroimaging and cognitive testing. The children were recruited during infancy as part of a vaccine study (Madhi et al., 2010) or as controls on the CHER follow-on neurodevelopmental studies. The study protocol was approved by the Human Research Ethics Committees of the University of Cape Town and University of Stellenbosch. Parents/guardians provided written informed consent and children provided assent., in person in their preferred language. Children were first familiarized with the scanning procedures on a mock scanner.

### Image acquisition

All imaging was done in the same session using a 32-channel head coil on a 3 Tesla Siemens Skyra MRI scanner (Erlangen, Germany) at the University of Cape Town’s Cape Universities Body Imaging Centre (CUBIC). **T1-weighted (T1w) structural** volumes were acquired for 6 mins using a multiecho magnetization prepared rapid gradient echo (MEMPRAGE) sequence with TR = 2530 ms, TEs = 1.69/3.54/5.39/7.24ms, inversion time (TI) 1,100ms, flip angle = 7°, and FOV = 224×224×176mm^3^. **Diffusion tensor imaging (DTI)** comprised two whole-brain diffusion-weighted (DW) acquisitions with the phase encoding direction (anterior-posterior in our case) inverted to correct for susceptibility artefacts (7 mins each). Acquisition parameters for DTI were: TR/TE 11100/92ms, 84 slices, 2×2×2mm^3^, FOV = 244×244×168 mm^3^, 30 non-collinear diffusion directions, b=1000s/mm^2^, and five non-diffusion-weighted (b0) acquisitions. **RS-fMRI** data were acquired for 6 mins using an interleaved multi-slice 2D gradient echo, echo planar imaging (EPI) sequence with the parameters: 33 interleaved slices, slice thickness 4mm, slice gap 1mm, voxel size 3×3×4mm^3^, FOV = 250×250×164mm^3^, TR/TE = 2,000/30ms, flip angle 90°, 180 EPI volumes.

### Neurocognitive Testing

Neurocognitive testing was performed at the Family Centre for Research with Ubuntu (FAMCRU), Stellenbosch University. Participants were administered the Kaufman Assessment Battery for Children – Second Edition (KABC-II) (Kaufman & Kaufman, 2014) in the language of their choice, by trained research assistants supervised by a licensed psychologist. The main outcomes of the battery are 1) Learning, which is a measure of the ability to store and retrieve information; 2) Planning, which assesses the ability to solve nonverbal problems; 3) Sequential processing, which asses the ability to solve problems using auditory and visual information presented serially, and 4) Simultaneous processing, which is a measure of the ability to solve spatial or logistical problems that require simultaneous processing of multiple related stimuli (Kaufman & Kaufman, 2014). Outcomes measured via various subtests are appropriately combined to compute two global scores, namely the Mental Processing Index and Nonverbal Index. In the current study we chose the subtests that are involved in auditory and/or language processing which were Number recall (pure auditory, working memory and attention); Word order (visual and short-term memory); Atlantis, Atlantis delayed, Rebus, Rebus delayed and Delayed recall (learning, long-term storage and retrieval); Story completion (visuospatial); and Sequential processing (auditory working memory). We added a verbal fluency test that is not part of the KABC, Animal naming, which tests word generation and categorization fluency (Strauss et al., 2010).

### Image pre-processing

The processing described here was done for each subject using Analysis of Functional Neuroimages (AFNI) (Cox, 1996), Functional and Tractographic Connectivity Analysis Toolbox (FATCAT) (Taylor & Saad, 2013)and Freesurfer image analysis suite (FreeSurfer v7.1.0: http://surfer.nmr.mgh.harvard.edu/). Images were first converted from DICOMs to NIFtI readable format.

#### T1w structural volume

Using the structural image we performed both manual segmentation of CAS regions and whole-brain automatic segmentation. We performed manual segmentation for the CAS regions as most automatic segmentation schemes do not include the CAS regions because of their sizes and anatomical locations. The manual segmentations were done by an experienced neuroanatomist using Freeview software from FreeSurfer on a Lenovo ThinkPad Yoga370 tablet. In total the CAS regions were four bilateral regions including a region at the junction between the brainstem pons and medulla and consisting of the cochlear nucleus and superior olivary complex (CN/SOC), the inferior colliculus (IC) in the midbrain, the MGN in the thalamus and the primary auditory cortex (PAC) in the superior temporal lobe. Automatic segmentation was performed using Connectome mapper (CMP) 3 (v3.0.0-RC4), an image processing pipeline that produces multi-resolution morphological subject atlases (Daducci et al., 2012; Tourbier et al., 2022). We used the lowest resolution which consisted of a set of 126 subcortical and cortical ROIs. We then merged the manually traced and automatically segmented regions. In locations where a manually traced region overlapped with an automatically segmented region, we excluded the entire automatically segmented region to keep the manually traced one. We also excluded white matter and ventril atlas segmentations. This resulted in a final whole brain structural atlas of 81×81 gray matter regions for each subject. A list of all the regions in the atlas and their abbreviations can be found in Appendix A: Automatically Segmented Regions.

#### DTI data

First, we visually inspected DW images (in both Anterior-Posterior, AP and Posterior-Anterior, PA directions) and manually removed all volumes with dropout slices. Any volume removed from the AP acquisition was also removed from the PA acquisition. Subjects with less than 15 DW volumes or at least one b0 volume were excluded. We then performed motion correction, corrected distortions arising from eddy currents and B0 inhomogeneities using the Tolerably Obsessive Registration and Tensor Optimization Indolent Software Ensemble (TORTOISE; version 3.2) in AFNI (Pierpaoli et al., 2009), and co-registered each child’s DTI volumes to his/her structural volume. As a structural reference image, we used a T2w-contrast image created by imitating the T1w structural image. After distortion correction, diffusion tensors (DT) were solved in each voxel and parameter maps generated. These included FA, MD, axial- (AD) and radial diffusivities (RD). We then inflated the structural atlas regions by 4 voxels to bring them closer to the adjacent white matter, as voxels closest to gray matter regions are typically below tractography thresholds. Each subject white matter mask was used as a reference dataset to prevent inflation into voxels labelled white matter. Subsequently, we performed ***probabilistic tractography*** using FATCAT (version 1.1) (Taylor & Saad, 2013) to identify WM tracts between the inflated structural atlas regions. Stopping criteria for tractography were FA < 0.2, angle > 50 degrees, length < 10 mm. Monte Carlo simulations were run with 5 starting seeds per voxel per iteration and 1000 iterations. Outputs included, for every DTI parameter (FA, MD, AD and RD, an adjacency matrix of means and standard deviations for the WM tracts connecting each pair of regions. Adjacency matrices were also generated for fractional number of tracts (fNT), defined as the total number of probabilistic streamlines in the tracked WM bundle divided by the total number of streamlines in the subject’s whole brain, and tract physical volume (PV) in mm^3^.

#### RS-fMRI data

We began by removing the first 5 EPI volumes to account for the time needed for signal stabilization and ‘despiked’ each remaining volume to truncate the signal in voxels with artificially high signal. After, we performed a temporal alignment of the volumes so each slice’s acquisition time would be the same as the beginning of the volume. The volumes were then aligned to both the reference EPI (the sixth volume) and to the participant’s own T1w structural volume. For each rs-fMRI volume, the motion correction algorithm outputs a summary measure of the within-TR motion called Enorm. Volumes with Enorm > 0.3 mm per TR were removed. Subsequently, we performed spatial smoothing with a Gaussian kernel of 6 mm full width at half maximum (FWHM) and discarded voxels outside the brain. We regressed the mean WM and cerebrospinal fluid (CSF) values as well as the motion parameter estimates and their derivatives out of the EPI data to extract fluctuations in the data that are not BOLD signal. Finally, each subject’s dataset was truncated to 160 consecutive EPI volumes which corresponded to the minimum number of usable volumes from the set of all subjects. Subjects with less than 160 EPI volumes were excluded. We used this “clean” dataset **to calculate FC** between structural atlas regions. We did this by calculating Pearson correlation coefficients between the mean time series of every pair of regions (*3dNetCorr)* (Taylor & Saad, 2013). The correlation coefficients were Fisher Z-transformed allowing an estimation of a normal distribution of the correlations.

### Structural and functional brain network definition

To define the **structural brain network**, we extracted each subject’s fNT matrix and identified regions that were common to all subjects; individual subject connectomes were created based on this set of regions. The total number of common regions were 71 resulting in each subject’s final matrix being 71×71. Since fNT represents a normalised value of the probabilistic streamlines between nodes, we did not threshold the structural brain network.

To define the **functional brain network**, we extracted each subject’s correlation matrix and identified 80 ROIs that were present in all subjects. This resulted in each subject’s final FC correlation matrix being 80×80. Since weak (i.e. Fisher Z-transformed r values < 0.1003 equivalent to Pearson-r < 0.1000) or spurious connections tend to obscure functional network topology (Fornito et al., 2016; Rubinov & Sporns, 2010), such edges were set to zero. Currently, there is no consensus on how to interpret and treat negative edges in graph analyses (Hallquist & Hillary, 2018). To allow for an intuitive interpretation of the relationship between nodes (i.e. strong positive weights mean nodes co-activate), we chose here to set negative edge weights to zero.

**For both structural and functional networks** we used iGraph (v 1.3.1) and brainGraph (v 3.0) packages in R (v 4.1.3) to calculate, for each node, 5 nodal graph theory measures. We calculated node degree and strength which measure the density of connections to a node and indicate the nodes that are likely hub regions in a network (Hoff et al., 2013; van den Heuvel & Sporns, 2013). We calculated node transitivity which measures the degree to which the node falls into a segregated cluster with the nodes around it potentially representing functional coordination (Fornito et al., 2016). Then we calculated nodal efficiency which indicates the extent to which a node integrates or interacts with all other nodes in the brain and, finally, nodal local efficiency which indicates the interconnectedness of a node’s neighbours in the event of damage to the node (Fornito et al., 2016). High nodal local efficiency reflects efficient integration with their neighbors (Fornito et al., 2016).

### Statistical analyses

#### Structural Brain Networks

To perform statistical analysis on DTI tractography data we identified the probabilistic WM connections that were present in at least 95% of the sample. These were 498 WM connections, 18 of which were between CAS regions and 109 that were with at least one CAS region. For these connections we extracted for each subject the statistics for the DTI parameters: FA, AD, RD, MD, fNT and PV.

Then, **for each connection and node in the structural network**, we identified outliers in any DTI parameter or nodal measure, respectively. Outliers were defined as values lying more than 1.5 times the interquartile range (IQR) above or below the upper (Q_U_) or lower (Q_L_) quartiles, respectively. We subsequently ran linear models for the set of connections and nodes to compare groups, CHUU vs CHEU. If including the outliers in the group comparison resulted in a different result than when not included, we excluded the outliers from the analysis. In addition, though both males and females experience significant WM maturation during adolescence, sex differences in trajectory rates have been observed (Gur & Gur, 2016; Simmonds et al., 2014). As such we included sex as a potential confounding variable in all models. Additionally, we included intracranial volume (ICV) as a confounding variable to control for head size differences and examined handedness as a potential confounder. We corrected for multiple comparisons using the false discovery rate (FDR) method. We chose an arbitrary FDR q-value of 0.05 as mark of significance.

#### Functional Brain Networks

To investigate **FC and functional nodal integrity** we employed Bayesian multilevel modelling (BML) (Chen, Bürkner, et al., 2019; Chen, Xiao, et al., 2019). Each subject’s final FC matrix was 80×80 meaning 3160 connections or region pairs (RP). In the traditional null hypothesis significance testing (NHST) model this would mean calculating, independently, the effect for each RP between groups then correcting for multiple comparisons. This results in information loss because regions do not contribute to the FC matrix with the same weight (Chen et al., 2021). BML provides a more robust approach by assuming that the likelihood of an effect for every RP follows a Gaussian distribution and estimates this effect at the level of the subject and group. In other words, to properly account for **shared region dependencies** in the matrix, a **multilevel model** is formulated. The model captures the overall mean effect, random effects for individual regions, subject-level variability and residual noise. This decomposes each RP correlation that is part of the FC matrix into a global effect at each RP and the contribution of each region (Chen et al., 2021; Chen, Xiao, et al., 2019). ROI effects are a distinctive output of MBA that represent the contribution of each region within the model. Chen, Burkner et al. (2019) suggest this feature allows one to identify important regions, similar to graph outcomes that represent hubs. The final effect for each region or RP is given as a distribution, so called the posterior distribution (Chen, Bürkner, et al., 2019; Chen, Xiao, et al., 2019). The posterior distribution is interpreted by considering the strength of the area under the curve (P+), that is noting the area that is greater than the point of no effect (zero) compared to the area that is below the point of no effect. A very high P+ represents a strong probability that the effect is greater than zero, while a very low P+ means there is a weak probability the effect is greater than zero. Note, a very low P+ corresponds with a strong probability of the effect being less than zero.

To compare FC of RPs between groups and nodal measures of the functional network, we ran matrix-based analysis (MBA) and region-based analysis (RBA) programs, respectively, which use BML to compute the integrative model via Monte-Carlo Markov chains at a predetermined number of iterations. Since it has been noted that functional activation of brain areas demonstrate a sex-associated pattern that may be confounded by pubertal hormones during adolescence (Gur & Gur, 2016; Lenroot & Giedd, 2010), we included sex as a confounding variable in all our models. Our final MBA model was of the form:

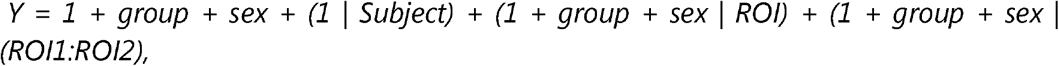

where ROI and ROI1:ROI2 represent a region and RP, respectively. Our final RBA model was of the form:

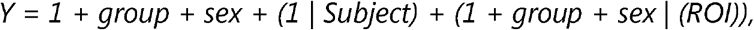

Where ROI represents a node. The resulting effect estimates were for every node.

To assess the quality of Monte-Carlo Markov chains we used the split Ř statistic which gauges the consistency of an ensemble of Markov chains. Fully converged chains correspond to Ř = 1. In the present study we used an arbitrary threshold of Ř < 1.1 for chains that converged well in both the MBA and RBA analyses.

#### Associations between structural/functional integrity outcomes and neurocognitive function

We ran Pearson correlation statistical tests between structural/functional integrity outcomes and neurocognitive measures. The structural integrity outcomes were MD, FA, AD and RD outcomes, as well the structural nodal measures degree strength and efficiency (nodal and local). The functional integrity outcomes were FC between RPs, as well as the functional nodal measures degree, strength and efficiency. We corrected for multiple comparisons using the FDR method and considered FDR q < 0.05 as our significance level.

## Results

In the DTI analysis, participants had an average of 28±3 DW volumes and 3±1 b0s and there were no significant group differences. We excluded 5 (2 CHEU and 3 CHUU) subjects due to motion artefacts present in more than half of the DWI volumes. We excluded an additional 6 (3 CHEU, 3 CHUU) participants based on having less than half of the average structural connections in the sample. The current DTI results are from a total of 37 participants (15 CHEU and 22 CHUU) (Table 1). In the RS-fMRI analysis we also excluded 11 participants (6 CHEU, 5 CHUU) due to poor image quality. The current RS-fMRI results are therefore also from a sample of 37 participants (14 CHEU, 23 CHUU) (Table 2) Notably, only two participants differed in exclusion status between the two analyses, indicating significant overlap in excluded subjects.

**Table 1:** Sample characteristics of children perinatally exposed to HIV (CHEU) and children unexposed (CHUU) (N=37) used in DTI analysis. Where a variable was not normally distributed (Shapiro-Wilk p <0.05) we summarize with median and IQR.

|  | *CHUU* | *CHEU* | *p-value<sup>a</sup>* |
| --- | --- | --- | --- |
| **Demographics** |  |  |  |
| N | 22 | 15 |  |
| Sex: Female, N (%) | 8(36) | 7(47) |  |
| Age at scan (years), Mean±SD | 11.65± 0.25 | 11.49± 0.22 | 0.08 |
| **Language-related measures** |  |  |  |
| **Sequential processing<sup>b</sup>**, Median (IQR) | 83(18.5) | 81(18.5) | 0.35 |
| Number recall<sup>c</sup>, Mean±SD | 7.41±2.54 | 7.87±2.50 | 0.09 |
| Word order<sup>c</sup>, Mean±SD | 6.42±2.42 | 6.4±1.84 | 0.31 |
| **Learning<sup>b</sup>**, Mean±SD | 91.91±13.24 | 93.13±11.94 | 0.41 |
| Atlantis<sup>c</sup>, Median (IQR) | 11(3.5) | 10(1.75) | 0.77 |
| Rebus<sup>c</sup>, Mean±SD | 6.5±3.22 | 7.53±3.32 | 0.97 |
| **Delayed recall<sup>b</sup>**, Median (IQR) | 91(10.5) | 89.5(13.5) | 0.96 |
| Rebus delayed<sup>c</sup>, Mean±SD | 6.86±2.82 | 7.47±2.64 | 0.28 |
| Atlantis delayed<sup>c</sup>, Median (IQR) | 10(2) | 10(2.75) | 0.63 |
| Story completion<sup>c</sup>, Mean±SD | 7.45±3.25 | 8.8±3.84 | 0.51 |
| Animal naming<sup>d</sup>, Mean±SD | 11.37±4.19 | 11.39±3.20 | 0.20 |
| **Nonverbal index<sup>b</sup>**, Median (IQR) | 82(23.5) | 86.5(25) | 0.59 |
| **Mental processing<sup>b</sup>**, Mean±SD | 82±13.77 | 86.87±14.45 | 0.63 |

<sup>a</sup>Student t-test, <sup>b</sup>Standardized score, <sup>c</sup>Scaled score, <sup>d</sup>Raw score

**Table 2:** Sample characteristics of children perinatally exposed to HIV (CHEU) and unexposed controls (CHUU) (N = 37) used in RS-fMRI analysis. Where a variable was not normally distributed (Shapiro-Wilk p <0.05) we summarize with median and IQR.

|  | *CHUU* | *CHEU* | *p-value<sup>a</sup>* |
| --- | --- | --- | --- |
| **Demographics** |  |  |  |
| N | 23 | 14 |  |
| Sex: Female, N (%) | 9 (39) | 7(50) |  |
| Age at scan (years), Mean±SD | 11.68± 0.43 | 11.49± 0.22 | 0.08 |
| **Language-related measures** |  |  |  |
| **Sequential processing<sup>b</sup>**,<br>Median (IQR) | 80(18.5) | 82(18.5) | **0.005** |
| Number recall<sup>c</sup>, Mean±SD | 7.41±2.54 | 7.87±2.50 | 0.08 |
| Word order<sup>c</sup>, Mean±SD | 6.42±2.42 | 6.4±1.84 | 0.20 |
| **Learning<sup>b</sup>**, Mean±SD | 88.78±11.80 | 91.92±13.02 | 0.61 |
| Atlantis<sup>c</sup>, Median (IQR) | 10(2.2) | 9.5(1.82) | 0.51 |
| Atlantis delayed<sup>c</sup>, Median (IQR) | 9.36(1.29) | 9.5(1.65) | 0.39 |
| Rebus<sup>c</sup>, Mean±SD | 5.78±2.90 | 8.22±1.20 | 0.56 |
| Rebus delayed<sup>c</sup>, Mean±SD | 6.28±2.19 | 7.33±2.67 | 0.29 |
| Story completion<sup>c</sup>, Mean±SD | 6.5±2.55 | 8±4.26 | 0.27 |
| **Delayed recall<sup>b</sup>**, Median (IQR) | 86.12(10.5) | 90.25(13.5) | 0.86 |
| Animal naming<sup>d</sup>, Mean±SD | 12.11±3.92 | 12.36±3.98 | 0.16 |
| **Nonverbal index<sup>b</sup>**, Median (IQR) | 75.81(14) | 85.5(13) | 0.37 |
| **Mental processing<sup>b</sup>**, Mean±SD | 77.16±9.81 | 84±15.43 | 0.94 |

<sup>a</sup>Student t-test, <sup>b</sup>Standardized score, <sup>c</sup>Scaled score, <sup>d</sup>Raw score

Handedness was assessed as a potential confounding variable in all analyses, but no significant effects were found.

### DTI-based tractography and structural network results

We found no significant differences in DTI measures (FA, MD, RD and AD; Figure 1) between the CHUU and CHEU groups after multiple comparison correction. The CHEU group demonstrated higher nodal strength in one auditory region, the left IC, and 3 non-auditory regions the bilateral rMFC and the right cuneus. Among unadjusted auditory findings, higher nodal strength in the right IC (p = 0.02; q = 0.11) and left MGN (p = 0.03; q = 0.15) were close to the q-value cutoff of 0.05.

**Figure 1:**
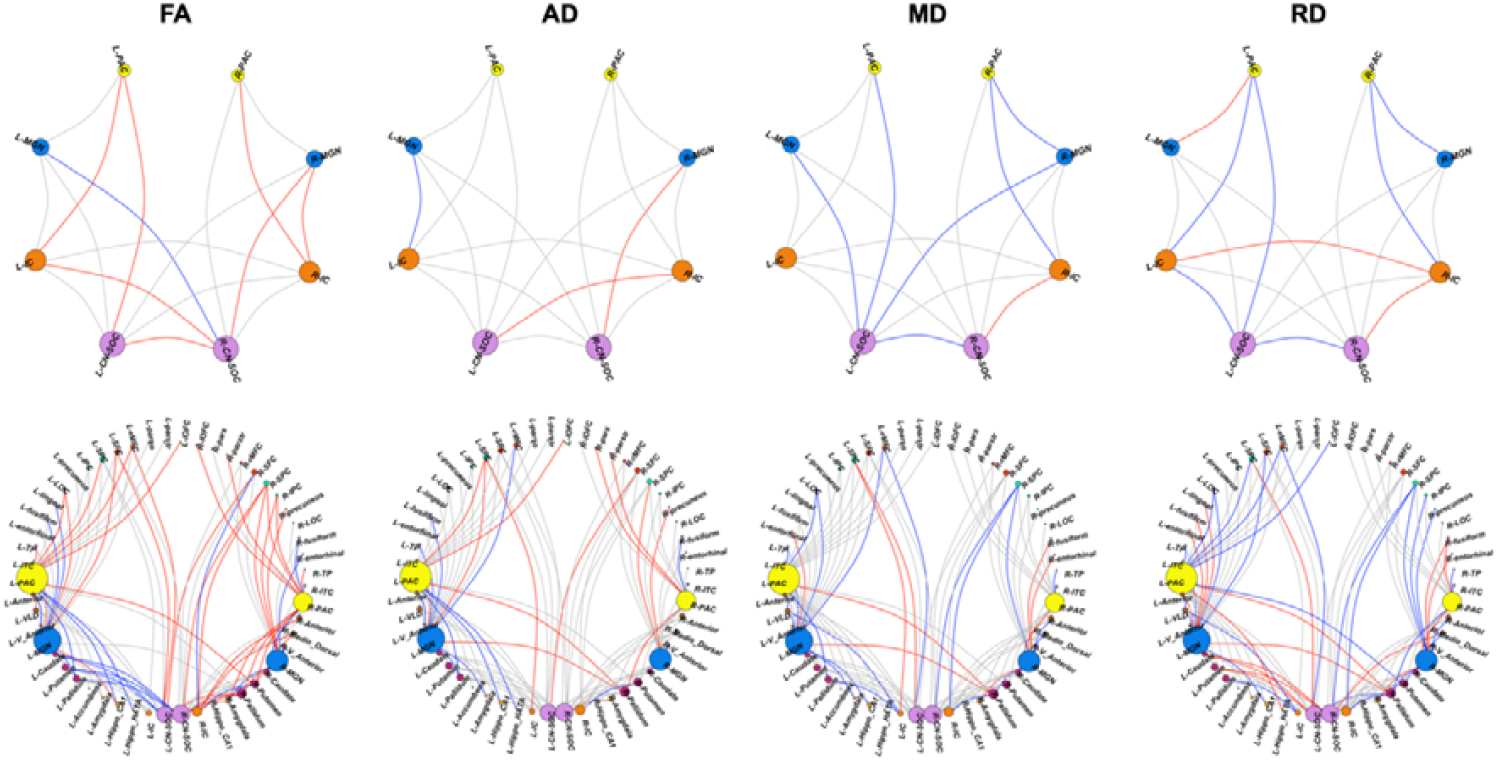
White matter connections showing higher (red) or lower (blue) FA, AD, MD or RD in CHEU compared to CHUU between auditory regions (top row) and between auditory and non-auditory regions (bottom row). Results shown are at an uncorrected p < 0.05. Connections showing no group differences at this significance level are shown in grey. ROIs are sized according to the number of links to the ROI. Colours of nodes are kept consistent across the two rows, to facilitate identification of auditory nodes in the bottom row. See Appendix A (page 19) for full description of ROI abbreviations.

*S*ummary statistics for all 127 examined WM connections across the DTI parameters between groups CHUU and CHEU can be found in the Appendices: Appendix B: DTI-based tractography. Summary statistics for all nodes between groups can be found in the Appendices section: Appendix C: DTI-based structural network.

### RS-fMRI FC and functional network results

#### Region effects

We found strong evidence (P+ >= 0.95) of lower FC in four ROIs in CHEU which included the left pericalcarine, left caudate, left hippocampus CA3 and left lingual gyrus. Four ROIs showed strong evidence of higher FC (P+ <= 0.05) in the CHEU group including right paracentral gyrus, right precentral gyrus, left postcentral gyrus and right rMFC.

#### Region Pair effects

Between the CHUU and CHEU groups, we found strong evidence of lower FC (P+ >= 0.95) between the bilateral IC in the CHEU group. Additionally, we found strong evidence of higher FC (P+ <= 0.05) in the CHEU group between the left MGN and the right precentral gyrus, left postcentral gyrus and right rMFC. The right PAC also showed strong evidence of higher FC to the right rMFC and left postcentral gyrus. Figure 2 shows the regions and RPs showing evidence of altered FC between groups. Among the results with evidence of group differences at the P+>= 0.90 level, we find reduced FC between the left hippocampus CA3 and the right IC as well as increased FC between the left MGN and right PAC. ROI and RP effect estimates for all connections to auditory regions can be found in Appendix D.

**Figure 2:**
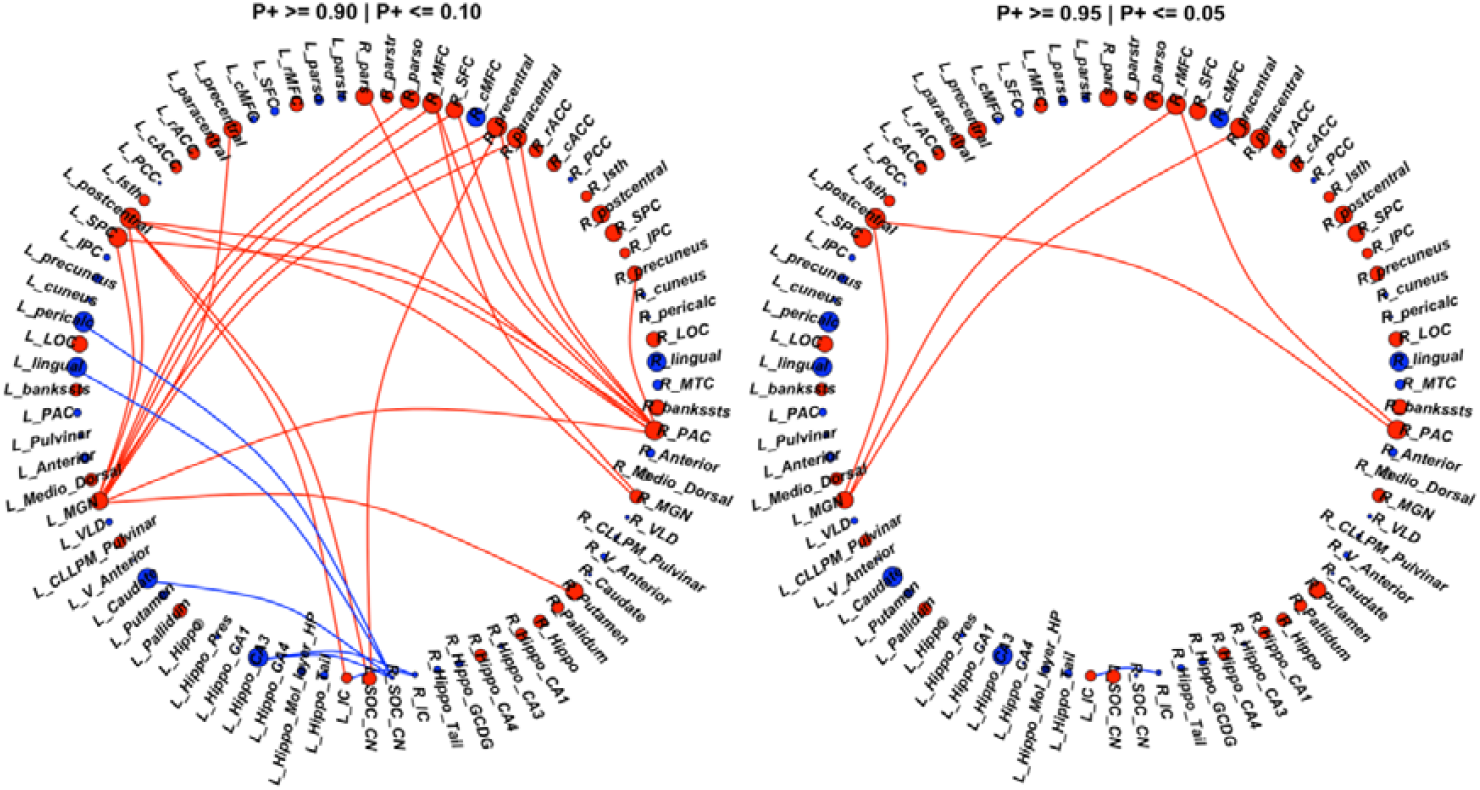
In total, 29 connections showed evidence (P+>=0.90 | P+<=0.10) of altered FC between the CHEU and CHUU groups (left). 6 connections showed lower FC (blue links) while 23 connections showed higher FC (red links) in the CHEU group compared to CHUU. Of the 29 connections, 6 showed **strong evidence (P+>=0.95** | **P+<=0.05)** (right) of altered FC (left) between groups. ROIs are colored according to direction of region effect with red and blue ROIs indicating higher and lower ROI FC in the CHEU compared to CHUU, respectively. Regions are sized according to the magnitude of ROI effect. See Appendix A (page 19) for full description of ROI abbreviations.

In the functional network analysis, we found no evidence of differences in nodal measures between CHUU and CHEU (Figure 3). Nodal effect for estimates for all nodes and graph measures can be found in Appendix E.

**Figure 3:**
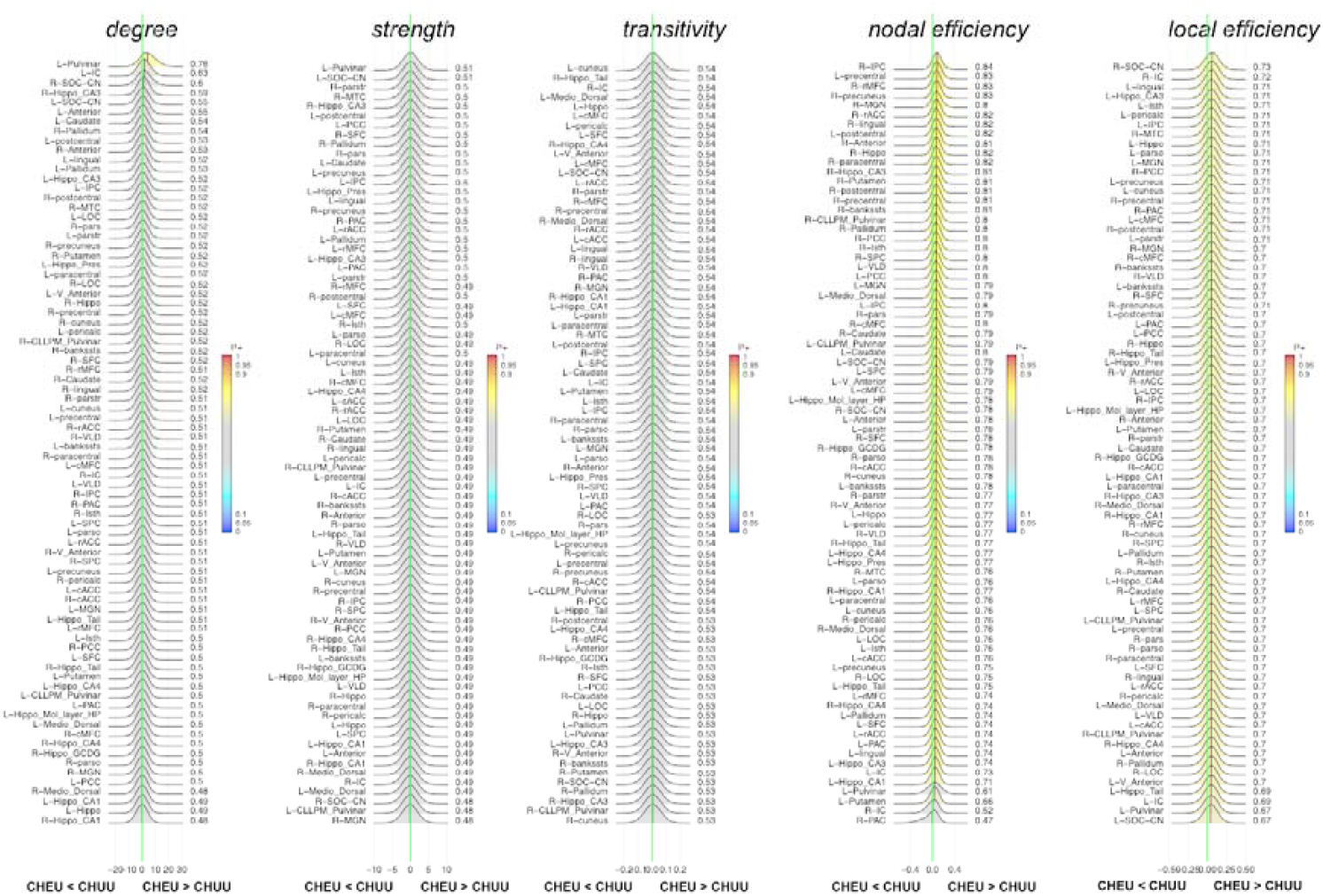
Posterior probability distribution plots for the effect magnitude of the comparison CHEU-CHUU difference between mean graph measures of functional connectivity. The graph measures include nodal measures degree, strength, transitivity, nodal and local efficiency. The green vertical line marks the location of no effect. See Appendix A (page 19) for full description of ROI abbreviations.

### Associations between structural/functional integrity outcomes and neurocognitive function

We found no associations between structural or functional integrity outcomes and neurocognitive function after correcting for multiple comparisons.

## Discussion

This study investigated the effects of exposure on WM integrity and connectivity as well as FC of the CAS and related regions in CHEU at age 11. Within the CAS, we found structural and functional connections involving the IC to be most affected by HIV exposure. In addition, we observed altered connectivity involving the bilateral MGN and R PAC to cortical regions. These findings suggest HIV exposure has subtle effects on both the connectivity of the CAS and its connectivity to other regions. However, we did not observe links between imaging and neurocognitive outcomes, suggesting the reported prenatal HIV and ART exposure related structural and functional changes do not relate to auditory and selected language capacities at age 11 in these children.

### HIV-exposure impacts the structural and functional connectivity of the IC within the CAS

In the midbrain, the IC is a communication center for ascending and descending auditory information (Suga, 2008; Xiong et al., 2015). It integrates data from the cortex as well information from the contralateral IC, and its primary target is the MGN through which it sends information to the cortex.

We reported lower FC between the bilateral IC. Interhemispheric coherence is crucial for efficient sound processing and the IC is especially crucial for sound localization (Grothe et al., 2010; Xiong et al., 2015). The left IC had higher structural nodal strength in CHEU compared to CHUU. Unadjusted results show the right IC with higher nodal strength among CHEU, suggesting a possible bilateral effect. Nodal strength is related to the ability of a region of interest to effectively communicate, and typically increases with age in childhood (Dennis et al., 2013; Hagmann et al., 2010; Wu et al., 2013). Strength is a measure of connectivity, and nodes with higher strength act as communication hubs. Increased nodal strength suggests stronger structural connectivity, with the IC performing more efficiently and/or compensating for disruptions within the system.

HEU children are at risk of neurodevelopmental disorders including autism spectrum disorder (ASD) (Budd et al., 2018; Piske et al., 2018). Alterations in the IC have been noted in relation to ASD, which includes hyper- or hypo-reactivity to sensory input (Hirota & King, 2023; Schelinski et al., 2022). In addition, the IC has been linked to tinnitus (Berger & Coomber, 2015) which is found to be more common in ASD populations (Danesh et al., 2015). Increased connectivity in the IC has also been related to exposure to environmental noise in children (Martínez-Vilavella et al., 2023). Martinez-Vilavella et al. reported higher FC between the IC and the MGN to be associated with higher exposure to annual average road traffic noise. An HIV exposure animal model looking at the gestational and breastfeeding period found ART disruptions to the developing auditory system (DeBacker et al., 2022). The auditory disruptions were influenced by specific ART regimens as well as noise and antibiotic exposure (DeBacker et al., 2022).

We did not have access to early drug/environmental exposures or neurodiversity measures and therefore cannot make direct links to observed changes and ASD, tinnitus or noise exposure effects. Future work examining the CAS in this population would benefit from the inclusion of these factors to further explore possible links with the IC.

### HIV-exposure affects connections between the MGN/PAC and other brain regions

The auditory system is involved with other parts of the brain, such as the prefrontal cortex and limbic system, that work together to understand and process auditory information (Suga, 2008). Our findings show altered functional connectivity to these areas, including the precentral gyrus, hippocampus and caudate. While most alterations reported represented increased FC in relation to HIV exposure, the region effects of the left hippocampus CA3 and region pair effects of the left hippocampus CA3 and right IC demonstrated reduced FC. While the IC is an important hub within the CAS, it also plays a key role in linking the auditory brainstem to sensory, motor, and limbic systems for auditory processing (Winer & Schreiner, 2005). The hippocampus is involved in the temporal processing of sensory information, with animal research linking hippocampal CA3 neurons to sensory gating (Bickford-Wimer et al., 1990). The reduction in FC to the left hippocampus CA3 may be related to the reported IC alterations, suggesting these changes may influence the auditory system as well as those that work with it.

Within the CAS, we observed increased FC between the L MGN and the R PAC. Uncorrected DTI results show higher nodal strength in the L MGN. Regions with higher strength act as communication centers between specialized parts of the brain. Both the MGN and PAC play important roles in integrating auditory information within the CAS but also communicate with other systems of the brain.

Both the MGN and PAC showed higher FC to the right rMFC in the CHEU group compared to CHUU. The bilateral rMFC showed higher nodal strength in the same group. The rMFC interacts with the CAS for efficient processing of literacy and numeracy tasks (Koyama et al., 2017). At age 11, children are undergoing substantial development in numeracy and literacy skills (Demetriou et al., 2014; Kail & Ferrer, 2007). Given the lack of HIV exposure differences in cognitive measures, and the increases in connectivity outcomes, our results may indicate overcompensation of the CAS with these processes. Alternatively, since we did not find any associations supporting increased connectivity or strength with cognitive outcomes, our results may reflect differences in brain development due to HIV exposure that do not lead to functional disruptions at this age.

## Limitations

A few limitations in our study of observed exposure alterations are worth noting. The lack of hearing data at the time of neuroimaging curbs the interpretations that can be drawn between imaging changes and auditory function. While we present neurocognitive scores, which rely on hearing abilities, they do not capture the full spectrum of auditory capacity.

The second limitation is the small sample size. We’ve included both corrected and uncorrected results in order to identify possible patterns related to HIV exposure for hypothesis generation. Future studies with larger populations of CHEU are needed. In addition, the inclusion of potential confounders related to HIV exposure, such as duration of ART exposure in utero and in infancy, are needed.

Finally there is a methodological difference in how the structural and functional networks were defined, which may impact the comparability between the two modalities. Structural connectivity was derived using probabilistic tractography to estimate the likelihood of white matter tracts between pairs of regions. To enhance the specificity and reduce false positives, we included only those tracts that were consistently estimated across all subjects. As a result, the final structural connectivity matrices contained fewer ROIs than originally defined in the atlas. In contrast, the functional connectivity networks retained all 80 ROIs, as functional connections were based on Pearson correlations of regional time series, which do not rely on anatomical tract presence. While this conservative approach to defining structural connections was intended to improve the reliability of the structural networks, it introduced a discrepancy in ROI coverage that may limit direct comparisons between structural and functional connectivity patterns.

## Future work

Given the age of the cohort, and the maturation the CAS undergoes in adolescence, it is possible that our results of altered FC and structural network outcomes may affect neurocognitive domains later in adolescence. Further cross-sectional analysis later in adolescence is needed to understand the potential consequences of these findings.

Longitudinal analysis of the CAS, including previous as well as later time points, could provide a better understanding of the nature of functional and structural HIV-exposure alterations. Considering the CHEU had reduced Sequential Processing measures at ages 7 and 9 years (van Wyhe et al., 2021), and then increased or similar outcomes to the CHUU group at age 11 within this subgroup, it would be particularly interesting to study the development of the IC in relation to auditory outcomes starting at age 5. In addition, future analysis could explore negative functional connectivity as well as associations between DTI and RS-fMRI to better understand the CAS structure-function relationship.

The inclusion of relevant complementary data, such as ART exposure, environmental noise, neurodiversity testing, hearing, immunological markers and congenital cytomegalovirus infection, may also provide a more complete picture of the alterations reported in this cohort.

## Conclusion

This is the first study examining the functional and structural connectivity of the CAS in HEU children. Within the CAS, we report functional and structural alterations related to the IC. In relation to the CAS and other brain regions, we find connectivity changes between the cortex and both the L MGN and R PAC. These findings suggest developmental differences in connectivity in HEU children within the CAS that do not relate to neurocognitive abilities .

Considering the small body of pediatric imaging literature in children exposed to HIV and HIV free, as well as work focused on the CAS, the work presented is an important contribution to an understudied population and health concern. Therefore, even with the limitations of this study, the data generated can motivate and inform future work in this field.

## Funding information

This work was supported by the National Institute On Deafness And Other Communication Disorders of the National Institutes of Health under Award Number R01DC015984; South African Medical Research Council (SAMRC); South African National Research Foundation (NRF) grants CPR20110614000019421 and CPRR150723129691; and the NRF/DST South African Research Chairs Initiative; Support for the CHER study, which provided the infrastructure for the neurodevelopmental study, was provided by the US National Institute of Allergy and Infectious Diseases through the CIPRA network, Grant U19 AI53217; the Departments of Health of the Western Cape and Gauteng, South Africa; and GlaxoSmithKline/Viiv Healthcare. Additional support was provided with Federal funds from the National Institute of Allergy and Infectious Diseases, National Institutes of Health, United States Department of Health and Human Services, under Contract No. HHSN272200800014C.

## Appendices

## Appendix A: Automatically Segmented Regions

**Table 3:** 126 automatically segmented ROIs. R – right, L – left.

| ROI | ROI abbreviation |
| --- | --- |
| R-lateralorbitofrontal | R-IOFC |
| R-parsorbitalis | R-pars |
| R-frontalpole | R-FP |
| R-medialorbitofrontal | R-mOFC |
| R-parstriangularis | R-parstr |
| R-parsopercularis | R-parso |
| R-rostralmiddlefrontal | R-rMFC |
| R-superiorfrontal | R-SFC |
| R-caudalmiddlefrontal | R-cMFC |
| R-precentral | R-precentral |
| R-paracentral | R-paracentral |
| R-rostralanteriorcingulate | R-rACC |
| R-caudalanteriorcingulate | R-cACC |
| R-posteriorcingulate | R-PCC |
| R-isthmuscingulate | R-Isth |
| R-postcentral | R-postcentral |
| R-supramarginal | R-supramarginal |
| R-superiorparietal | R-SPC |
| R-inferiorparietal | R-IPC |
| R-precuneus | R-precuneus |
| R-cuneus | R-cuneus |
| R-pericalcarine | R-pericalc |
| R-lateraloccipital | R-LOC |
| R-lingual | R-lingual |
| R-fusiform | R-fusiform |
| R-parahippocampal | R-paraHippo |
| R-entorhinal | R-entorhinal |
| R-temporalpole | R-TP |
| R-inferiortemporal | R-ITC |
| R-middletemporal | R-MTC |
| R-bankssts | R-bankssts |
| R-superiortemporal | R-STC |
| R-transversetemporal | R-TTC |
| R-insula | R-insula |
| R-Pulvinar | R-Pulvinar |
| R-Anterior | R-Anterior |
| R-Medio_Dorsal | R-Medio_Dorsal |
| R-Ventral_Latero_Dorsal | R-VLD |
| R-Central_Lateral-Lateral_Posterior-Medial_Pulvinar | R-CLLPM_Pulvinar |
| R-Ventral_Anterior | R-Ventral_Anterior |
| R-Ventral_Latero_Ventral | R-VLV |
| R-Caudate | R-Caudate |
| R-Putamen | R-Putamen |
| R-Pallidum | R-Pallidum |
| R-Accumbens_area | R-Accumbens |
| R-Amygdala | R-Amygdala |
| R-Hippocampus | R-Hippo |
| R-Hippocampus_Parasubiculum | R-Hippo_Para |
| R-Hippocampus_Presubiculum | R-Hippo_Pres |
| R-Hippocampus_Subiculum | R-Hippo_Sub |
| R-Hippocampus_CA1 | R-Hippo_CA1 |
| R-Hippocampus_CA3 | R-Hippo_CA3 |
| R-Hippocampus_CA4 | R-Hippo_CA4 |
| R-Hippocampus_GCDG | R-Hippo_GCDG |
| R-Hippocampus_HATA | R-Hippo_HATA |
| R-Hippocampus_Fimbria | R-Hippo_Fimb |
| R-Hippocampus_Molecular_layer_HP | R-Hippo_Mol_layer_HP |
| R-Hippocampus_Hippocampal_fissure | R-Hippo_fissure |
| R-Hippocampus_Tail | R-Hippo_Tail |
| R-VentralDC | R-VDC |
| R-Hypothalamus | R-Hypothalamus |
| L-lateralorbitofrontal | L-IOFC |
| L-parsorbitalis | L-pars |
| L-frontalpole | L-FP |
| L-medialorbitofrontal | L-mOFC |
| L-parstriangularis | L-parstr |
| L-parsopercularis | L-parso |
| L-rostralmiddlefrontal | L-rMFC |
| L-superiorfrontal | L-SFC |
| L-caudalmiddlefrontal | L-cMFC |
| L-precentral | L-precentral |
| L-paracentral | L-paracentral |
| L-rostralanteriorcingulate | L-rACC |
| L-caudalanteriorcingulate | L-cACC |
| L-posteriorcingulate | L-PCC |
| L-isthmuscingulate | L-Isth |
| L-postcentral | L-postcentral |
| L-supramarginal | L-supramarginal |
| L-superiorparietal | L-SPC |
| L-inferiorparietal | L-IPC |
| L-precuneus | L-precuneus |
| L-cuneus | L-cuneus |
| L-pericalcarine | L-pericalc |
| L-lateraloccipital | L-LOC |
| L-lingual | L-lingual |
| L-fusiform | L-fusiform |
| L-parahippocampal | L-paraHippo |
| L-entorhinal | L-entorhinal |
| L-temporalpole | L-TP |
| L-inferiortemporal | L-ITC |
| L-middletemporal | L-MTC |
| L-bankssts | L-bankssts |
| L-superiortemporal | L-STC |
| L-transversetemporal | L-TTC |
| L-insula | L-insula |
| L-Pulvinar | L-Pulvinar |
| L-Anterior | L-Anterior |
| L-Medio_Dorsal | L-Medio_Dorsal |
| L-Ventral_Latero_Dorsal | L-VLD |
| L-Central_Lateral-Lateral_Posterior-Medial_Pulvinar | L-CLLPM_Pulvinar |
| L-Ventral_Anterior | L-Ventral_Anterior |
| L-Ventral_Latero_Ventral | L-VLV |
| L-Caudate | L-Caudate |
| L-Putamen | L-Putamen |
| L-Pallidum | L-Pallidum |
| L-Accumbens_area | L-Accumbens |
| L-Amygdala | L-Amygdala |
| L-Hippocampus | L-Hippo |
| L-Hippocampus_Parasubiculum | L-Hippo_Para |
| L-Hippocampus_Presubiculum | L-Hippo_Pres |
| L-Hippocampus_Subiculum | L-Hippo_Sub |
| L-Hippocampus_CA1 | L-Hippo_CA1 |
| L-Hippocampus_CA3 | L-Hippo_CA3 |
| L-Hippocampus_CA4 | L-Hippo_CA4 |
| L-Hippocampus_GCDG | L-Hippo_GCDG |
| L-Hippocampus_HATA | L-Hippo_HATA |
| L-Hippocampus_Fimbria | L-Hippo_Fimb |
| L-Hippocampus_Molecular_layer_HP | L-Hippo_Mol_layer_HP |
| L-Hippocampus_Hippocampal_fissure | L-Hippo_fissure |
| L-Hippocampus_Tail | L-Hippo_Tail |
| L-VentralDC | L-VDC |
| L-Hypothalamus | L-Hypothalamus |
| Brain_Stem-Midbrain | BS-mid |
| Brain_Stem-Pons | BS-pons |
| Brain_Stem-Medulla | BS-medulla |
| Brain_Stem-SCP | BS-SCP |

## Appendix B: DTI-based tractography

**Table 4:** Standardized Beta (Std β), Standard Error (Std SE) values, p values and FDR corrected q values for the comparison between CHUU and CHEU. Each row represents the relevant statistics for a WM connection and DTI measure thus each WM connection is represented in 6 rows for 6 DTI measures. L – left, R – right.

| ROI1 | ROI2 | DTI measure | Std $\beta$ | Std SE | p-value | FDR q value |
| --- | --- | --- | --- | --- | --- | --- |
| L IC | L MGN | AD | 0.1737 | 0.1737 | 0.3249 | 0.9951 |
| L IC | L MGN | FA | 0.0654 | 0.1749 | 0.7110 | 0.9395 |
| L IC | L MGN | fNT | 0.2047 | 0.1738 | 0.2482 | 0.6924 |
| L IC | L MGN | MD | 0.0133 | 0.1815 | 0.9420 | 0.9987 |
| L IC | L MGN | PV | 0.2161 | 0.1750 | 0.2262 | 0.9039 |
| L IC | L MGN | RD | 0.1053 | 0.1731 | 0.5469 | 0.9964 |
| L IC | L CN/SOC | AD | 0.0222 | 0.1194 | 0.8533 | 0.9951 |
| L IC | L CN/SOC | FA | 0.0117 | 0.1170 | 0.9206 | 0.9895 |
| L IC | L CN/SOC | fNT | 0.0594 | 0.1089 | 0.5874 | 0.8723 |
| L IC | L CN/SOC | MD | 0.0425 | 0.1191 | 0.7222 | 0.9987 |
| L IC | L CN/SOC | PV | 0.0920 | 0.1188 | 0.4410 | 0.9083 |
| L IC | L CN/SOC | RD | 0.0904 | 0.1165 | 0.4405 | 0.9964 |
| L MGN | L CN/SOC | AD | 0.0798 | 0.1183 | 0.5021 | 0.9951 |
| L MGN | L CN/SOC | FA | 0.0616 | 0.1173 | 0.6010 | 0.9215 |
| L MGN | L CN/SOC | fNT | 0.0316 | 0.1225 | 0.7970 | 0.9411 |
| L MGN | L CN/SOC | MD | 0.0917 | 0.1196 | 0.4458 | 0.9987 |
| L MGN | L CN/SOC | PV | 0.1215 | 0.1194 | 0.3123 | 0.9039 |
| L MGN | L CN/SOC | RD | 0.0032 | 0.1187 | 0.9785 | 0.9964 |
| L PAC | L IC | AD | 0.0511 | 0.1746 | 0.7719 | 0.9951 |
| L PAC | L IC | FA | 0.1888 | 0.1614 | 0.2507 | 0.9215 |
| L PAC | L IC | fNT | 0.2770 | 0.1560 | 0.0852 | 0.4089 |
| L PAC | L IC | MD | 0.0161 | 0.1782 | 0.9286 | 0.9987 |
| L PAC | L IC | PV | 0.2823 | 0.1683 | 0.1035 | 0.7850 |
| L PAC | L IC | RD | 0.1346 | 0.1724 | 0.4405 | 0.9964 |
| L PAC | L MGN | AD | 0.0220 | 0.1736 | 0.9001 | 0.9951 |
| L PAC | L MGN | FA | 0.1072 | 0.1733 | 0.5405 | 0.9215 |
| L PAC | L MGN | fNT | 0.0189 | 0.1770 | 0.9155 | 0.9826 |
| L PAC | L MGN | MD | 0.0830 | 0.1728 | 0.6343 | 0.9987 |
| L PAC | L MGN | PV | 0.1306 | 0.1614 | 0.4246 | 0.9083 |
| L PAC | L MGN | RD | 0.1370 | 0.1716 | 0.4305 | 0.9964 |
| L PAC | L CN/SOC | AD | 0.0196 | 0.1164 | 0.8665 | 0.9951 |
| L PAC | L CN/SOC | FA | 0.2715 | 0.1029 | 0.0103 | 0.4498 |
| L PAC | L CN/SOC | fNT | 0.2511 | 0.1032 | 0.0176 | 0.2073 |
| L PAC | L CN/SOC | MD | 0.2700 | 0.1169 | 0.0240 | 0.9987 |
| L PAC | L CN/SOC | PV | 0.1844 | 0.1197 | 0.1283 | 0.8585 |
| L PAC | L CN/SOC | RD | 0.2927 | 0.1132 | 0.0118 | 0.9678 |
| R CN/SOC | L IC | AD | 0.0398 | 0.1824 | 0.8287 | 0.9951 |
| R CN/SOC | L IC | FA | 0.1371 | 0.1723 | 0.4320 | 0.9215 |
| R CN/SOC | L IC | fNT | 0.1636 | 0.1687 | 0.3393 | 0.7610 |
| R CN/SOC | L IC | MD | 0.0003 | 0.1718 | 0.9987 | 0.9987 |
| R CN/SOC | L IC | PV | 0.3893 | 0.1540 | 0.0166 | 0.4604 |
| R CN/SOC | L IC | RD | 0.0252 | 0.1722 | 0.8846 | 0.9964 |
| R CN/SOC | L MGN | AD | 0.0226 | 0.1740 | 0.8973 | 0.9951 |
| R CN/SOC | L MGN | FA | 0.1881 | 0.1727 | 0.2847 | 0.9215 |
| R CN/SOC | L MGN | fNT | 0.1052 | 0.1730 | 0.5476 | 0.8674 |
| R CN/SOC | L MGN | MD | 0.0214 | 0.1760 | 0.9038 | 0.9987 |
| R CN/SOC | L MGN | PV | 0.0402 | 0.1754 | 0.8201 | 0.9779 |
| R CN/SOC | L MGN | RD | 0.1043 | 0.1711 | 0.5464 | 0.9964 |
| R CN/SOC | L CN/SOC | AD | 0.0303 | 0.1148 | 0.7928 | 0.9951 |
| R CN/SOC | L CN/SOC | FA | 0.3312 | 0.1112 | 0.0040 | 0.4429 |
| R CN/SOC | L CN/SOC | fNT | 0.1005 | 0.1191 | 0.4014 | 0.7875 |
| R CN/SOC | L CN/SOC | MD | 0.1163 | 0.1126 | 0.3052 | 0.9987 |
| R CN/SOC | L CN/SOC | PV | 0.0381 | 0.1207 | 0.7533 | 0.9629 |
| R CN/SOC | L CN/SOC | RD | 0.2589 | 0.1137 | 0.0260 | 0.9964 |
| R IC | L IC | AD | 0.0439 | 0.1741 | 0.8026 | 0.9951 |
| R IC | L IC | FA | 0.0513 | 0.1704 | 0.7652 | 0.9557 |
| R IC | L IC | fNT | 0.0043 | 0.1781 | 0.9808 | 0.9899 |
| R IC | L IC | MD | 0.0368 | 0.1739 | 0.8338 | 0.9987 |
| R IC | L IC | PV | 0.0378 | 0.1799 | 0.8351 | 0.9799 |
| R IC | L IC | RD | 0.1674 | 0.1782 | 0.3552 | 0.9964 |
| R IC | L CN/SOC | AD | 0.1272 | 0.1721 | 0.4651 | 0.9951 |
| R IC | L CN/SOC | FA | 0.0673 | 0.1649 | 0.6857 | 0.9395 |
| R IC | L CN/SOC | fNT | 0.3665 | 0.1542 | 0.0236 | 0.2400 |
| R IC | L CN/SOC | MD | 0.1093 | 0.1730 | 0.5318 | 0.9987 |
| R IC | L CN/SOC | PV | 0.1136 | 0.1873 | 0.5490 | 0.9245 |
| R IC | L CN/SOC | RD | 0.0068 | 0.1702 | 0.9682 | 0.9964 |
| R IC | R MGN | AD | 0.0490 | 0.1741 | 0.7800 | 0.9951 |
| R IC | R MGN | FA | 0.4933 | 0.1536 | 0.0032 | 0.4429 |
| R IC | R MGN | fNT | 0.4461 | 0.1786 | 0.0194 | 0.2150 |
| R IC | R MGN | MD | 0.0011 | 0.1728 | 0.9947 | 0.9987 |
| R IC | R MGN | PV | 0.3634 | 0.1569 | 0.0271 | 0.4814 |
| R IC | R MGN | RD | 0.0461 | 0.1695 | 0.7874 | 0.9964 |
| R IC | R CN/SOC | AD | 0.1089 | 0.1713 | 0.5293 | 0.9951 |
| R IC | R CN/SOC | FA | 0.0164 | 0.1695 | 0.9235 | 0.9895 |
| R IC | R CN/SOC | fNT | 0.3125 | 0.1704 | 0.0763 | 0.4089 |
| R IC | R CN/SOC | MD | 0.1306 | 0.1725 | 0.4543 | 0.9987 |
| R IC | R CN/SOC | PV | 0.3028 | 0.1741 | 0.0923 | 0.7648 |
| R IC | R CN/SOC | RD | 0.1279 | 0.1728 | 0.4642 | 0.9964 |
| R MGN | L CN/SOC | AD | 0.0421 | 0.1766 | 0.8130 | 0.9951 |
| R MGN | L CN/SOC | FA | 0.0294 | 0.1676 | 0.8618 | 0.9772 |
| R MGN | L CN/SOC | fNT | 0.0922 | 0.1634 | 0.5769 | 0.8723 |
| R MGN | L CN/SOC | MD | 0.2055 | 0.1739 | 0.2467 | 0.9987 |
| R MGN | L CN/SOC | PV | 0.2285 | 0.1780 | 0.2091 | 0.9039 |
| R MGN | L CN/SOC | RD | 0.0517 | 0.1734 | 0.7675 | 0.9964 |
| R MGN | R CN/SOC | AD | 0.1865 | 0.1626 | 0.2599 | 0.9951 |
| R MGN | R CN/SOC | FA | 0.2064 | 0.1658 | 0.2222 | 0.9215 |
| R MGN | R CN/SOC | fNT | 0.0421 | 0.1767 | 0.8135 | 0.9411 |
| R MGN | R CN/SOC | MD | 0.1057 | 0.1674 | 0.5320 | 0.9987 |
| R MGN | R CN/SOC | PV | 0.2513 | 0.1738 | 0.1582 | 0.9039 |
| R MGN | R CN/SOC | RD | 0.0817 | 0.1733 | 0.6404 | 0.9964 |
| R PAC | R IC | AD | 0.1026 | 0.1815 | 0.5762 | 0.9951 |
| R PAC | R IC | FA | 0.2510 | 0.1829 | 0.1811 | 0.9215 |
| R PAC | R IC | fNT | 0.2061 | 0.1645 | 0.2196 | 0.6924 |
| R PAC | R IC | MD | 0.1238 | 0.1730 | 0.4798 | 0.9987 |
| R PAC | R IC | PV | 0.0307 | 0.1730 | 0.8603 | 0.9844 |
| R PAC | R IC | RD | 0.1621 | 0.1729 | 0.3556 | 0.9964 |
| R PAC | R MGN | AD | 0.0502 | 0.1693 | 0.7688 | 0.9951 |
| R PAC | R MGN | FA | 0.0794 | 0.1872 | 0.6748 | 0.9395 |
| R PAC | R MGN | fNT | 0.1037 | 0.1788 | 0.5664 | 0.8723 |
| R PAC | R MGN | MD | 0.1196 | 0.1723 | 0.4926 | 0.9987 |
| R PAC | R MGN | PV | 0.1117 | 0.1733 | 0.5240 | 0.9164 |
| R PAC | R MGN | RD | 0.1605 | 0.1718 | 0.3572 | 0.9964 |
| R PAC | R CN/SOC | AD | 0.0489 | 0.1738 | 0.7801 | 0.9951 |
| R PAC | R CN/SOC | FA | 0.0644 | 0.1826 | 0.7269 | 0.9395 |
| R PAC | R CN/SOC | fNT | 0.2724 | 0.1493 | 0.0771 | 0.4089 |
| R PAC | R CN/SOC | MD | 0.0317 | 0.1726 | 0.8554 | 0.9987 |
| R PAC | R CN/SOC | PV | 0.1255 | 0.1795 | 0.4899 | 0.9103 |
| R PAC | R CN/SOC | RD | 0.0471 | 0.1715 | 0.7855 | 0.9964 |
| L Pallidum | L IC | AD | 0.0274 | 0.1759 | 0.8771 | 0.9951 |
| L Pallidum | L IC | FA | 0.0980 | 0.1754 | 0.5800 | 0.9215 |
| L Pallidum | L IC | fNT | 0.4254 | 0.1635 | 0.0147 | 0.1973 |
| L Pallidum | L IC | MD | 0.1606 | 0.1743 | 0.3637 | 0.9987 |
| L Pallidum | L IC | PV | 0.1923 | 0.1665 | 0.2564 | 0.9039 |
| L Pallidum | L IC | RD | 0.2177 | 0.1727 | 0.2167 | 0.9964 |
| L Putamen | L IC | AD | 0.0437 | 0.1677 | 0.7961 | 0.9951 |
| L Putamen | L IC | FA | 0.0163 | 0.1697 | 0.9239 | 0.9895 |
| L Putamen | L IC | fNT | 0.4427 | 0.1624 | 0.0106 | 0.1817 |
| L Putamen | L IC | MD | 0.0809 | 0.1704 | 0.6383 | 0.9987 |
| L Putamen | L IC | PV | 0.2487 | 0.1684 | 0.1492 | 0.9039 |
| L Putamen | L IC | RD | 0.0832 | 0.1731 | 0.6339 | 0.9964 |
| L SPC | L IC | AD | 0.1445 | 0.1597 | 0.3723 | 0.9951 |
| L SPC | L IC | FA | 0.0638 | 0.1734 | 0.7153 | 0.9395 |
| L SPC | L IC | fNT | 0.3486 | 0.1623 | 0.0392 | 0.3018 |
| L SPC | L IC | MD | 0.1140 | 0.1580 | 0.4758 | 0.9987 |
| L SPC | L IC | PV | 0.2442 | 0.1687 | 0.1572 | 0.9039 |
| L SPC | L IC | RD | 0.0659 | 0.1626 | 0.6878 | 0.9964 |
| L VLD | L IC | AD | 0.0896 | 0.1699 | 0.6016 | 0.9951 |
| L VLD | L IC | FA | 0.0219 | 0.1639 | 0.8945 | 0.9883 |
| L VLD | L IC | fNT | 0.4207 | 0.1669 | 0.0175 | 0.2073 |
| L VLD | L IC | MD | 0.1527 | 0.1748 | 0.3889 | 0.9987 |
| L VLD | L IC | PV | 0.1101 | 0.1710 | 0.5243 | 0.9164 |
| L VLD | L IC | RD | 0.1523 | 0.1683 | 0.3721 | 0.9964 |
| L Amygdala | L MGN | AD | 0.1903 | 0.1696 | 0.2698 | 0.9951 |
| L Amygdala | L MGN | FA | 0.0682 | 0.1705 | 0.6918 | 0.9395 |
| L Amygdala | L MGN | fNT | 0.1102 | 0.1723 | 0.5267 | 0.8622 |
| L Amygdala | L MGN | MD | 0.1474 | 0.1665 | 0.3825 | 0.9987 |
| L Amygdala | L MGN | PV | 0.1105 | 0.1722 | 0.5254 | 0.9164 |
| L Amygdala | L MGN | RD | 0.0581 | 0.1675 | 0.7307 | 0.9964 |
| L Anterior | L MGN | AD | 0.0166 | 0.1731 | 0.9240 | 0.9951 |
| L Anterior | L MGN | FA | 0.0880 | 0.1731 | 0.6145 | 0.9260 |
| L Anterior | L MGN | fNT | 0.0205 | 0.1696 | 0.9046 | 0.9789 |
| L Anterior | L MGN | MD | 0.0949 | 0.1727 | 0.5866 | 0.9987 |
| L Anterior | L MGN | PV | 0.1652 | 0.1573 | 0.3012 | 0.9039 |
| L Anterior | L MGN | RD | 0.1308 | 0.1726 | 0.4538 | 0.9964 |
| L Caudate | L MGN | AD | 0.0361 | 0.1683 | 0.8314 | 0.9951 |
| L Caudate | L MGN | FA | 0.0824 | 0.1666 | 0.6241 | 0.9260 |
| L Caudate | L MGN | fNT | 0.1453 | 0.1648 | 0.3842 | 0.7832 |
| L Caudate | L MGN | MD | 0.0861 | 0.1702 | 0.6164 | 0.9987 |
| L Caudate | L MGN | PV | 0.0299 | 0.1689 | 0.8607 | 0.9844 |
| L Caudate | L MGN | RD | 0.1173 | 0.1708 | 0.4970 | 0.9964 |
| L entorhinal | L MGN | AD | 0.0997 | 0.1719 | 0.5661 | 0.9951 |
| L entorhinal | L MGN | FA | 0.0234 | 0.1769 | 0.8955 | 0.9883 |
| L entorhinal | L MGN | fNT | 0.0113 | 0.1741 | 0.9487 | 0.9896 |
| L entorhinal | L MGN | MD | 0.1539 | 0.1632 | 0.3528 | 0.9987 |
| L entorhinal | L MGN | PV | 0.3898 | 0.1671 | 0.0268 | 0.4814 |
| L entorhinal | L MGN | RD | 0.1259 | 0.1674 | 0.4574 | 0.9964 |
| L fusiform | L MGN | AD | 0.1120 | 0.1551 | 0.4752 | 0.9951 |
| L fusiform | L MGN | FA | 0.1754 | 0.1714 | 0.3135 | 0.9215 |
| L fusiform | L MGN | fNT | 0.0291 | 0.1731 | 0.8675 | 0.9643 |
| L fusiform | L MGN | MD | 0.0309 | 0.1520 | 0.8402 | 0.9987 |
| L fusiform | L MGN | PV | 0.0797 | 0.1727 | 0.6476 | 0.9626 |
| L fusiform | L MGN | RD | 0.1369 | 0.1641 | 0.4101 | 0.9964 |
| L Hippo_CA1 | L MGN | AD | 0.0982 | 0.1726 | 0.5733 | 0.9951 |
| L Hippo_CA1 | L MGN | FA | 0.2556 | 0.1606 | 0.1210 | 0.8145 |
| L Hippo_CA1 | L MGN | fNT | 0.0307 | 0.1717 | 0.8592 | 0.9643 |
| L Hippo_CA1 | L MGN | MD | 0.2822 | 0.1623 | 0.0914 | 0.9987 |
| L Hippo_CA1 | L MGN | PV | 0.0500 | 0.1716 | 0.7726 | 0.9629 |
| L Hippo_CA1 | L MGN | RD | 0.2895 | 0.1591 | 0.0780 | 0.9964 |
| L IPC | L MGN | AD | 0.0185 | 0.1488 | 0.9020 | 0.9951 |
| L IPC | L MGN | FA | 0.0618 | 0.1766 | 0.7284 | 0.9395 |
| L IPC | L MGN | fNT | 0.1833 | 0.1735 | 0.2987 | 0.7291 |
| L IPC | L MGN | MD | 0.0041 | 0.1553 | 0.9793 | 0.9987 |
| L IPC | L MGN | PV | 0.0845 | 0.1717 | 0.6257 | 0.9566 |
| L IPC | L MGN | RD | 0.0199 | 0.1661 | 0.9055 | 0.9964 |
| L ITC | L MGN | AD | 0.1538 | 0.1529 | 0.3218 | 0.9951 |
| L ITC | L MGN | FA | 0.3239 | 0.1640 | 0.0568 | 0.6458 |
| L ITC | L MGN | fNT | 0.0330 | 0.1725 | 0.8493 | 0.9626 |
| L ITC | L MGN | MD | 0.1145 | 0.1506 | 0.4526 | 0.9987 |
| L ITC | L MGN | PV | 0.0956 | 0.1698 | 0.5773 | 0.9457 |
| L ITC | L MGN | RD | 0.4978 | 0.1577 | 0.0037 | 0.7924 |
| L lingual | L MGN | AD | 0.1040 | 0.1705 | 0.5458 | 0.9951 |
| L lingual | L MGN | FA | 0.2435 | 0.1690 | 0.1591 | 0.9215 |
| L lingual | L MGN | fNT | 0.0841 | 0.1490 | 0.5763 | 0.8723 |
| L lingual | L MGN | MD | 0.0679 | 0.1714 | 0.6945 | 0.9987 |
| L lingual | L MGN | PV | 0.0706 | 0.1661 | 0.6736 | 0.9629 |
| L lingual | L MGN | RD | 0.1758 | 0.1707 | 0.3105 | 0.9964 |
| L LOC | L MGN | AD | 0.0727 | 0.1658 | 0.6642 | 0.9951 |
| L LOC | L MGN | FA | 0.1948 | 0.1605 | 0.2340 | 0.9215 |
| L LOC | L MGN | fNT | 0.0624 | 0.1753 | 0.7241 | 0.9189 |
| L LOC | L MGN | MD | 0.1599 | 0.1560 | 0.3130 | 0.9987 |
| L LOC | L MGN | PV | 0.0618 | 0.1702 | 0.7192 | 0.9629 |
| L LOC | L MGN | RD | 0.2016 | 0.1532 | 0.1977 | 0.9964 |
| L Pallidum | L MGN | AD | 0.1414 | 0.1744 | 0.4234 | 0.9951 |
| L Pallidum | L MGN | FA | 0.1514 | 0.1732 | 0.3887 | 0.9215 |
| L Pallidum | L MGN | fNT | 0.0367 | 0.1694 | 0.8301 | 0.9503 |
| L Pallidum | L MGN | MD | 0.1171 | 0.1756 | 0.5098 | 0.9987 |
| L Pallidum | L MGN | PV | 0.1084 | 0.1705 | 0.5295 | 0.9164 |
| L Pallidum | L MGN | RD | 0.0455 | 0.1763 | 0.7981 | 0.9964 |
| L precuneus | L MGN | AD | 0.0147 | 0.1653 | 0.9296 | 0.9951 |
| L precuneus | L MGN | FA | 0.0631 | 0.1691 | 0.7114 | 0.9395 |
| L precuneus | L MGN | fNT | 0.4206 | 0.1601 | 0.0131 | 0.1973 |
| L precuneus | L MGN | MD | 0.0409 | 0.1608 | 0.8006 | 0.9987 |
| L precuneus | L MGN | PV | 0.1782 | 0.1705 | 0.3035 | 0.9039 |
| L precuneus | L MGN | RD | 0.0590 | 0.1589 | 0.7127 | 0.9964 |
| L Putamen | L MGN | AD | 0.0698 | 0.1694 | 0.6832 | 0.9951 |
| L Putamen | L MGN | FA | 0.0301 | 0.1729 | 0.8627 | 0.9772 |
| L Putamen | L MGN | fNT | 0.2749 | 0.1598 | 0.0947 | 0.4301 |
| L Putamen | L MGN | MD | 0.0925 | 0.1686 | 0.5871 | 0.9987 |
| L Putamen | L MGN | PV | 0.1378 | 0.1702 | 0.4239 | 0.9083 |
| L Putamen | L MGN | RD | 0.1010 | 0.1695 | 0.5552 | 0.9964 |
| L rMFC | L MGN | AD | 0.1181 | 0.1715 | 0.4964 | 0.9951 |
| L rMFC | L MGN | FA | 0.0197 | 0.1786 | 0.9131 | 0.9885 |
| L rMFC | L MGN | fNT | 0.2238 | 0.1762 | 0.2138 | 0.6924 |
| L rMFC | L MGN | MD | 0.1359 | 0.1614 | 0.4065 | 0.9987 |
| L rMFC | L MGN | PV | 0.0894 | 0.1818 | 0.6266 | 0.9566 |
| L rMFC | L MGN | RD | 0.1063 | 0.1649 | 0.5243 | 0.9964 |
| L SFC | L MGN | AD | 0.0795 | 0.1664 | 0.6361 | 0.9951 |
| L SFC | L MGN | FA | 0.1886 | 0.1688 | 0.2718 | 0.9215 |
| L SFC | L MGN | fNT | 0.0437 | 0.1717 | 0.8005 | 0.9411 |
| L SFC | L MGN | MD | 0.0384 | 0.1639 | 0.8161 | 0.9987 |
| L SFC | L MGN | PV | 0.0812 | 0.1662 | 0.6284 | 0.9566 |
| L SFC | L MGN | RD | 0.1180 | 0.1642 | 0.4773 | 0.9964 |
| L SPC | L MGN | AD | 0.1166 | 0.1609 | 0.4738 | 0.9951 |
| L SPC | L MGN | FA | 0.1102 | 0.1682 | 0.5169 | 0.9215 |
| L SPC | L MGN | fNT | 0.0448 | 0.1733 | 0.7978 | 0.9411 |
| L SPC | L MGN | MD | 0.0522 | 0.1566 | 0.7409 | 0.9987 |
| L SPC | L MGN | PV | 0.2697 | 0.1641 | 0.1099 | 0.8045 |
| L SPC | L MGN | RD | 0.0011 | 0.1583 | 0.9944 | 0.9964 |
| L TP | L MGN | AD | 0.3479 | 0.1766 | 0.0585 | 0.9951 |
| L TP | L MGN | FA | 0.0546 | 0.1703 | 0.7505 | 0.9510 |
| L TP | L MGN | fNT | 0.1909 | 0.1623 | 0.2481 | 0.6924 |
| L TP | L MGN | MD | 0.0902 | 0.1632 | 0.5840 | 0.9987 |
| L TP | L MGN | PV | 0.0529 | 0.1717 | 0.7601 | 0.9629 |
| L TP | L MGN | RD | 0.0184 | 0.1646 | 0.9118 | 0.9964 |
| L Ventral_Anterior | L MGN | AD | 0.1727 | 0.1589 | 0.2850 | 0.9951 |
| L Ventral_Anterior | L MGN | FA | 0.2372 | 0.1580 | 0.1429 | 0.8663 |
| L Ventral_Anterior | L MGN | fNT | 0.0777 | 0.1708 | 0.6519 | 0.8865 |
| L Ventral_Anterior | L MGN | MD | 0.0231 | 0.1694 | 0.8923 | 0.9987 |
| L Ventral_Anterior | L MGN | PV | 0.1963 | 0.1577 | 0.2219 | 0.9039 |
| L Ventral_Anterior | L MGN | RD | 0.1498 | 0.1721 | 0.3906 | 0.9964 |
| L VLD | L MGN | AD | 0.1035 | 0.1669 | 0.5393 | 0.9951 |
| L VLD | L MGN | FA | 0.0203 | 0.1656 | 0.9033 | 0.9883 |
| L VLD | L MGN | fNT | 0.1942 | 0.1708 | 0.2637 | 0.6948 |
| L VLD | L MGN | MD | 0.1537 | 0.1690 | 0.3695 | 0.9987 |
| L VLD | L MGN | PV | 0.1006 | 0.1647 | 0.5453 | 0.9245 |
| L VLD | L MGN | RD | 0.1451 | 0.1714 | 0.4033 | 0.9964 |
| R Pallidum | L MGN | AD | 0.3680 | 0.1685 | 0.0375 | 0.9951 |
| R Pallidum | L MGN | FA | 0.0803 | 0.1733 | 0.6464 | 0.9287 |
| R Pallidum | L MGN | fNT | 0.0524 | 0.1741 | 0.7653 | 0.9315 |
| R Pallidum | L MGN | MD | 0.1958 | 0.1732 | 0.2666 | 0.9987 |
| R Pallidum | L MGN | PV | 0.0821 | 0.1707 | 0.6338 | 0.9566 |
| R Pallidum | L MGN | RD | 0.2136 | 0.1717 | 0.2225 | 0.9964 |
| L Accumbens | L PAC | AD | 0.0764 | 0.1783 | 0.6714 | 0.9951 |
| L Accumbens | L PAC | FA | 0.1043 | 0.1794 | 0.5650 | 0.9215 |
| L Accumbens | L PAC | fNT | 0.5339 | 0.1664 | 0.0035 | 0.1751 |
| L Accumbens | L PAC | MD | 0.0813 | 0.1750 | 0.6455 | 0.9987 |
| L Accumbens | L PAC | PV | 0.2289 | 0.1751 | 0.2006 | 0.9039 |
| L Accumbens | L PAC | RD | 0.0432 | 0.1759 | 0.8075 | 0.9964 |
| L Amygdala | L PAC | AD | 0.0684 | 0.1715 | 0.6927 | 0.9951 |
| L Amygdala | L PAC | FA | 0.0986 | 0.1598 | 0.5415 | 0.9215 |
| L Amygdala | L PAC | fNT | 0.1866 | 0.1594 | 0.2503 | 0.6924 |
| L Amygdala | L PAC | MD | 0.0049 | 0.1636 | 0.9761 | 0.9987 |
| L Amygdala | L PAC | PV | 0.0931 | 0.1707 | 0.5890 | 0.9566 |
| L Amygdala | L PAC | RD | 0.0518 | 0.1600 | 0.7483 | 0.9964 |
| L Anterior | L PAC | AD | 0.0718 | 0.1729 | 0.6806 | 0.9951 |
| L Anterior | L PAC | FA | 0.3448 | 0.1598 | 0.0385 | 0.5645 |
| L Anterior | L PAC | fNT | 0.0787 | 0.1695 | 0.6457 | 0.8865 |
| L Anterior | L PAC | MD | 0.0666 | 0.1698 | 0.6973 | 0.9987 |
| L Anterior | L PAC | PV | 0.0245 | 0.1733 | 0.8886 | 0.9890 |
| L Anterior | L PAC | RD | 0.1659 | 0.1643 | 0.3202 | 0.9964 |
| L Caudate | L PAC | AD | 0.0309 | 0.1736 | 0.8599 | 0.9951 |
| L Caudate | L PAC | FA | 0.0782 | 0.1703 | 0.6490 | 0.9287 |
| L Caudate | L PAC | fNT | 0.2091 | 0.1701 | 0.2276 | 0.6924 |
| L Caudate | L PAC | MD | 0.0028 | 0.1721 | 0.9870 | 0.9987 |
| L Caudate | L PAC | PV | 0.1988 | 0.1708 | 0.2529 | 0.9039 |
| L Caudate | L PAC | RD | 0.0214 | 0.1715 | 0.9016 | 0.9964 |
| L fusiform | L PAC | AD | 0.0279 | 0.1405 | 0.8440 | 0.9951 |
| L fusiform | L PAC | FA | 0.0631 | 0.1739 | 0.7190 | 0.9395 |
| L fusiform | L PAC | fNT | 0.0644 | 0.1713 | 0.7094 | 0.9153 |
| L fusiform | L PAC | MD | 0.0091 | 0.1481 | 0.9514 | 0.9987 |
| L fusiform | L PAC | PV | 0.0381 | 0.1734 | 0.8274 | 0.9787 |
| L fusiform | L PAC | RD | 0.0321 | 0.1646 | 0.8465 | 0.9964 |
| L Hippo_CA1 | L PAC | AD | 0.3507 | 0.1600 | 0.0360 | 0.9951 |
| L Hippo_CA1 | L PAC | FA | 0.2315 | 0.1727 | 0.1899 | 0.9215 |
| L Hippo_CA1 | L PAC | fNT | 0.0603 | 0.1539 | 0.6978 | 0.9121 |
| L Hippo_CA1 | L PAC | MD | 0.0786 | 0.1737 | 0.6541 | 0.9987 |
| L Hippo_CA1 | L PAC | PV | 0.0039 | 0.1746 | 0.9823 | 0.9996 |
| L Hippo_CA1 | L PAC | RD | 0.1202 | 0.1763 | 0.5003 | 0.9964 |
| L Hippo_HATA | L PAC | AD | 0.2480 | 0.1556 | 0.1210 | 0.9951 |
| L Hippo_HATA | L PAC | FA | 0.2871 | 0.1646 | 0.0911 | 0.7687 |
| L Hippo_HATA | L PAC | fNT | 0.0391 | 0.1659 | 0.8152 | 0.9411 |
| L Hippo_HATA | L PAC | MD | 0.0800 | 0.1619 | 0.6247 | 0.9987 |
| L Hippo_HATA | L PAC | PV | 0.1494 | 0.1669 | 0.3777 | 0.9083 |
| L Hippo_HATA | L PAC | RD | 0.1252 | 0.1691 | 0.4645 | 0.9964 |
| L IPC | L PAC | AD | 0.0843 | 0.1505 | 0.5791 | 0.9951 |
| L IPC | L PAC | FA | 0.0909 | 0.1706 | 0.5978 | 0.9215 |
| L IPC | L PAC | fNT | 0.0964 | 0.1675 | 0.5689 | 0.8723 |
| L IPC | L PAC | MD | 0.0223 | 0.1459 | 0.8797 | 0.9987 |
| L IPC | L PAC | PV | 0.2043 | 0.1579 | 0.2046 | 0.9039 |
| L IPC | L PAC | RD | 0.0297 | 0.1555 | 0.8495 | 0.9964 |
| L ITC | L PAC | AD | 0.0321 | 0.1350 | 0.8138 | 0.9951 |
| L ITC | L PAC | FA | 0.0046 | 0.1650 | 0.9780 | 0.9976 |
| L ITC | L PAC | fNT | 0.2074 | 0.1627 | 0.2114 | 0.6924 |
| L ITC | L PAC | MD | 0.0172 | 0.1466 | 0.9072 | 0.9987 |
| L ITC | L PAC | PV | 0.0684 | 0.1723 | 0.6940 | 0.9629 |
| L ITC | L PAC | RD | 0.0037 | 0.1592 | 0.9817 | 0.9964 |
| L LOC | L PAC | AD | 0.0522 | 0.1593 | 0.7453 | 0.9951 |
| L LOC | L PAC | FA | 0.2692 | 0.1698 | 0.1230 | 0.8168 |
| L LOC | L PAC | fNT | 0.1320 | 0.1749 | 0.4562 | 0.8035 |
| L LOC | L PAC | MD | 0.1561 | 0.1480 | 0.2999 | 0.9987 |
| L LOC | L PAC | PV | 0.1847 | 0.1736 | 0.2958 | 0.9039 |
| L LOC | L PAC | RD | 0.2499 | 0.1553 | 0.1176 | 0.9964 |
| L IOFC | L PAC | AD | 0.1322 | 0.1607 | 0.4166 | 0.9951 |
| L IOFC | L PAC | FA | 0.2787 | 0.1517 | 0.0752 | 0.7269 |
| L IOFC | L PAC | fNT | 0.2421 | 0.1675 | 0.1578 | 0.6086 |
| L IOFC | L PAC | MD | 0.0253 | 0.1417 | 0.8591 | 0.9987 |
| L IOFC | L PAC | PV | 0.1243 | 0.1663 | 0.4601 | 0.9083 |
| L IOFC | L PAC | RD | 0.1549 | 0.1331 | 0.2531 | 0.9964 |
| L Pallidum | L PAC | AD | 0.1693 | 0.1694 | 0.3251 | 0.9951 |
| L Pallidum | L PAC | FA | 0.1520 | 0.1717 | 0.3827 | 0.9215 |
| L Pallidum | L PAC | fNT | 0.0519 | 0.1762 | 0.7700 | 0.9315 |
| L Pallidum | L PAC | MD | 0.1147 | 0.1678 | 0.4991 | 0.9987 |
| L Pallidum | L PAC | PV | 0.1919 | 0.1611 | 0.2423 | 0.9039 |
| L Pallidum | L PAC | RD | 0.0426 | 0.1682 | 0.8018 | 0.9964 |
| L parso | L PAC | AD | 0.0516 | 0.1528 | 0.7376 | 0.9951 |
| L parso | L PAC | FA | 0.0743 | 0.1721 | 0.6688 | 0.9395 |
| L parso | L PAC | fNT | 0.0214 | 0.1703 | 0.9008 | 0.9789 |
| L parso | L PAC | MD | 0.0894 | 0.1579 | 0.5753 | 0.9987 |
| L parso | L PAC | PV | 0.0813 | 0.1757 | 0.6467 | 0.9626 |
| L parso | L PAC | RD | 0.0849 | 0.1694 | 0.6197 | 0.9964 |
| L parstr | L PAC | AD | 0.0236 | 0.1723 | 0.8920 | 0.9951 |
| L parstr | L PAC | FA | 0.0561 | 0.1777 | 0.7542 | 0.9533 |
| L parstr | L PAC | fNT | 0.1825 | 0.1774 | 0.3115 | 0.7391 |
| L parstr | L PAC | MD | 0.0007 | 0.1649 | 0.9966 | 0.9987 |
| L parstr | L PAC | PV | 0.2362 | 0.1710 | 0.1771 | 0.9039 |
| L parstr | L PAC | RD | 0.0232 | 0.1655 | 0.8894 | 0.9964 |
| L Putamen | L PAC | AD | 0.0720 | 0.1649 | 0.6654 | 0.9951 |
| L Putamen | L PAC | FA | 0.0044 | 0.1656 | 0.9790 | 0.9976 |
| L Putamen | L PAC | fNT | 0.1246 | 0.1723 | 0.4746 | 0.8207 |
| L Putamen | L PAC | MD | 0.0716 | 0.1578 | 0.6529 | 0.9987 |
| L Putamen | L PAC | PV | 0.1985 | 0.1671 | 0.2434 | 0.9039 |
| L Putamen | L PAC | RD | 0.0555 | 0.1559 | 0.7239 | 0.9964 |
| L rMFC | L PAC | AD | 0.0628 | 0.1667 | 0.7088 | 0.9951 |
| L rMFC | L PAC | FA | 0.2219 | 0.1654 | 0.1888 | 0.9215 |
| L rMFC | L PAC | fNT | 0.2510 | 0.1672 | 0.1428 | 0.5878 |
| L rMFC | L PAC | MD | 0.0508 | 0.1610 | 0.7542 | 0.9987 |
| L rMFC | L PAC | PV | 0.1544 | 0.1701 | 0.3705 | 0.9083 |
| L rMFC | L PAC | RD | 0.1448 | 0.1590 | 0.3690 | 0.9964 |
| L SFC | L PAC | AD | 0.0416 | 0.1634 | 0.8006 | 0.9951 |
| L SFC | L PAC | FA | 0.0924 | 0.1675 | 0.5851 | 0.9215 |
| L SFC | L PAC | fNT | 0.2251 | 0.1650 | 0.1819 | 0.6573 |
| L SFC | L PAC | MD | 0.0349 | 0.1538 | 0.8217 | 0.9987 |
| L SFC | L PAC | PV | 0.1580 | 0.1391 | 0.2644 | 0.9039 |
| L SFC | L PAC | RD | 0.0699 | 0.1575 | 0.6599 | 0.9964 |
| L SPC | L PAC | AD | 0.2346 | 0.1661 | 0.1671 | 0.9951 |
| L SPC | L PAC | FA | 0.3079 | 0.1551 | 0.0555 | 0.6458 |
| L SPC | L PAC | fNT | 0.3207 | 0.1701 | 0.0688 | 0.4089 |
| L SPC | L PAC | MD | 0.0055 | 0.1508 | 0.9713 | 0.9987 |
| L SPC | L PAC | PV | 0.0642 | 0.1700 | 0.7082 | 0.9629 |
| L SPC | L PAC | RD | 0.1698 | 0.1482 | 0.2602 | 0.9964 |
| L TP | L PAC | AD | 0.0420 | 0.1577 | 0.7919 | 0.9951 |
| L TP | L PAC | FA | 0.1444 | 0.1631 | 0.3823 | 0.9215 |
| L TP | L PAC | fNT | 0.2177 | 0.1665 | 0.2000 | 0.6924 |
| L TP | L PAC | MD | 0.1146 | 0.1491 | 0.4478 | 0.9987 |
| L TP | L PAC | PV | 0.1101 | 0.1630 | 0.5041 | 0.9105 |
| L TP | L PAC | RD | 0.1404 | 0.1507 | 0.3582 | 0.9964 |
| L Ventral_Anterior | L PAC | AD | 0.1683 | 0.1704 | 0.3305 | 0.9951 |
| L Ventral_Anterior | L PAC | FA | 0.1601 | 0.1648 | 0.3383 | 0.9215 |
| L Ventral_Anterior | L PAC | fNT | 0.0186 | 0.1631 | 0.9101 | 0.9789 |
| L Ventral_Anterior | L PAC | MD | 0.1060 | 0.1703 | 0.5380 | 0.9987 |
| L Ventral_Anterior | L PAC | PV | 0.0175 | 0.1740 | 0.9206 | 0.9984 |
| L Ventral_Anterior | L PAC | RD | 0.0279 | 0.1693 | 0.8702 | 0.9964 |
| L VLD | L PAC | AD | 0.0162 | 0.1733 | 0.9262 | 0.9951 |
| L VLD | L PAC | FA | 0.1465 | 0.1678 | 0.3889 | 0.9215 |
| L VLD | L PAC | fNT | 0.3047 | 0.1702 | 0.0832 | 0.4089 |
| L VLD | L PAC | MD | 0.1352 | 0.1713 | 0.4356 | 0.9987 |
| L VLD | L PAC | PV | 0.0535 | 0.1736 | 0.7600 | 0.9629 |
| L VLD | L PAC | RD | 0.3729 | 0.1702 | 0.0366 | 0.9964 |
| R Pallidum | L PAC | AD | 0.3416 | 0.1674 | 0.0502 | 0.9951 |
| R Pallidum | L PAC | FA | 0.0481 | 0.1628 | 0.7696 | 0.9557 |
| R Pallidum | L PAC | fNT | 0.1895 | 0.1672 | 0.2651 | 0.6948 |
| R Pallidum | L PAC | MD | 0.1850 | 0.1632 | 0.2649 | 0.9987 |
| R Pallidum | L PAC | PV | 0.4071 | 0.1550 | 0.0131 | 0.4604 |
| R Pallidum | L PAC | RD | 0.1363 | 0.1605 | 0.4021 | 0.9964 |
| R Putamen | L PAC | AD | 0.0011 | 0.1841 | 0.9951 | 0.9951 |
| R Putamen | L PAC | FA | 0.2092 | 0.1814 | 0.2579 | 0.9215 |
| R Putamen | L PAC | fNT | 0.0223 | 0.1810 | 0.9029 | 0.9789 |
| R Putamen | L PAC | MD | 0.0938 | 0.1840 | 0.6139 | 0.9987 |
| R Putamen | L PAC | PV | 0.2398 | 0.1717 | 0.1727 | 0.9039 |
| R Putamen | L PAC | RD | 0.1328 | 0.1837 | 0.4756 | 0.9964 |
| L Anterior | L CN/SOC | AD | 0.0091 | 0.1728 | 0.9582 | 0.9951 |
| L Anterior | L CN/SOC | FA | 0.0293 | 0.1733 | 0.8666 | 0.9772 |
| L Anterior | L CN/SOC | fNT | 0.1808 | 0.1706 | 0.2972 | 0.7290 |
| L Anterior | L CN/SOC | MD | 0.0235 | 0.1730 | 0.8928 | 0.9987 |
| L Anterior | L CN/SOC | PV | 0.1721 | 0.1703 | 0.3196 | 0.9039 |
| L Anterior | L CN/SOC | RD | 0.0500 | 0.1732 | 0.7747 | 0.9964 |
| L Caudate | L CN/SOC | AD | 0.0376 | 0.1753 | 0.8314 | 0.9951 |
| L Caudate | L CN/SOC | FA | 0.1583 | 0.1738 | 0.3693 | 0.9215 |
| L Caudate | L CN/SOC | fNT | 0.0068 | 0.1695 | 0.9681 | 0.9899 |
| L Caudate | L CN/SOC | MD | 0.0402 | 0.1724 | 0.8170 | 0.9987 |
| L Caudate | L CN/SOC | PV | 0.1501 | 0.1746 | 0.3963 | 0.9083 |
| L Caudate | L CN/SOC | RD | 0.1252 | 0.1690 | 0.4642 | 0.9964 |
| L Hippo_CA1 | L CN/SOC | AD | 0.0289 | 0.1727 | 0.8683 | 0.9951 |
| L Hippo_CA1 | L CN/SOC | FA | 0.0620 | 0.1737 | 0.7236 | 0.9395 |
| L Hippo_CA1 | L CN/SOC | fNT | 0.3340 | 0.1814 | 0.0765 | 0.4089 |
| L Hippo_CA1 | L CN/SOC | MD | 0.0075 | 0.1719 | 0.9654 | 0.9987 |
| L Hippo_CA1 | L CN/SOC | PV | 0.2597 | 0.1655 | 0.1263 | 0.8585 |
| L Hippo_CA1 | L CN/SOC | RD | 0.0426 | 0.1724 | 0.8064 | 0.9964 |
| L Pallidum | L CN/SOC | AD | 0.0064 | 0.1767 | 0.9712 | 0.9951 |
| L Pallidum | L CN/SOC | FA | 0.0027 | 0.1754 | 0.9879 | 0.9983 |
| L Pallidum | L CN/SOC | fNT | 0.1175 | 0.1601 | 0.4685 | 0.8129 |
| L Pallidum | L CN/SOC | MD | 0.0061 | 0.1763 | 0.9728 | 0.9987 |
| L Pallidum | L CN/SOC | PV | 0.0539 | 0.1605 | 0.7393 | 0.9629 |
| L Pallidum | L CN/SOC | RD | 0.0184 | 0.1756 | 0.9171 | 0.9964 |
| L Putamen | L CN/SOC | AD | 0.0101 | 0.1739 | 0.9542 | 0.9951 |
| L Putamen | L CN/SOC | FA | 0.1756 | 0.1699 | 0.3089 | 0.9215 |
| L Putamen | L CN/SOC | fNT | 0.1260 | 0.1577 | 0.4300 | 0.7928 |
| L Putamen | L CN/SOC | MD | 0.0756 | 0.1734 | 0.6659 | 0.9987 |
| L Putamen | L CN/SOC | PV | 0.0384 | 0.1733 | 0.8259 | 0.9787 |
| L Putamen | L CN/SOC | RD | 0.1322 | 0.1720 | 0.4476 | 0.9964 |
| L rMFC | L CN/SOC | AD | 0.0490 | 0.1694 | 0.7742 | 0.9951 |
| L rMFC | L CN/SOC | FA | 0.1460 | 0.1719 | 0.4023 | 0.9215 |
| L rMFC | L CN/SOC | fNT | 0.1768 | 0.1756 | 0.3218 | 0.7420 |
| L rMFC | L CN/SOC | MD | 0.1418 | 0.1589 | 0.3792 | 0.9987 |
| L rMFC | L CN/SOC | PV | 0.1323 | 0.1763 | 0.4585 | 0.9083 |
| L rMFC | L CN/SOC | RD | 0.1763 | 0.1593 | 0.2768 | 0.9964 |
| L SFC | L CN/SOC | AD | 0.0921 | 0.1682 | 0.5877 | 0.9951 |
| L SFC | L CN/SOC | FA | 0.1248 | 0.1678 | 0.4625 | 0.9215 |
| L SFC | L CN/SOC | fNT | 0.0772 | 0.1592 | 0.6308 | 0.8849 |
| L SFC | L CN/SOC | MD | 0.0115 | 0.1663 | 0.9452 | 0.9987 |
| L SFC | L CN/SOC | PV | 0.1270 | 0.1658 | 0.4491 | 0.9083 |
| L SFC | L CN/SOC | RD | 0.0802 | 0.1668 | 0.6338 | 0.9964 |
| L SPC | L CN/SOC | AD | 0.0954 | 0.1781 | 0.5959 | 0.9951 |
| L SPC | L CN/SOC | FA | 0.0984 | 0.1782 | 0.5846 | 0.9215 |
| L SPC | L CN/SOC | fNT | 0.4041 | 0.1690 | 0.0235 | 0.2400 |
| L SPC | L CN/SOC | MD | 0.1479 | 0.1763 | 0.4079 | 0.9987 |
| L SPC | L CN/SOC | PV | 0.1855 | 0.1746 | 0.2963 | 0.9039 |
| L SPC | L CN/SOC | RD | 0.1678 | 0.1754 | 0.3460 | 0.9964 |
| L VLD | L CN/SOC | AD | 0.0105 | 0.1581 | 0.9477 | 0.9951 |
| L VLD | L CN/SOC | FA | 0.0111 | 0.1557 | 0.9434 | 0.9959 |
| L VLD | L CN/SOC | fNT | 0.1774 | 0.1661 | 0.2936 | 0.7246 |
| L VLD | L CN/SOC | MD | 0.0065 | 0.1734 | 0.9705 | 0.9987 |
| L VLD | L CN/SOC | PV | 0.1074 | 0.1762 | 0.5465 | 0.9245 |
| L VLD | L CN/SOC | RD | 0.0208 | 0.1720 | 0.9043 | 0.9964 |
| R Pallidum | L CN/SOC | AD | 0.0014 | 0.1739 | 0.9936 | 0.9951 |
| R Pallidum | L CN/SOC | FA | 0.1530 | 0.1654 | 0.3618 | 0.9215 |
| R Pallidum | L CN/SOC | fNT | 0.2980 | 0.1664 | 0.0825 | 0.4089 |
| R Pallidum | L CN/SOC | MD | 0.0798 | 0.1724 | 0.6465 | 0.9987 |
| R Pallidum | L CN/SOC | PV | 0.0864 | 0.1700 | 0.6147 | 0.9566 |
| R Pallidum | L CN/SOC | RD | 0.1612 | 0.1657 | 0.3377 | 0.9964 |
| R Putamen | L CN/SOC | AD | 0.0693 | 0.1755 | 0.6955 | 0.9951 |
| R Putamen | L CN/SOC | FA | 0.0698 | 0.1696 | 0.6834 | 0.9395 |
| R Putamen | L CN/SOC | fNT | 0.1449 | 0.1730 | 0.4083 | 0.7882 |
| R Putamen | L CN/SOC | MD | 0.0401 | 0.1745 | 0.8196 | 0.9987 |
| R Putamen | L CN/SOC | PV | 0.1430 | 0.1741 | 0.4177 | 0.9083 |
| R Putamen | L CN/SOC | RD | 0.0032 | 0.1708 | 0.9853 | 0.9964 |
| R SFC | L CN/SOC | AD | 0.1453 | 0.1673 | 0.3912 | 0.9951 |
| R SFC | L CN/SOC | FA | 0.3321 | 0.1640 | 0.0516 | 0.6270 |
| R SFC | L CN/SOC | fNT | 0.1381 | 0.1696 | 0.4211 | 0.7928 |
| R SFC | L CN/SOC | MD | 0.0568 | 0.1664 | 0.7350 | 0.9987 |
| R SFC | L CN/SOC | PV | 0.1263 | 0.1686 | 0.4588 | 0.9083 |
| R SFC | L CN/SOC | RD | 0.0646 | 0.1615 | 0.6917 | 0.9964 |
| R Anterior | R IC | AD | 0.1148 | 0.1723 | 0.5099 | 0.9951 |
| R Anterior | R IC | FA | 0.2006 | 0.1496 | 0.1893 | 0.9215 |
| R Anterior | R IC | fNT | 0.0571 | 0.1697 | 0.7387 | 0.9240 |
| R Anterior | R IC | MD | 0.0159 | 0.1728 | 0.9274 | 0.9987 |
| R Anterior | R IC | PV | 0.2629 | 0.1657 | 0.1223 | 0.8458 |
| R Anterior | R IC | RD | 0.0672 | 0.1660 | 0.6881 | 0.9964 |
| R Caudate | R IC | AD | 0.0741 | 0.1746 | 0.6740 | 0.9951 |
| R Caudate | R IC | FA | 0.0075 | 0.1748 | 0.9660 | 0.9976 |
| R Caudate | R IC | fNT | 0.2140 | 0.1676 | 0.2108 | 0.6924 |
| R Caudate | R IC | MD | 0.0695 | 0.1694 | 0.6845 | 0.9987 |
| R Caudate | R IC | PV | 0.1247 | 0.1736 | 0.4776 | 0.9099 |
| R Caudate | R IC | RD | 0.0465 | 0.1663 | 0.7815 | 0.9964 |
| R Medio_Dorsal | R IC | AD | 0.1039 | 0.1756 | 0.5581 | 0.9951 |
| R Medio_Dorsal | R IC | FA | 0.3259 | 0.1484 | 0.0354 | 0.5348 |
| R Medio_Dorsal | R IC | fNT | 0.0146 | 0.1552 | 0.9257 | 0.9848 |
| R Medio_Dorsal | R IC | MD | 0.0152 | 0.1766 | 0.9320 | 0.9987 |
| R Medio_Dorsal | R IC | PV | 0.2902 | 0.1688 | 0.0952 | 0.7648 |
| R Medio_Dorsal | R IC | RD | 0.0719 | 0.1726 | 0.6799 | 0.9964 |
| R Pallidum | R IC | AD | 0.1031 | 0.1728 | 0.5548 | 0.9951 |
| R Pallidum | R IC | FA | 0.3442 | 0.1612 | 0.0403 | 0.5732 |
| R Pallidum | R IC | fNT | 0.2252 | 0.1574 | 0.1619 | 0.6153 |
| R Pallidum | R IC | MD | 0.1323 | 0.1724 | 0.4481 | 0.9987 |
| R Pallidum | R IC | PV | 0.1723 | 0.1716 | 0.3227 | 0.9039 |
| R Pallidum | R IC | RD | 0.2910 | 0.1652 | 0.0875 | 0.9964 |
| R Putamen | R IC | AD | 0.1327 | 0.1713 | 0.4444 | 0.9951 |
| R Putamen | R IC | FA | 0.1421 | 0.1747 | 0.4219 | 0.9215 |
| R Putamen | R IC | fNT | 0.2343 | 0.1615 | 0.1566 | 0.6086 |
| R Putamen | R IC | MD | 0.0560 | 0.1718 | 0.7467 | 0.9987 |
| R Putamen | R IC | PV | 0.1597 | 0.1743 | 0.3665 | 0.9083 |
| R Putamen | R IC | RD | 0.0311 | 0.1733 | 0.8588 | 0.9964 |
| R SFC | R IC | AD | 0.0284 | 0.1669 | 0.8659 | 0.9951 |
| R SFC | R IC | FA | 0.1146 | 0.1597 | 0.4779 | 0.9215 |
| R SFC | R IC | fNT | 0.0229 | 0.1726 | 0.8954 | 0.9789 |
| R SFC | R IC | MD | 0.0258 | 0.1634 | 0.8757 | 0.9987 |
| R SFC | R IC | PV | 0.0461 | 0.1705 | 0.7885 | 0.9629 |
| R SFC | R IC | RD | 0.0616 | 0.1593 | 0.7017 | 0.9964 |
| R SPC | R IC | AD | 0.0042 | 0.1749 | 0.9808 | 0.9951 |
| R SPC | R IC | FA | 0.2361 | 0.1704 | 0.1759 | 0.9215 |
| R SPC | R IC | fNT | 0.0137 | 0.1788 | 0.9395 | 0.9882 |
| R SPC | R IC | MD | 0.1180 | 0.1709 | 0.4950 | 0.9987 |
| R SPC | R IC | PV | 0.1829 | 0.1762 | 0.3073 | 0.9039 |
| R SPC | R IC | RD | 0.1792 | 0.1694 | 0.2984 | 0.9964 |
| R Amygdala | R MGN | AD | 0.0652 | 0.1737 | 0.7099 | 0.9951 |
| R Amygdala | R MGN | FA | 0.1213 | 0.1702 | 0.4810 | 0.9215 |
| R Amygdala | R MGN | fNT | 0.1009 | 0.1693 | 0.5555 | 0.8723 |
| R Amygdala | R MGN | MD | 0.1933 | 0.1678 | 0.2574 | 0.9987 |
| R Amygdala | R MGN | PV | 0.2631 | 0.1581 | 0.1056 | 0.7850 |
| R Amygdala | R MGN | RD | 0.2033 | 0.1662 | 0.2300 | 0.9964 |
| R Anterior | R MGN | AD | 0.0537 | 0.1728 | 0.7581 | 0.9951 |
| R Anterior | R MGN | FA | 0.0840 | 0.1709 | 0.6265 | 0.9260 |
| R Anterior | R MGN | fNT | 0.3690 | 0.1597 | 0.0275 | 0.2590 |
| R Anterior | R MGN | MD | 0.1261 | 0.1682 | 0.4586 | 0.9987 |
| R Anterior | R MGN | PV | 0.4063 | 0.1555 | 0.0137 | 0.4604 |
| R Anterior | R MGN | RD | 0.3671 | 0.1535 | 0.0231 | 0.9964 |
| R Caudate | R MGN | AD | 0.4970 | 0.1760 | 0.0096 | 0.9951 |
| R Caudate | R MGN | FA | 0.1582 | 0.1703 | 0.3598 | 0.9215 |
| R Caudate | R MGN | fNT | 0.2082 | 0.1662 | 0.2191 | 0.6924 |
| R Caudate | R MGN | MD | 0.3181 | 0.1531 | 0.0461 | 0.9987 |
| R Caudate | R MGN | PV | 0.0559 | 0.1696 | 0.7437 | 0.9629 |
| R Caudate | R MGN | RD | 0.0044 | 0.1670 | 0.9793 | 0.9964 |
| R entorhinal | R MGN | AD | 0.0198 | 0.1667 | 0.9062 | 0.9951 |
| R entorhinal | R MGN | FA | 0.4498 | 0.1654 | 0.0109 | 0.4498 |
| R entorhinal | R MGN | fNT | 0.0780 | 0.1694 | 0.6483 | 0.8865 |
| R entorhinal | R MGN | MD | 0.2195 | 0.1622 | 0.1851 | 0.9987 |
| R entorhinal | R MGN | PV | 0.0584 | 0.1698 | 0.7332 | 0.9629 |
| R entorhinal | R MGN | RD | 0.3345 | 0.1595 | 0.0436 | 0.9964 |
| R fusiform | R MGN | AD | 0.0320 | 0.1667 | 0.8489 | 0.9951 |
| R fusiform | R MGN | FA | 0.1108 | 0.1728 | 0.5259 | 0.9215 |
| R fusiform | R MGN | fNT | 0.0073 | 0.1685 | 0.9656 | 0.9899 |
| R fusiform | R MGN | MD | 0.1366 | 0.1665 | 0.4179 | 0.9987 |
| R fusiform | R MGN | PV | 0.1925 | 0.1693 | 0.2639 | 0.9039 |
| R fusiform | R MGN | RD | 0.1495 | 0.1695 | 0.3843 | 0.9964 |
| R Hippo_CA1 | R MGN | AD | 0.0162 | 0.1796 | 0.9289 | 0.9951 |
| R Hippo_CA1 | R MGN | FA | 0.0829 | 0.1752 | 0.6393 | 0.9281 |
| R Hippo_CA1 | R MGN | fNT | 0.5010 | 0.1726 | 0.0078 | 0.1751 |
| R Hippo_CA1 | R MGN | MD | 0.0523 | 0.1783 | 0.7711 | 0.9987 |
| R Hippo_CA1 | R MGN | PV | 0.3820 | 0.1684 | 0.0310 | 0.4974 |
| R Hippo_CA1 | R MGN | RD | 0.0582 | 0.1771 | 0.7446 | 0.9964 |
| R IPC | R MGN | AD | 0.0398 | 0.1602 | 0.8056 | 0.9951 |
| R IPC | R MGN | FA | 0.2652 | 0.1594 | 0.1060 | 0.8052 |
| R IPC | R MGN | fNT | 0.0226 | 0.1743 | 0.8978 | 0.9789 |
| R IPC | R MGN | MD | 0.0859 | 0.1459 | 0.5601 | 0.9987 |
| R IPC | R MGN | PV | 0.0293 | 0.1742 | 0.8673 | 0.9844 |
| R IPC | R MGN | RD | 0.1537 | 0.1447 | 0.2962 | 0.9964 |
| R ITC | R MGN | AD | 0.0895 | 0.1592 | 0.5777 | 0.9951 |
| R ITC | R MGN | FA | 0.0083 | 0.1694 | 0.9610 | 0.9959 |
| R ITC | R MGN | fNT | 0.3846 | 0.1507 | 0.0155 | 0.2033 |
| R ITC | R MGN | MD | 0.0672 | 0.1627 | 0.6822 | 0.9987 |
| R ITC | R MGN | PV | 0.3647 | 0.1513 | 0.0217 | 0.4604 |
| R ITC | R MGN | RD | 0.0385 | 0.1666 | 0.8185 | 0.9964 |
| R Medio_Dorsal | R MGN | AD | 0.0374 | 0.1745 | 0.8316 | 0.9951 |
| R Medio_Dorsal | R MGN | FA | 0.1760 | 0.1748 | 0.3215 | 0.9215 |
| R Medio_Dorsal | R MGN | fNT | 0.2997 | 0.1616 | 0.0729 | 0.4089 |
| R Medio_Dorsal | R MGN | MD | 0.1905 | 0.1674 | 0.2636 | 0.9987 |
| R Medio_Dorsal | R MGN | PV | 0.1262 | 0.1714 | 0.4669 | 0.9083 |
| R Medio_Dorsal | R MGN | RD | 0.4784 | 0.1464 | 0.0027 | 0.7924 |
| R Pallidum | R MGN | AD | 0.0514 | 0.1697 | 0.7639 | 0.9951 |
| R Pallidum | R MGN | FA | 0.3194 | 0.1530 | 0.0446 | 0.5769 |
| R Pallidum | R MGN | fNT | 0.0464 | 0.1695 | 0.7859 | 0.9376 |
| R Pallidum | R MGN | MD | 0.1467 | 0.1686 | 0.3905 | 0.9987 |
| R Pallidum | R MGN | PV | 0.0705 | 0.1679 | 0.6773 | 0.9629 |
| R Pallidum | R MGN | RD | 0.3972 | 0.1470 | 0.0110 | 0.9678 |
| R precuneus | R MGN | AD | 0.1449 | 0.1604 | 0.3730 | 0.9951 |
| R precuneus | R MGN | FA | 0.3420 | 0.1689 | 0.0516 | 0.6270 |
| R precuneus | R MGN | fNT | 0.0017 | 0.1672 | 0.9920 | 0.9957 |
| R precuneus | R MGN | MD | 0.1043 | 0.1615 | 0.5228 | 0.9987 |
| R precuneus | R MGN | PV | 0.3257 | 0.1588 | 0.0483 | 0.5482 |
| R precuneus | R MGN | RD | 0.2237 | 0.1658 | 0.1865 | 0.9964 |
| R Putamen | R MGN | AD | 0.0065 | 0.1727 | 0.9703 | 0.9951 |
| R Putamen | R MGN | FA | 0.1446 | 0.1677 | 0.3949 | 0.9215 |
| R Putamen | R MGN | fNT | 0.1457 | 0.1659 | 0.3865 | 0.7832 |
| R Putamen | R MGN | MD | 0.0684 | 0.1674 | 0.6854 | 0.9987 |
| R Putamen | R MGN | PV | 0.1901 | 0.1690 | 0.2691 | 0.9039 |
| R Putamen | R MGN | RD | 0.1176 | 0.1619 | 0.4727 | 0.9964 |
| R rMFC | R MGN | AD | 0.0728 | 0.1660 | 0.6641 | 0.9951 |
| R rMFC | R MGN | FA | 0.0478 | 0.1708 | 0.7813 | 0.9557 |
| R rMFC | R MGN | fNT | 0.3036 | 0.1706 | 0.0852 | 0.4089 |
| R rMFC | R MGN | MD | 0.0359 | 0.1663 | 0.8306 | 0.9987 |
| R rMFC | R MGN | PV | 0.0936 | 0.1710 | 0.5880 | 0.9566 |
| R rMFC | R MGN | RD | 0.0067 | 0.1680 | 0.9685 | 0.9964 |
| R SFC | R MGN | AD | 0.0542 | 0.1683 | 0.7493 | 0.9951 |
| R SFC | R MGN | FA | 0.0663 | 0.1596 | 0.6804 | 0.9395 |
| R SFC | R MGN | fNT | 0.1381 | 0.1701 | 0.4226 | 0.7928 |
| R SFC | R MGN | MD | 0.0140 | 0.1616 | 0.9313 | 0.9987 |
| R SFC | R MGN | PV | 0.1704 | 0.1617 | 0.2998 | 0.9039 |
| R SFC | R MGN | RD | 0.0630 | 0.1555 | 0.6881 | 0.9964 |
| R SPC | R MGN | AD | 0.1208 | 0.1632 | 0.4648 | 0.9951 |
| R SPC | R MGN | FA | 0.4416 | 0.1542 | 0.0073 | 0.4498 |
| R SPC | R MGN | fNT | 0.1417 | 0.1733 | 0.4197 | 0.7928 |
| R SPC | R MGN | MD | 0.1215 | 0.1602 | 0.4536 | 0.9987 |
| R SPC | R MGN | PV | 0.0373 | 0.1690 | 0.8269 | 0.9787 |
| R SPC | R MGN | RD | 0.2755 | 0.1571 | 0.0890 | 0.9964 |
| R Ventral_Anterior | R MGN | AD | 0.2002 | 0.1666 | 0.2380 | 0.9951 |
| R Ventral_Anterior | R MGN | FA | 0.2564 | 0.1661 | 0.1322 | 0.8439 |
| R Ventral_Anterior | R MGN | fNT | 0.3249 | 0.1667 | 0.0607 | 0.3827 |
| R Ventral_Anterior | R MGN | MD | 0.1155 | 0.1700 | 0.5017 | 0.9987 |
| R Ventral_Anterior | R MGN | PV | 0.4003 | 0.1562 | 0.0153 | 0.4604 |
| R Ventral_Anterior | R MGN | RD | 0.0286 | 0.1735 | 0.8700 | 0.9964 |
| R Amygdala | R PAC | AD | 0.0815 | 0.1639 | 0.6223 | 0.9951 |
| R Amygdala | R PAC | FA | 0.1469 | 0.1677 | 0.3873 | 0.9215 |
| R Amygdala | R PAC | fNT | 0.0053 | 0.1710 | 0.9753 | 0.9899 |
| R Amygdala | R PAC | MD | 0.0078 | 0.1552 | 0.9603 | 0.9987 |
| R Amygdala | R PAC | PV | 0.0965 | 0.1639 | 0.5599 | 0.9325 |
| R Amygdala | R PAC | RD | 0.0662 | 0.1611 | 0.6838 | 0.9964 |
| R Anterior | R PAC | AD | 0.0214 | 0.1732 | 0.9024 | 0.9951 |
| R Anterior | R PAC | FA | 0.0402 | 0.1744 | 0.8191 | 0.9559 |
| R Anterior | R PAC | fNT | 0.0527 | 0.1694 | 0.7576 | 0.9315 |
| R Anterior | R PAC | MD | 0.0145 | 0.1666 | 0.9311 | 0.9987 |
| R Anterior | R PAC | PV | 0.1117 | 0.1656 | 0.5049 | 0.9105 |
| R Anterior | R PAC | RD | 0.0474 | 0.1648 | 0.7753 | 0.9964 |
| R Caudate | R PAC | AD | 0.0756 | 0.1699 | 0.6592 | 0.9951 |
| R Caudate | R PAC | FA | 0.2007 | 0.1731 | 0.2550 | 0.9215 |
| R Caudate | R PAC | fNT | 0.4498 | 0.1375 | 0.0028 | 0.1751 |
| R Caudate | R PAC | MD | 0.1361 | 0.1698 | 0.4288 | 0.9987 |
| R Caudate | R PAC | PV | 0.1101 | 0.1729 | 0.5288 | 0.9164 |
| R Caudate | R PAC | RD | 0.1778 | 0.1703 | 0.3044 | 0.9964 |
| R fusiform | R PAC | AD | 0.1297 | 0.1435 | 0.3729 | 0.9951 |
| R fusiform | R PAC | FA | 0.2179 | 0.1709 | 0.2117 | 0.9215 |
| R fusiform | R PAC | fNT | 0.2542 | 0.1665 | 0.1367 | 0.5674 |
| R fusiform | R PAC | MD | 0.0750 | 0.1593 | 0.6410 | 0.9987 |
| R fusiform | R PAC | PV | 0.1541 | 0.1656 | 0.3591 | 0.9083 |
| R fusiform | R PAC | RD | 0.1662 | 0.1705 | 0.3372 | 0.9964 |
| R IPC | R PAC | AD | 0.0133 | 0.1505 | 0.9300 | 0.9951 |
| R IPC | R PAC | FA | 0.1471 | 0.1715 | 0.3971 | 0.9215 |
| R IPC | R PAC | fNT | 0.3265 | 0.1512 | 0.0382 | 0.3018 |
| R IPC | R PAC | MD | 0.0280 | 0.1485 | 0.8515 | 0.9987 |
| R IPC | R PAC | PV | 0.1376 | 0.1463 | 0.3538 | 0.9083 |
| R IPC | R PAC | RD | 0.0565 | 0.1544 | 0.7170 | 0.9964 |
| R ITC | R PAC | AD | 0.0063 | 0.1242 | 0.9601 | 0.9951 |
| R ITC | R PAC | FA | 0.0145 | 0.1721 | 0.9332 | 0.9933 |
| R ITC | R PAC | fNT | 0.3219 | 0.1574 | 0.0488 | 0.3474 |
| R ITC | R PAC | MD | 0.0272 | 0.1371 | 0.8442 | 0.9987 |
| R ITC | R PAC | PV | 0.1976 | 0.1491 | 0.1943 | 0.9039 |
| R ITC | R PAC | RD | 0.0453 | 0.1591 | 0.7777 | 0.9964 |
| R LOC | R PAC | AD | 0.0070 | 0.1450 | 0.9616 | 0.9951 |
| R LOC | R PAC | FA | 0.0986 | 0.1729 | 0.5721 | 0.9215 |
| R LOC | R PAC | fNT | 0.0042 | 0.1742 | 0.9808 | 0.9899 |
| R LOC | R PAC | MD | 0.0669 | 0.1432 | 0.6433 | 0.9987 |
| R LOC | R PAC | PV | 0.0446 | 0.1717 | 0.7964 | 0.9629 |
| R LOC | R PAC | RD | 0.1010 | 0.1554 | 0.5201 | 0.9964 |
| R IOFC | R PAC | AD | 0.0796 | 0.1585 | 0.6188 | 0.9951 |
| R IOFC | R PAC | FA | 0.1201 | 0.1644 | 0.4703 | 0.9215 |
| R IOFC | R PAC | fNT | 0.1649 | 0.1680 | 0.3334 | 0.7535 |
| R IOFC | R PAC | MD | 0.0076 | 0.1480 | 0.9592 | 0.9987 |
| R IOFC | R PAC | PV | 0.3571 | 0.1597 | 0.0324 | 0.5044 |
| R IOFC | R PAC | RD | 0.0568 | 0.1518 | 0.7105 | 0.9964 |
| R Pallidum | R PAC | AD | 0.3303 | 0.1469 | 0.0319 | 0.9951 |
| R Pallidum | R PAC | FA | 0.2859 | 0.1538 | 0.0720 | 0.7179 |
| R Pallidum | R PAC | fNT | 0.0863 | 0.1690 | 0.6128 | 0.8733 |
| R Pallidum | R PAC | MD | 0.0110 | 0.1511 | 0.9424 | 0.9987 |
| R Pallidum | R PAC | PV | 0.1245 | 0.1629 | 0.4502 | 0.9083 |
| R Pallidum | R PAC | RD | 0.1080 | 0.1549 | 0.4907 | 0.9964 |
| R pars | R PAC | AD | 0.1058 | 0.1488 | 0.4823 | 0.9951 |
| R pars | R PAC | FA | 0.0193 | 0.1573 | 0.9031 | 0.9883 |
| R pars | R PAC | fNT | 0.2849 | 0.1725 | 0.1095 | 0.4725 |
| R pars | R PAC | MD | 0.0599 | 0.1462 | 0.6849 | 0.9987 |
| R pars | R PAC | PV | 0.0348 | 0.1651 | 0.8346 | 0.9799 |
| R pars | R PAC | RD | 0.0368 | 0.1479 | 0.8052 | 0.9964 |
| R parstr | R PAC | AD | 0.1237 | 0.1550 | 0.4308 | 0.9951 |
| R parstr | R PAC | FA | 0.1243 | 0.1628 | 0.4506 | 0.9215 |
| R parstr | R PAC | fNT | 0.0124 | 0.1744 | 0.9439 | 0.9882 |
| R parstr | R PAC | MD | 0.0721 | 0.1517 | 0.6377 | 0.9987 |
| R parstr | R PAC | PV | 0.1855 | 0.1695 | 0.2821 | 0.9039 |
| R parstr | R PAC | RD | 0.0287 | 0.1542 | 0.8536 | 0.9964 |
| R Putamen | R PAC | AD | 0.0362 | 0.1367 | 0.7929 | 0.9951 |
| R Putamen | R PAC | FA | 0.1601 | 0.1639 | 0.3359 | 0.9215 |
| R Putamen | R PAC | fNT | 0.1535 | 0.1719 | 0.3785 | 0.7794 |
| R Putamen | R PAC | MD | 0.0311 | 0.1418 | 0.8279 | 0.9987 |
| R Putamen | R PAC | PV | 0.1763 | 0.1637 | 0.2897 | 0.9039 |
| R Putamen | R PAC | RD | 0.0761 | 0.1533 | 0.6228 | 0.9964 |
| R SPC | R PAC | AD | 0.0563 | 0.1449 | 0.6999 | 0.9951 |
| R SPC | R PAC | FA | 0.1366 | 0.1714 | 0.4309 | 0.9215 |
| R SPC | R PAC | fNT | 0.0270 | 0.1685 | 0.8735 | 0.9688 |
| R SPC | R PAC | MD | 0.0311 | 0.1482 | 0.8352 | 0.9987 |
| R SPC | R PAC | PV | 0.3041 | 0.1566 | 0.0608 | 0.6056 |
| R SPC | R PAC | RD | 0.0805 | 0.1596 | 0.6171 | 0.9964 |
| R TP | R PAC | AD | 0.0819 | 0.1517 | 0.5931 | 0.9951 |
| R TP | R PAC | FA | 0.1125 | 0.1581 | 0.4818 | 0.9215 |
| R TP | R PAC | fNT | 0.3259 | 0.1764 | 0.0749 | 0.4089 |
| R TP | R PAC | MD | 0.1241 | 0.1339 | 0.3610 | 0.9987 |
| R TP | R PAC | PV | 0.1542 | 0.1713 | 0.3749 | 0.9083 |
| R TP | R PAC | RD | 0.1170 | 0.1399 | 0.4093 | 0.9964 |
| L Caudate | R CN/SOC | AD | 0.0332 | 0.1790 | 0.8540 | 0.9951 |
| L Caudate | R CN/SOC | FA | 0.1543 | 0.1753 | 0.3858 | 0.9215 |
| L Caudate | R CN/SOC | fNT | 0.2444 | 0.1770 | 0.1776 | 0.6503 |
| L Caudate | R CN/SOC | MD | 0.1059 | 0.1737 | 0.5467 | 0.9987 |
| L Caudate | R CN/SOC | PV | 0.1138 | 0.1814 | 0.5353 | 0.9192 |
| L Caudate | R CN/SOC | RD | 0.1790 | 0.1661 | 0.2896 | 0.9964 |
| L Pallidum | R CN/SOC | AD | 0.0828 | 0.1761 | 0.6415 | 0.9951 |
| L Pallidum | R CN/SOC | FA | 0.1811 | 0.1718 | 0.2998 | 0.9215 |
| L Pallidum | R CN/SOC | fNT | 0.1550 | 0.1736 | 0.3788 | 0.7794 |
| L Pallidum | R CN/SOC | MD | 0.0277 | 0.1757 | 0.8759 | 0.9987 |
| L Pallidum | R CN/SOC | PV | 0.0476 | 0.1688 | 0.7797 | 0.9629 |
| L Pallidum | R CN/SOC | RD | 0.0379 | 0.1744 | 0.8292 | 0.9964 |
| L Putamen | R CN/SOC | AD | 0.0569 | 0.1739 | 0.7454 | 0.9951 |
| L Putamen | R CN/SOC | FA | 0.1233 | 0.1692 | 0.4715 | 0.9215 |
| L Putamen | R CN/SOC | fNT | 0.1268 | 0.1710 | 0.4635 | 0.8071 |
| L Putamen | R CN/SOC | MD | 0.1012 | 0.1726 | 0.5616 | 0.9987 |
| L Putamen | R CN/SOC | PV | 0.0147 | 0.1720 | 0.9324 | 0.9984 |
| L Putamen | R CN/SOC | RD | 0.1358 | 0.1705 | 0.4316 | 0.9964 |
| L SFC | R CN/SOC | AD | 0.0979 | 0.1694 | 0.5673 | 0.9951 |
| L SFC | R CN/SOC | FA | 0.0831 | 0.1665 | 0.6210 | 0.9260 |
| L SFC | R CN/SOC | fNT | 0.1444 | 0.1709 | 0.4043 | 0.7875 |
| L SFC | R CN/SOC | MD | 0.0731 | 0.1636 | 0.6577 | 0.9987 |
| L SFC | R CN/SOC | PV | 0.0156 | 0.1716 | 0.9281 | 0.9984 |
| L SFC | R CN/SOC | RD | 0.0298 | 0.1605 | 0.8537 | 0.9964 |
| R Amygdala | R CN/SOC | AD | 0.1580 | 0.1741 | 0.3710 | 0.9951 |
| R Amygdala | R CN/SOC | FA | 0.2211 | 0.1697 | 0.2023 | 0.9215 |
| R Amygdala | R CN/SOC | fNT | 0.0225 | 0.1694 | 0.8954 | 0.9789 |
| R Amygdala | R CN/SOC | MD | 0.0389 | 0.1775 | 0.8279 | 0.9987 |
| R Amygdala | R CN/SOC | PV | 0.3044 | 0.1593 | 0.0652 | 0.6248 |
| R Amygdala | R CN/SOC | RD | 0.1023 | 0.1753 | 0.5636 | 0.9964 |
| R Anterior | R CN/SOC | AD | 0.0367 | 0.1735 | 0.8336 | 0.9951 |
| R Anterior | R CN/SOC | FA | 0.0836 | 0.1720 | 0.6303 | 0.9260 |
| R Anterior | R CN/SOC | fNT | 0.1979 | 0.1668 | 0.2439 | 0.6924 |
| R Anterior | R CN/SOC | MD | 0.0248 | 0.1735 | 0.8872 | 0.9987 |
| R Anterior | R CN/SOC | PV | 0.1425 | 0.1725 | 0.4146 | 0.9083 |
| R Anterior | R CN/SOC | RD | 0.0842 | 0.1725 | 0.6288 | 0.9964 |
| R Caudate | R CN/SOC | AD | 0.0748 | 0.1725 | 0.6676 | 0.9951 |
| R Caudate | R CN/SOC | FA | 0.0336 | 0.1762 | 0.8498 | 0.9772 |
| R Caudate | R CN/SOC | fNT | 0.3345 | 0.1734 | 0.0642 | 0.3950 |
| R Caudate | R CN/SOC | MD | 0.0705 | 0.1681 | 0.6780 | 0.9987 |
| R Caudate | R CN/SOC | PV | 0.0002 | 0.1719 | 0.9993 | 0.9996 |
| R Caudate | R CN/SOC | RD | 0.0370 | 0.1689 | 0.8280 | 0.9964 |
| R Medio_Dorsal | R CN/SOC | AD | 0.0137 | 0.1762 | 0.9384 | 0.9951 |
| R Medio_Dorsal | R CN/SOC | FA | 0.0738 | 0.1757 | 0.6772 | 0.9395 |
| R Medio_Dorsal | R CN/SOC | fNT | 0.0989 | 0.1413 | 0.4891 | 0.8342 |
| R Medio_Dorsal | R CN/SOC | MD | 0.0781 | 0.1754 | 0.6591 | 0.9987 |
| R Medio_Dorsal | R CN/SOC | PV | 0.2875 | 0.1665 | 0.0938 | 0.7648 |
| R Medio_Dorsal | R CN/SOC | RD | 0.1191 | 0.1746 | 0.5000 | 0.9964 |
| R Pallidum | R CN/SOC | AD | 0.0028 | 0.1741 | 0.9875 | 0.9951 |
| R Pallidum | R CN/SOC | FA | 0.1517 | 0.1702 | 0.3791 | 0.9215 |
| R Pallidum | R CN/SOC | fNT | 0.0960 | 0.1556 | 0.5414 | 0.8670 |
| R Pallidum | R CN/SOC | MD | 0.0623 | 0.1739 | 0.7226 | 0.9987 |
| R Pallidum | R CN/SOC | PV | 0.1616 | 0.1686 | 0.3449 | 0.9039 |
| R Pallidum | R CN/SOC | RD | 0.1080 | 0.1727 | 0.5358 | 0.9964 |
| R Putamen | R CN/SOC | AD | 0.0694 | 0.1763 | 0.6963 | 0.9951 |
| R Putamen | R CN/SOC | FA | 0.0617 | 0.1712 | 0.7209 | 0.9395 |
| R Putamen | R CN/SOC | fNT | 0.1368 | 0.1627 | 0.4067 | 0.7880 |
| R Putamen | R CN/SOC | MD | 0.0414 | 0.1751 | 0.8147 | 0.9987 |
| R Putamen | R CN/SOC | PV | 0.2400 | 0.1707 | 0.1695 | 0.9039 |
| R Putamen | R CN/SOC | RD | 0.0044 | 0.1728 | 0.9797 | 0.9964 |
| R rMFC | R CN/SOC | AD | 0.0839 | 0.1692 | 0.6235 | 0.9951 |
| R rMFC | R CN/SOC | FA | 0.0008 | 0.1720 | 0.9965 | 0.9998 |
| R rMFC | R CN/SOC | fNT | 0.0624 | 0.1737 | 0.7216 | 0.9189 |
| R rMFC | R CN/SOC | MD | 0.0869 | 0.1683 | 0.6091 | 0.9987 |
| R rMFC | R CN/SOC | PV | 0.0837 | 0.1740 | 0.6339 | 0.9566 |
| R rMFC | R CN/SOC | RD | 0.0671 | 0.1691 | 0.6941 | 0.9964 |
| R SFC | R CN/SOC | AD | 0.0916 | 0.1687 | 0.5909 | 0.9951 |
| R SFC | R CN/SOC | FA | 0.0923 | 0.1638 | 0.5768 | 0.9215 |
| R SFC | R CN/SOC | fNT | 0.0383 | 0.1733 | 0.8265 | 0.9484 |
| R SFC | R CN/SOC | MD | 0.0388 | 0.1676 | 0.8182 | 0.9987 |
| R SFC | R CN/SOC | PV | 0.0901 | 0.1663 | 0.5916 | 0.9566 |
| R SFC | R CN/SOC | RD | 0.0217 | 0.1636 | 0.8954 | 0.9964 |
| R SPC | R CN/SOC | AD | 0.0084 | 0.1741 | 0.9618 | 0.9951 |
| R SPC | R CN/SOC | FA | 0.2555 | 0.1688 | 0.1404 | 0.8632 |
| R SPC | R CN/SOC | fNT | 0.0764 | 0.1715 | 0.6592 | 0.8866 |
| R SPC | R CN/SOC | MD | 0.1582 | 0.1688 | 0.3559 | 0.9987 |
| R SPC | R CN/SOC | PV | 0.0783 | 0.1780 | 0.6631 | 0.9629 |
| R SPC | R CN/SOC | RD | 0.2418 | 0.1659 | 0.1550 | 0.9964 |

## Appendix C: DTI-based structural network

**Table 5:**
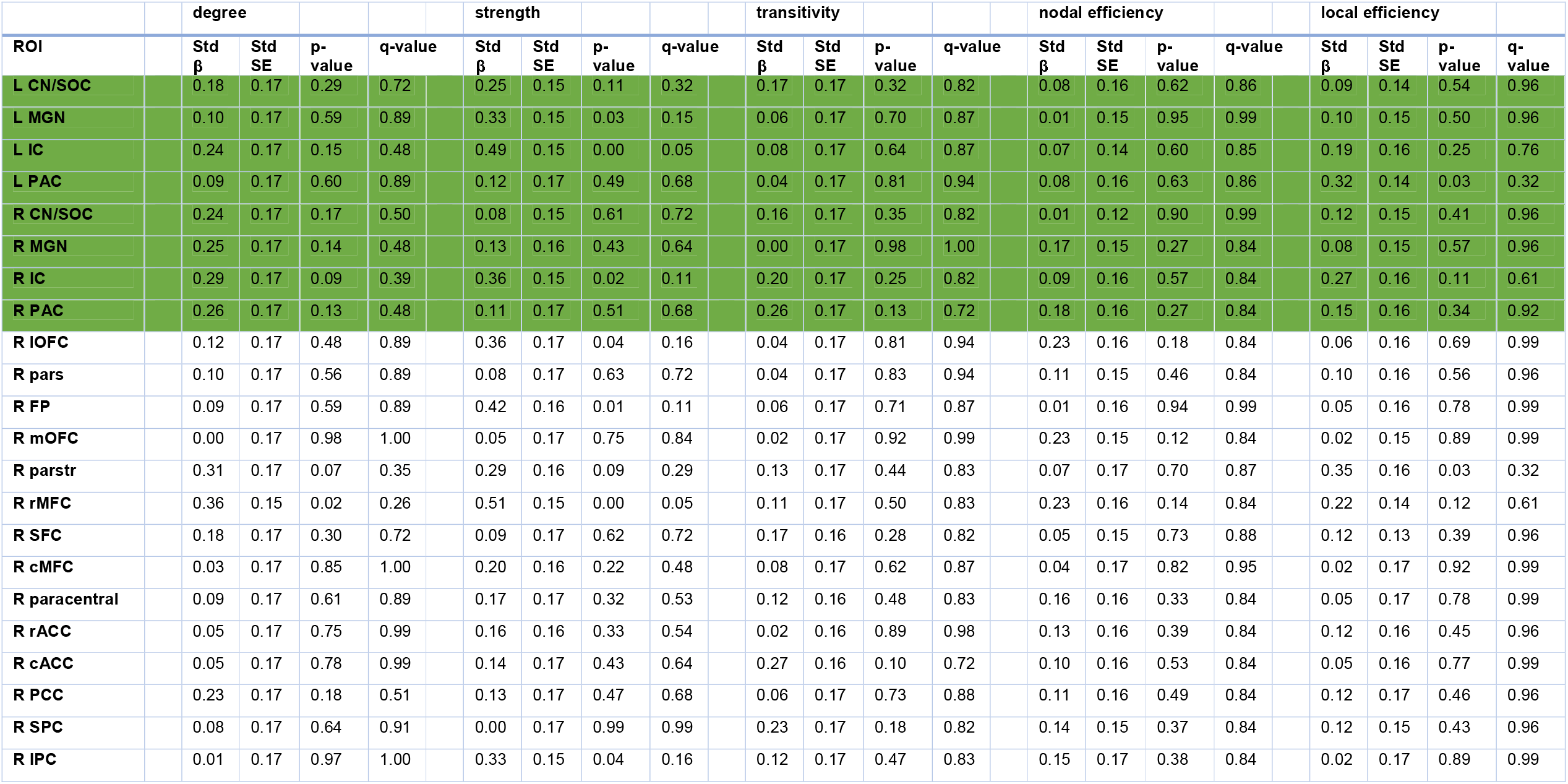
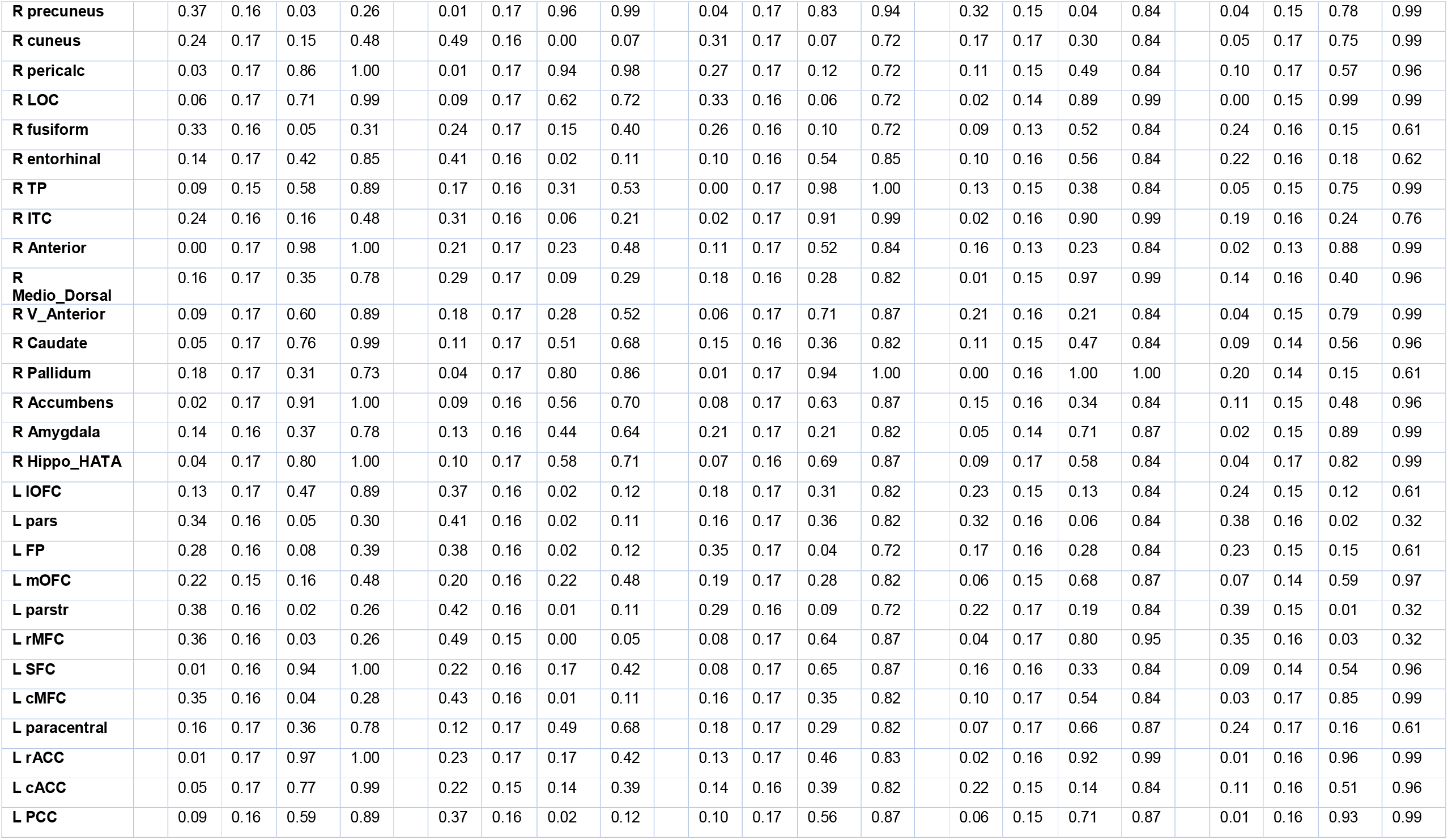
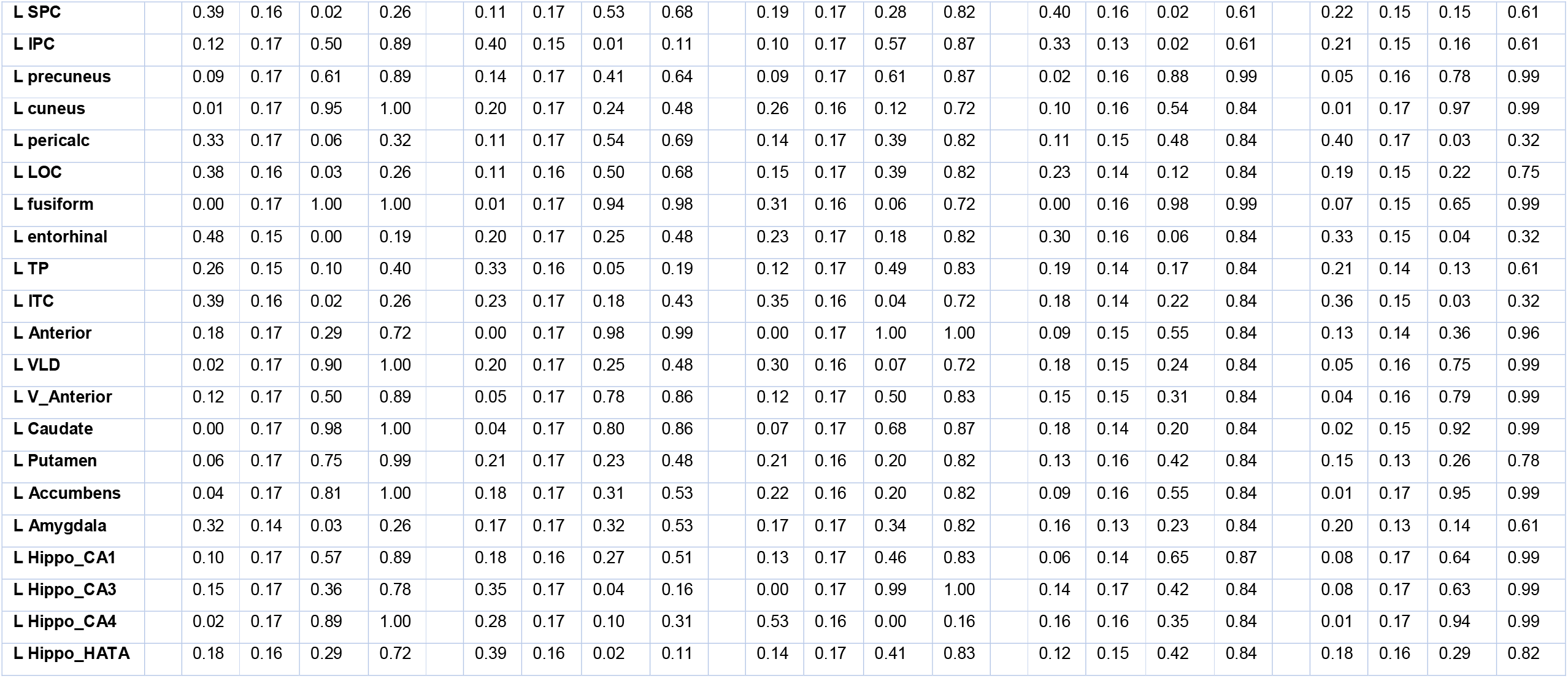
Standardized Beta (Std β), Standard Error (Std SE) values, p values and FDR corrected q values for the comparison between CHUU and CPHIV over the nodal graph measures degree, strength, transitivity, nodal efficiency and local efficiency. CAS regions are highlighted in green.

|  | degree |  |  |  | strength |  |  |  | transitivity |  |  |  | nodal efficiency |  |  |  | local efficiency |  |  |  |
| --- | --- | --- | --- | --- | --- | --- | --- | --- | --- | --- | --- | --- | --- | --- | --- | --- | --- | --- | --- | --- |
| ROI | Std $\beta$ | Std SE | p-value | q-value | Std $\beta$ | Std SE | p-value | q-value | Std $\beta$ | Std SE | p-value | q-value | Std $\beta$ | Std SE | p-value | q-value | Std $\beta$ | Std SE | p-value | q-value |
| L CN/SOC | 0.18 | 0.17 | 0.29 | 0.72 | 0.25 | 0.15 | 0.11 | 0.32 | 0.17 | 0.17 | 0.32 | 0.82 | 0.08 | 0.16 | 0.62 | 0.86 | 0.09 | 0.14 | 0.54 | 0.96 |
| L MGN | 0.10 | 0.17 | 0.59 | 0.89 | 0.33 | 0.15 | 0.03 | 0.15 | 0.06 | 0.17 | 0.70 | 0.87 | 0.01 | 0.15 | 0.95 | 0.99 | 0.10 | 0.15 | 0.50 | 0.96 |
| L IC | 0.24 | 0.17 | 0.15 | 0.48 | 0.49 | 0.15 | 0.00 | 0.05 | 0.08 | 0.17 | 0.64 | 0.87 | 0.07 | 0.14 | 0.60 | 0.85 | 0.19 | 0.16 | 0.25 | 0.76 |
| L PAC | 0.09 | 0.17 | 0.60 | 0.89 | 0.12 | 0.17 | 0.49 | 0.68 | 0.04 | 0.17 | 0.81 | 0.94 | 0.08 | 0.16 | 0.63 | 0.86 | 0.32 | 0.14 | 0.03 | 0.32 |
| R CN/SOC | 0.24 | 0.17 | 0.17 | 0.50 | 0.08 | 0.15 | 0.61 | 0.72 | 0.16 | 0.17 | 0.35 | 0.82 | 0.01 | 0.12 | 0.90 | 0.99 | 0.12 | 0.15 | 0.41 | 0.96 |
| R MGN | 0.25 | 0.17 | 0.14 | 0.48 | 0.13 | 0.16 | 0.43 | 0.64 | 0.00 | 0.17 | 0.98 | 1.00 | 0.17 | 0.15 | 0.27 | 0.84 | 0.08 | 0.15 | 0.57 | 0.96 |
| R IC | 0.29 | 0.17 | 0.09 | 0.39 | 0.36 | 0.15 | 0.02 | 0.11 | 0.20 | 0.17 | 0.25 | 0.82 | 0.09 | 0.16 | 0.57 | 0.84 | 0.27 | 0.16 | 0.11 | 0.61 |
| R PAC | 0.26 | 0.17 | 0.13 | 0.48 | 0.11 | 0.17 | 0.51 | 0.68 | 0.26 | 0.17 | 0.13 | 0.72 | 0.18 | 0.16 | 0.27 | 0.84 | 0.15 | 0.16 | 0.34 | 0.92 |
| R IOFC | 0.12 | 0.17 | 0.48 | 0.89 | 0.36 | 0.17 | 0.04 | 0.16 | 0.04 | 0.17 | 0.81 | 0.94 | 0.23 | 0.16 | 0.18 | 0.84 | 0.06 | 0.16 | 0.69 | 0.99 |
| R pars | 0.10 | 0.17 | 0.56 | 0.89 | 0.08 | 0.17 | 0.63 | 0.72 | 0.04 | 0.17 | 0.83 | 0.94 | 0.11 | 0.15 | 0.46 | 0.84 | 0.10 | 0.16 | 0.56 | 0.96 |
| R FP | 0.09 | 0.17 | 0.59 | 0.89 | 0.42 | 0.16 | 0.01 | 0.11 | 0.06 | 0.17 | 0.71 | 0.87 | 0.01 | 0.16 | 0.94 | 0.99 | 0.05 | 0.16 | 0.78 | 0.99 |
| R mOFC | 0.00 | 0.17 | 0.98 | 1.00 | 0.05 | 0.17 | 0.75 | 0.84 | 0.02 | 0.17 | 0.92 | 0.99 | 0.23 | 0.15 | 0.12 | 0.84 | 0.02 | 0.15 | 0.89 | 0.99 |
| R parstr | 0.31 | 0.17 | 0.07 | 0.35 | 0.29 | 0.16 | 0.09 | 0.29 | 0.13 | 0.17 | 0.44 | 0.83 | 0.07 | 0.17 | 0.70 | 0.87 | 0.35 | 0.16 | 0.03 | 0.32 |
| R rMFC | 0.36 | 0.15 | 0.02 | 0.26 | 0.51 | 0.15 | 0.00 | 0.05 | 0.11 | 0.17 | 0.50 | 0.83 | 0.23 | 0.16 | 0.14 | 0.84 | 0.22 | 0.14 | 0.12 | 0.61 |
| R SFC | 0.18 | 0.17 | 0.30 | 0.72 | 0.09 | 0.17 | 0.62 | 0.72 | 0.17 | 0.16 | 0.28 | 0.82 | 0.05 | 0.15 | 0.73 | 0.88 | 0.12 | 0.13 | 0.39 | 0.96 |
| R cMFC | 0.03 | 0.17 | 0.85 | 1.00 | 0.20 | 0.16 | 0.22 | 0.48 | 0.08 | 0.17 | 0.62 | 0.87 | 0.04 | 0.17 | 0.82 | 0.95 | 0.02 | 0.17 | 0.92 | 0.99 |
| R paracentral | 0.09 | 0.17 | 0.61 | 0.89 | 0.17 | 0.17 | 0.32 | 0.53 | 0.12 | 0.16 | 0.48 | 0.83 | 0.16 | 0.16 | 0.33 | 0.84 | 0.05 | 0.17 | 0.78 | 0.99 |
| R rACC | 0.05 | 0.17 | 0.75 | 0.99 | 0.16 | 0.16 | 0.33 | 0.54 | 0.02 | 0.16 | 0.89 | 0.98 | 0.13 | 0.16 | 0.39 | 0.84 | 0.12 | 0.16 | 0.45 | 0.96 |
| R cACC | 0.05 | 0.17 | 0.78 | 0.99 | 0.14 | 0.17 | 0.43 | 0.64 | 0.27 | 0.16 | 0.10 | 0.72 | 0.10 | 0.16 | 0.53 | 0.84 | 0.05 | 0.16 | 0.77 | 0.99 |
| R PCC | 0.23 | 0.17 | 0.18 | 0.51 | 0.13 | 0.17 | 0.47 | 0.68 | 0.06 | 0.17 | 0.73 | 0.88 | 0.11 | 0.16 | 0.49 | 0.84 | 0.12 | 0.17 | 0.46 | 0.96 |
| R SPC | 0.08 | 0.17 | 0.64 | 0.91 | 0.00 | 0.17 | 0.99 | 0.99 | 0.23 | 0.17 | 0.18 | 0.82 | 0.14 | 0.15 | 0.37 | 0.84 | 0.12 | 0.15 | 0.43 | 0.96 |
| R IPC | 0.01 | 0.17 | 0.97 | 1.00 | 0.33 | 0.15 | 0.04 | 0.16 | 0.12 | 0.17 | 0.47 | 0.83 | 0.15 | 0.17 | 0.38 | 0.84 | 0.02 | 0.17 | 0.89 | 0.99 |

|  |  |  |  |  |  |  |  |  |  |  |  |  |  |  |  |  |  |  |  |  |  |  |  |  |  |
| --- | --- | --- | --- | --- | --- | --- | --- | --- | --- | --- | --- | --- | --- | --- | --- | --- | --- | --- | --- | --- | --- | --- | --- | --- | --- |
| R precuneus |  | 0.37 | 0.16 | 0.03 | 0.26 |  | 0.01 | 0.17 | 0.96 | 0.99 |  | 0.04 | 0.17 | 0.83 | 0.94 |  | 0.32 | 0.15 | 0.04 | 0.84 |  | 0.04 | 0.15 | 0.78 | 0.99 |
| R cuneus |  | 0.24 | 0.17 | 0.15 | 0.48 |  | 0.49 | 0.16 | 0.00 | 0.07 |  | 0.31 | 0.17 | 0.07 | 0.72 |  | 0.17 | 0.17 | 0.30 | 0.84 |  | 0.05 | 0.17 | 0.75 | 0.99 |
| R pericalc |  | 0.03 | 0.17 | 0.86 | 1.00 |  | 0.01 | 0.17 | 0.94 | 0.98 |  | 0.27 | 0.17 | 0.12 | 0.72 |  | 0.11 | 0.15 | 0.49 | 0.84 |  | 0.10 | 0.17 | 0.57 | 0.96 |
| R LOC |  | 0.06 | 0.17 | 0.71 | 0.99 |  | 0.09 | 0.17 | 0.62 | 0.72 |  | 0.33 | 0.16 | 0.06 | 0.72 |  | 0.02 | 0.14 | 0.89 | 0.99 |  | 0.00 | 0.15 | 0.99 | 0.99 |
| R fusiform |  | 0.33 | 0.16 | 0.05 | 0.31 |  | 0.24 | 0.17 | 0.15 | 0.40 |  | 0.26 | 0.16 | 0.10 | 0.72 |  | 0.09 | 0.13 | 0.52 | 0.84 |  | 0.24 | 0.16 | 0.15 | 0.61 |
| R entorhinal |  | 0.14 | 0.17 | 0.42 | 0.85 |  | 0.41 | 0.16 | 0.02 | 0.11 |  | 0.10 | 0.16 | 0.54 | 0.85 |  | 0.10 | 0.16 | 0.56 | 0.84 |  | 0.22 | 0.16 | 0.18 | 0.62 |
| R TP |  | 0.09 | 0.15 | 0.58 | 0.89 |  | 0.17 | 0.16 | 0.31 | 0.53 |  | 0.00 | 0.17 | 0.98 | 1.00 |  | 0.13 | 0.15 | 0.38 | 0.84 |  | 0.05 | 0.15 | 0.75 | 0.99 |
| R ITC |  | 0.24 | 0.16 | 0.16 | 0.48 |  | 0.31 | 0.16 | 0.06 | 0.21 |  | 0.02 | 0.17 | 0.91 | 0.99 |  | 0.02 | 0.16 | 0.90 | 0.99 |  | 0.19 | 0.16 | 0.24 | 0.76 |
| R Anterior |  | 0.00 | 0.17 | 0.98 | 1.00 |  | 0.21 | 0.17 | 0.23 | 0.48 |  | 0.11 | 0.17 | 0.52 | 0.84 |  | 0.16 | 0.13 | 0.23 | 0.84 |  | 0.02 | 0.13 | 0.88 | 0.99 |
| R Medio_Dorsal |  | 0.16 | 0.17 | 0.35 | 0.78 |  | 0.29 | 0.17 | 0.09 | 0.29 |  | 0.18 | 0.16 | 0.28 | 0.82 |  | 0.01 | 0.15 | 0.97 | 0.99 |  | 0.14 | 0.16 | 0.40 | 0.96 |
| R V_Anterior |  | 0.09 | 0.17 | 0.60 | 0.89 |  | 0.18 | 0.17 | 0.28 | 0.52 |  | 0.06 | 0.17 | 0.71 | 0.87 |  | 0.21 | 0.16 | 0.21 | 0.84 |  | 0.04 | 0.15 | 0.79 | 0.99 |
| R Caudate |  | 0.05 | 0.17 | 0.76 | 0.99 |  | 0.11 | 0.17 | 0.51 | 0.68 |  | 0.15 | 0.16 | 0.36 | 0.82 |  | 0.11 | 0.15 | 0.47 | 0.84 |  | 0.09 | 0.14 | 0.56 | 0.96 |
| R Pallidum |  | 0.18 | 0.17 | 0.31 | 0.73 |  | 0.04 | 0.17 | 0.80 | 0.86 |  | 0.01 | 0.17 | 0.94 | 1.00 |  | 0.00 | 0.16 | 1.00 | 1.00 |  | 0.20 | 0.14 | 0.15 | 0.61 |
| R Accumbens |  | 0.02 | 0.17 | 0.91 | 1.00 |  | 0.09 | 0.16 | 0.56 | 0.70 |  | 0.08 | 0.17 | 0.63 | 0.87 |  | 0.15 | 0.16 | 0.34 | 0.84 |  | 0.11 | 0.15 | 0.48 | 0.96 |
| R Amygdala |  | 0.14 | 0.16 | 0.37 | 0.78 |  | 0.13 | 0.16 | 0.44 | 0.64 |  | 0.21 | 0.17 | 0.21 | 0.82 |  | 0.05 | 0.14 | 0.71 | 0.87 |  | 0.02 | 0.15 | 0.89 | 0.99 |
| R Hippo_HATA |  | 0.04 | 0.17 | 0.80 | 1.00 |  | 0.10 | 0.17 | 0.58 | 0.71 |  | 0.07 | 0.16 | 0.69 | 0.87 |  | 0.09 | 0.17 | 0.58 | 0.84 |  | 0.04 | 0.17 | 0.82 | 0.99 |
| L IOFC |  | 0.13 | 0.17 | 0.47 | 0.89 |  | 0.37 | 0.16 | 0.02 | 0.12 |  | 0.18 | 0.17 | 0.31 | 0.82 |  | 0.23 | 0.15 | 0.13 | 0.84 |  | 0.24 | 0.15 | 0.12 | 0.61 |
| L pars |  | 0.34 | 0.16 | 0.05 | 0.30 |  | 0.41 | 0.16 | 0.02 | 0.11 |  | 0.16 | 0.17 | 0.36 | 0.82 |  | 0.32 | 0.16 | 0.06 | 0.84 |  | 0.38 | 0.16 | 0.02 | 0.32 |
| L FP |  | 0.28 | 0.16 | 0.08 | 0.39 |  | 0.38 | 0.16 | 0.02 | 0.12 |  | 0.35 | 0.17 | 0.04 | 0.72 |  | 0.17 | 0.16 | 0.28 | 0.84 |  | 0.23 | 0.15 | 0.15 | 0.61 |
| L mOFC |  | 0.22 | 0.15 | 0.16 | 0.48 |  | 0.20 | 0.16 | 0.22 | 0.48 |  | 0.19 | 0.17 | 0.28 | 0.82 |  | 0.06 | 0.15 | 0.68 | 0.87 |  | 0.07 | 0.14 | 0.59 | 0.97 |
| L parstr |  | 0.38 | 0.16 | 0.02 | 0.26 |  | 0.42 | 0.16 | 0.01 | 0.11 |  | 0.29 | 0.16 | 0.09 | 0.72 |  | 0.22 | 0.17 | 0.19 | 0.84 |  | 0.39 | 0.15 | 0.01 | 0.32 |
| L rMFC |  | 0.36 | 0.16 | 0.03 | 0.26 |  | 0.49 | 0.15 | 0.00 | 0.05 |  | 0.08 | 0.17 | 0.64 | 0.87 |  | 0.04 | 0.17 | 0.80 | 0.95 |  | 0.35 | 0.16 | 0.03 | 0.32 |
| L SFC |  | 0.01 | 0.16 | 0.94 | 1.00 |  | 0.22 | 0.16 | 0.17 | 0.42 |  | 0.08 | 0.17 | 0.65 | 0.87 |  | 0.16 | 0.16 | 0.33 | 0.84 |  | 0.09 | 0.14 | 0.54 | 0.96 |
| L cMFC |  | 0.35 | 0.16 | 0.04 | 0.28 |  | 0.43 | 0.16 | 0.01 | 0.11 |  | 0.16 | 0.17 | 0.35 | 0.82 |  | 0.10 | 0.17 | 0.54 | 0.84 |  | 0.03 | 0.17 | 0.85 | 0.99 |
| L paracentral |  | 0.16 | 0.17 | 0.36 | 0.78 |  | 0.12 | 0.17 | 0.49 | 0.68 |  | 0.18 | 0.17 | 0.29 | 0.82 |  | 0.07 | 0.17 | 0.66 | 0.87 |  | 0.24 | 0.17 | 0.16 | 0.61 |
| L rACC |  | 0.01 | 0.17 | 0.97 | 1.00 |  | 0.23 | 0.17 | 0.17 | 0.42 |  | 0.13 | 0.17 | 0.46 | 0.83 |  | 0.02 | 0.16 | 0.92 | 0.99 |  | 0.01 | 0.16 | 0.96 | 0.99 |
| L cACC |  | 0.05 | 0.17 | 0.77 | 0.99 |  | 0.22 | 0.15 | 0.14 | 0.39 |  | 0.14 | 0.16 | 0.39 | 0.82 |  | 0.22 | 0.15 | 0.14 | 0.84 |  | 0.11 | 0.16 | 0.51 | 0.96 |
| L PCC |  | 0.09 | 0.16 | 0.59 | 0.89 |  | 0.37 | 0.16 | 0.02 | 0.12 |  | 0.10 | 0.17 | 0.56 | 0.87 |  | 0.06 | 0.15 | 0.71 | 0.87 |  | 0.01 | 0.16 | 0.93 | 0.99 |
| L SPC |  | 0.39 | 0.16 | 0.02 | 0.26 |  | 0.11 | 0.17 | 0.53 | 0.68 |  | 0.19 | 0.17 | 0.28 | 0.82 |  | 0.40 | 0.16 | 0.02 | 0.61 |  | 0.22 | 0.15 | 0.15 | 0.61 |
| L IPC |  | 0.12 | 0.17 | 0.50 | 0.89 |  | 0.40 | 0.15 | 0.01 | 0.11 |  | 0.10 | 0.17 | 0.57 | 0.87 |  | 0.33 | 0.13 | 0.02 | 0.61 |  | 0.21 | 0.15 | 0.16 | 0.61 |
| L precuneus |  | 0.09 | 0.17 | 0.61 | 0.89 |  | 0.14 | 0.17 | 0.41 | 0.64 |  | 0.09 | 0.17 | 0.61 | 0.87 |  | 0.02 | 0.16 | 0.88 | 0.99 |  | 0.05 | 0.16 | 0.78 | 0.99 |
| L cuneus |  | 0.01 | 0.17 | 0.95 | 1.00 |  | 0.20 | 0.17 | 0.24 | 0.48 |  | 0.26 | 0.16 | 0.12 | 0.72 |  | 0.10 | 0.16 | 0.54 | 0.84 |  | 0.01 | 0.17 | 0.97 | 0.99 |
| L pericalc |  | 0.33 | 0.17 | 0.06 | 0.32 |  | 0.11 | 0.17 | 0.54 | 0.69 |  | 0.14 | 0.17 | 0.39 | 0.82 |  | 0.11 | 0.15 | 0.48 | 0.84 |  | 0.40 | 0.17 | 0.03 | 0.32 |
| L LOC |  | 0.38 | 0.16 | 0.03 | 0.26 |  | 0.11 | 0.16 | 0.50 | 0.68 |  | 0.15 | 0.17 | 0.39 | 0.82 |  | 0.23 | 0.14 | 0.12 | 0.84 |  | 0.19 | 0.15 | 0.22 | 0.75 |
| L fusiform |  | 0.00 | 0.17 | 1.00 | 1.00 |  | 0.01 | 0.17 | 0.94 | 0.98 |  | 0.31 | 0.16 | 0.06 | 0.72 |  | 0.00 | 0.16 | 0.98 | 0.99 |  | 0.07 | 0.15 | 0.65 | 0.99 |
| L entorhinal |  | 0.48 | 0.15 | 0.00 | 0.19 |  | 0.20 | 0.17 | 0.25 | 0.48 |  | 0.23 | 0.17 | 0.18 | 0.82 |  | 0.30 | 0.16 | 0.06 | 0.84 |  | 0.33 | 0.15 | 0.04 | 0.32 |
| L TP |  | 0.26 | 0.15 | 0.10 | 0.40 |  | 0.33 | 0.16 | 0.05 | 0.19 |  | 0.12 | 0.17 | 0.49 | 0.83 |  | 0.19 | 0.14 | 0.17 | 0.84 |  | 0.21 | 0.14 | 0.13 | 0.61 |
| L ITC |  | 0.39 | 0.16 | 0.02 | 0.26 |  | 0.23 | 0.17 | 0.18 | 0.43 |  | 0.35 | 0.16 | 0.04 | 0.72 |  | 0.18 | 0.14 | 0.22 | 0.84 |  | 0.36 | 0.15 | 0.03 | 0.32 |
| L Anterior |  | 0.18 | 0.17 | 0.29 | 0.72 |  | 0.00 | 0.17 | 0.98 | 0.99 |  | 0.00 | 0.17 | 1.00 | 1.00 |  | 0.09 | 0.15 | 0.55 | 0.84 |  | 0.13 | 0.14 | 0.36 | 0.96 |
| L VLD |  | 0.02 | 0.17 | 0.90 | 1.00 |  | 0.20 | 0.17 | 0.25 | 0.48 |  | 0.30 | 0.16 | 0.07 | 0.72 |  | 0.18 | 0.15 | 0.24 | 0.84 |  | 0.05 | 0.16 | 0.75 | 0.99 |
| L V_Anterior |  | 0.12 | 0.17 | 0.50 | 0.89 |  | 0.05 | 0.17 | 0.78 | 0.86 |  | 0.12 | 0.17 | 0.50 | 0.83 |  | 0.15 | 0.15 | 0.31 | 0.84 |  | 0.04 | 0.16 | 0.79 | 0.99 |
| L Caudate |  | 0.00 | 0.17 | 0.98 | 1.00 |  | 0.04 | 0.17 | 0.80 | 0.86 |  | 0.07 | 0.17 | 0.68 | 0.87 |  | 0.18 | 0.14 | 0.20 | 0.84 |  | 0.02 | 0.15 | 0.92 | 0.99 |
| L Putamen |  | 0.06 | 0.17 | 0.75 | 0.99 |  | 0.21 | 0.17 | 0.23 | 0.48 |  | 0.21 | 0.16 | 0.20 | 0.82 |  | 0.13 | 0.16 | 0.42 | 0.84 |  | 0.15 | 0.13 | 0.26 | 0.78 |
| L Accumbens |  | 0.04 | 0.17 | 0.81 | 1.00 |  | 0.18 | 0.17 | 0.31 | 0.53 |  | 0.22 | 0.16 | 0.20 | 0.82 |  | 0.09 | 0.16 | 0.55 | 0.84 |  | 0.01 | 0.17 | 0.95 | 0.99 |
| L Amygdala |  | 0.32 | 0.14 | 0.03 | 0.26 |  | 0.17 | 0.17 | 0.32 | 0.53 |  | 0.17 | 0.17 | 0.34 | 0.82 |  | 0.16 | 0.13 | 0.23 | 0.84 |  | 0.20 | 0.13 | 0.14 | 0.61 |
| L Hippo_CA1 |  | 0.10 | 0.17 | 0.57 | 0.89 |  | 0.18 | 0.16 | 0.27 | 0.51 |  | 0.13 | 0.17 | 0.46 | 0.83 |  | 0.06 | 0.14 | 0.65 | 0.87 |  | 0.08 | 0.17 | 0.64 | 0.99 |
| L Hippo_CA3 |  | 0.15 | 0.17 | 0.36 | 0.78 |  | 0.35 | 0.17 | 0.04 | 0.16 |  | 0.00 | 0.17 | 0.99 | 1.00 |  | 0.14 | 0.17 | 0.42 | 0.84 |  | 0.08 | 0.17 | 0.63 | 0.99 |
| L Hippo_CA4 |  | 0.02 | 0.17 | 0.89 | 1.00 |  | 0.28 | 0.17 | 0.10 | 0.31 |  | 0.53 | 0.16 | 0.00 | 0.16 |  | 0.16 | 0.16 | 0.35 | 0.84 |  | 0.01 | 0.17 | 0.94 | 0.99 |
| L Hippo_HATA |  | 0.18 | 0.16 | 0.29 | 0.72 |  | 0.39 | 0.16 | 0.02 | 0.11 |  | 0.14 | 0.17 | 0.41 | 0.83 |  | 0.12 | 0.15 | 0.42 | 0.84 |  | 0.18 | 0.16 | 0.29 | 0.82 |

## Appendix D: RS-fMRI Functional Connectivity

**Table 6:**
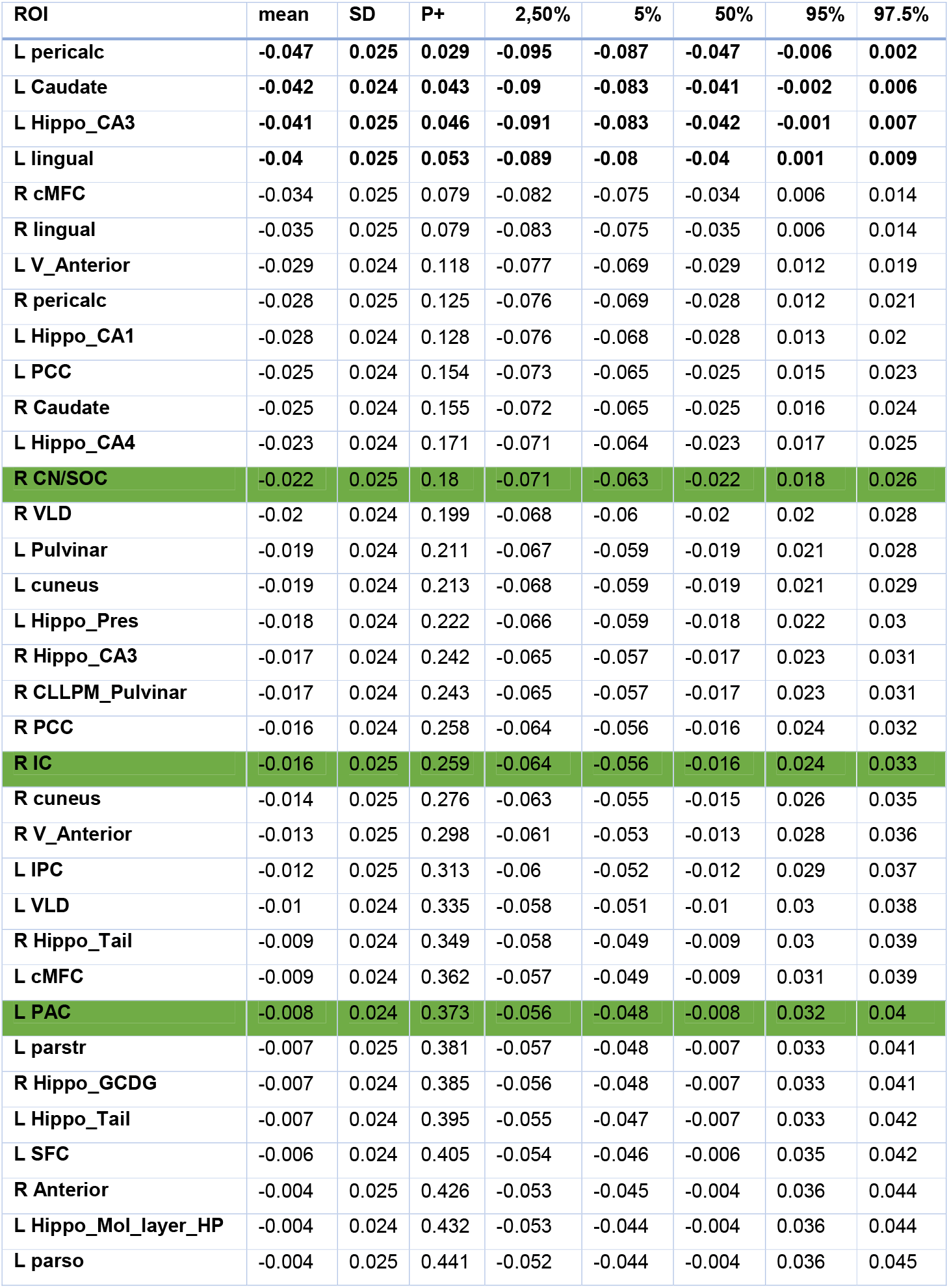
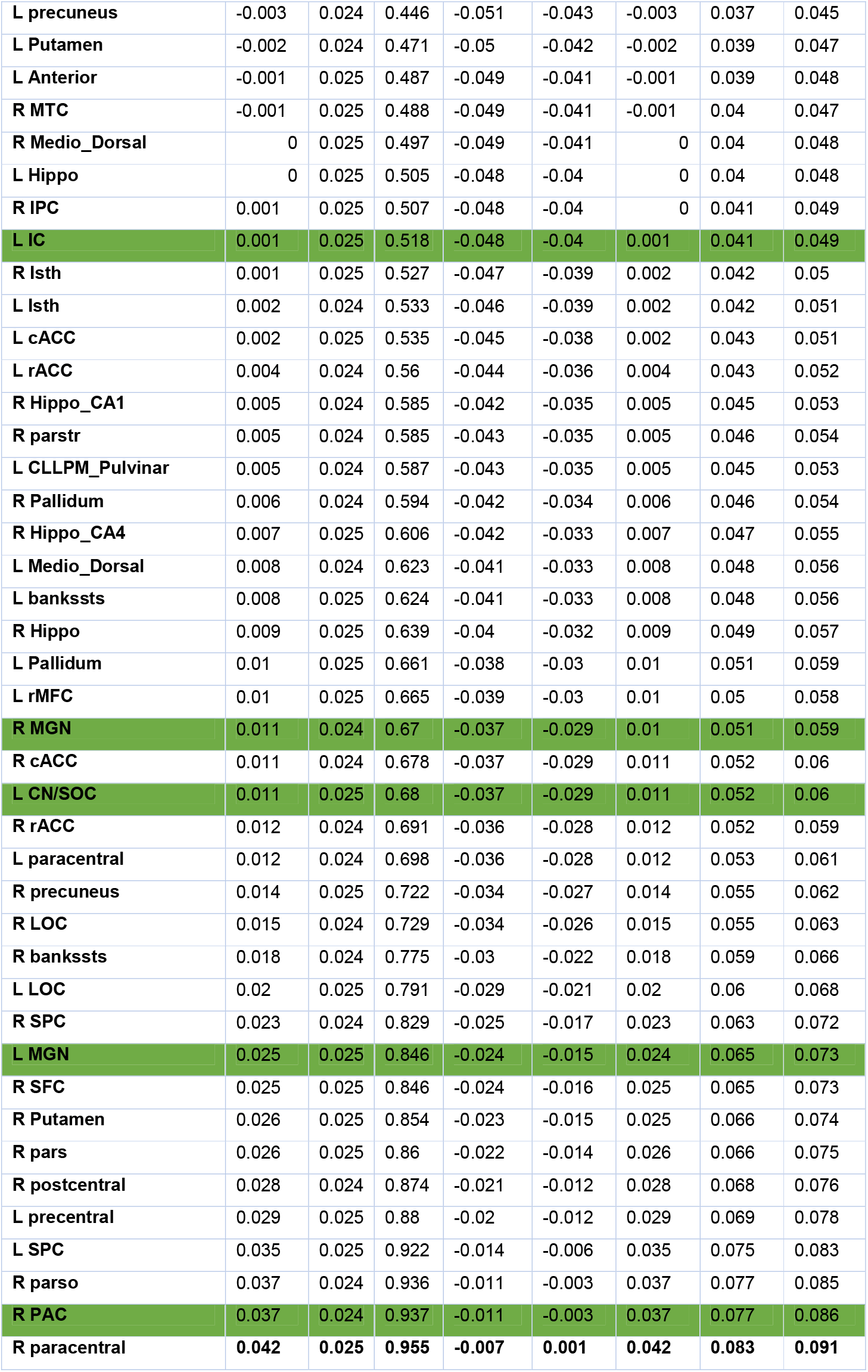
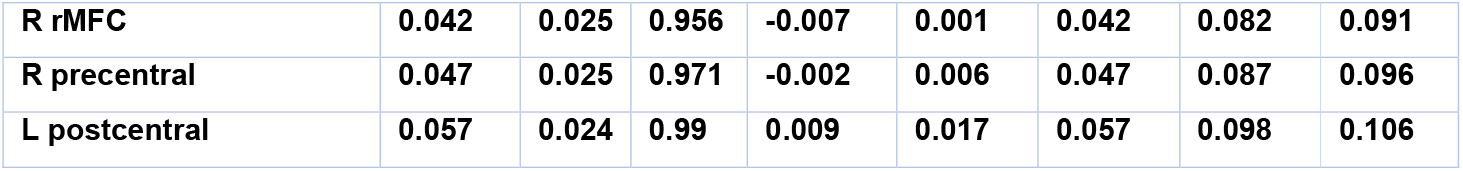
ROI effect estimates, their standard deviations, posterior probabilities (P+) for the comparison CHEU minus CHUU being positive, as well as two-sided quantile intervals. The ROIs are organized in P+ descending order. Auditory region rows are marked in green and regions showing evidence of altered FC (P+ >= 0.95 | P+<= 0.05) are in bold.

**Table 7:** Effect estimates for the ROI pairs comparison between CHUU and CHEU. P+ values are arranged in ascending order.

| ROI1 | ROI2 | mean | SD | P+ |
| --- | --- | --- | --- | --- |
| **L-postcentral** | **R-PAC** | **-0.088** | **0.047** | **0.032** |
| **L-MGN** | **L-postcentral** | **-0.083** | **0.047** | **0.04** |
| **R-rMFC** | **R-PAC** | **-0.082** | **0.047** | **0.04** |
| **L-MGN** | **R-precentral** | **-0.082** | **0.047** | **0.041** |
| **L-MGN** | **R-rMFC** | **-0.078** | **0.047** | **0.049** |
| **R-paracentral** | **R-PAC** | **-0.075** | **0.047** | **0.053** |
| **L-MGN** | R-parso | -0.075 | 0.047 | 0.054 |
| **L-SPC** | R-PAC | -0.073 | 0.047 | 0.059 |
| **L-postcentral** | L-CN-SOC | -0.072 | 0.047 | 0.064 |
| **R-precentral** | R-PAC | -0.072 | 0.047 | 0.064 |
| **L-MGN** | R-paracentral | -0.069 | 0.047 | 0.069 |
| **L-postcentral** | R-MGN | -0.068 | 0.047 | 0.073 |
| **R-precuneus** | R-PAC | -0.066 | 0.047 | 0.077 |
| **L-MGN** | L-SPC | -0.066 | 0.047 | 0.079 |
| **R-pars** | R-PAC | -0.065 | 0.047 | 0.081 |
| **L-CN-SOC** | R-precentral | -0.064 | 0.047 | 0.086 |
| **R-SFC** | R-PAC | -0.063 | 0.047 | 0.087 |
| **R-MGN** | R-rMFC | -0.063 | 0.047 | 0.087 |
| **L-MGN** | R-PAC | -0.063 | 0.047 | 0.088 |
| **L-IC** | L-postcentral | -0.062 | 0.047 | 0.091 |
| **L-MGN** | L-precentral | -0.062 | 0.047 | 0.093 |
| **L-MGN** | R-Putamen | -0.062 | 0.047 | 0.093 |
| **L-MGN** | R-SFC | -0.062 | 0.047 | 0.093 |
| **R-SPC** | R-PAC | -0.06 | 0.047 | 0.101 |
| **L-MGN** | R-pars | -0.059 | 0.047 | 0.102 |
| **L-MGN** | R-postcentral | -0.058 | 0.047 | 0.105 |
| **L-precentral** | R-PAC | -0.058 | 0.047 | 0.107 |
| **R-MGN** | R-parso | -0.058 | 0.047 | 0.107 |
| **R-parso** | R-PAC | -0.058 | 0.047 | 0.108 |
| **L-CN-SOC** | R-paracentral | -0.058 | 0.047 | 0.108 |
| **R-MGN** | R-precentral | -0.058 | 0.047 | 0.108 |
| **L-CN-SOC** | R-rMFC | -0.057 | 0.047 | 0.111 |
| **R-Isth** | R-PAC | -0.056 | 0.047 | 0.116 |
| **L-LOC** | R-PAC | -0.054 | 0.047 | 0.122 |
| **L-Isth** | R-PAC | -0.054 | 0.047 | 0.123 |
| R-LOC | R-PAC | -0.054 | 0.047 | 0.123 |
| L-IC | R-precentral | -0.054 | 0.047 | 0.123 |
| R-Hippo_CA4 | R-PAC | -0.053 | 0.047 | 0.124 |
| L-CN-SOC | R-PAC | -0.053 | 0.047 | 0.129 |
| L-CN-SOC | R-parso | -0.052 | 0.047 | 0.13 |
| R-Putamen | R-PAC | -0.052 | 0.047 | 0.131 |
| L-MGN | R-SPC | -0.052 | 0.047 | 0.132 |
| R-MGN | R-PAC | -0.051 | 0.047 | 0.137 |
| R-MGN | R-paracentral | -0.051 | 0.047 | 0.139 |
| R-Hippo | R-PAC | -0.05 | 0.047 | 0.142 |
| L-IC | R-rMFC | -0.05 | 0.047 | 0.142 |
| L-CN-SOC | L-SPC | -0.05 | 0.047 | 0.143 |
| L-Hippo | R-PAC | -0.049 | 0.047 | 0.144 |
| R-rACC | R-PAC | -0.049 | 0.047 | 0.144 |
| L-rMFC | R-PAC | -0.049 | 0.047 | 0.147 |
| L-precuneus | R-PAC | -0.049 | 0.047 | 0.148 |
| L-CN-SOC | R-SFC | -0.048 | 0.047 | 0.15 |
| L-MGN | L-rMFC | -0.048 | 0.047 | 0.151 |
| R-postcentral | R-PAC | -0.048 | 0.047 | 0.152 |
| L-postcentral | R-IC | -0.047 | 0.047 | 0.154 |
| L-paracentral | R-PAC | -0.047 | 0.047 | 0.155 |
| L-Hippo_Mol_layer_HP | R-PAC | -0.047 | 0.047 | 0.156 |
| R-Hippo_CA1 | R-PAC | -0.047 | 0.047 | 0.156 |
| L-precentral | L-CN-SOC | -0.047 | 0.047 | 0.157 |
| R-MGN | R-SFC | -0.047 | 0.047 | 0.159 |
| L-SPC | R-MGN | -0.046 | 0.047 | 0.161 |
| R-IPC | R-PAC | -0.046 | 0.047 | 0.161 |
| L-MGN | R-bankssts | -0.046 | 0.047 | 0.162 |
| L-MGN | R-LOC | -0.045 | 0.047 | 0.164 |
| L-CLLPM_Pulvinar | R-PAC | -0.045 | 0.047 | 0.164 |
| L-LOC | L-MGN | -0.045 | 0.047 | 0.165 |
| L-IC | R-paracentral | -0.045 | 0.047 | 0.166 |
| R-bankssts | R-PAC | -0.044 | 0.047 | 0.17 |
| L-MGN | R-cACC | -0.044 | 0.047 | 0.173 |
| L-Pallidum | R-PAC | -0.043 | 0.047 | 0.173 |
| L-CN-SOC | R-Putamen | -0.043 | 0.047 | 0.178 |
| L-CN-SOC | R-pars | -0.043 | 0.047 | 0.18 |
| L-Medio_Dorsal | R-PAC | -0.043 | 0.047 | 0.181 |
| R-cACC | R-PAC | -0.042 | 0.047 | 0.183 |
| R-MTC | R-PAC | -0.042 | 0.047 | 0.183 |
| L-postcentral | R-CN-SOC | -0.042 | 0.047 | 0.184 |
| L-CN-SOC | R-postcentral | -0.042 | 0.047 | 0.185 |
| **L-MGN** | R-precuneus | -0.04 | 0.047 | 0.187 |
| **R-MGN** | R-pars | -0.041 | 0.047 | 0.188 |
| **R-IC** | R-precentral | -0.041 | 0.047 | 0.19 |
| **L-precentral** | R-MGN | -0.041 | 0.047 | 0.191 |
| **L-rACC** | R-PAC | -0.041 | 0.047 | 0.191 |
| **L-MGN** | L-paracentral | -0.04 | 0.047 | 0.194 |
| **L-IC** | R-SFC | -0.04 | 0.047 | 0.194 |
| **L-IC** | L-precentral | -0.039 | 0.047 | 0.195 |
| **L-Anterior** | R-PAC | -0.04 | 0.047 | 0.195 |
| **R-MGN** | R-Putamen | -0.04 | 0.047 | 0.197 |
| **L-IC** | R-parso | -0.039 | 0.047 | 0.198 |
| **R-Hippo_GCDG** | R-PAC | -0.039 | 0.047 | 0.199 |
| **L-MGN** | R-parstr | -0.039 | 0.047 | 0.2 |
| **L-IC** | L-SPC | -0.039 | 0.047 | 0.201 |
| **L-IC** | R-PAC | -0.039 | 0.047 | 0.201 |
| **R-Anterior** | R-PAC | -0.038 | 0.047 | 0.205 |
| **L-MGN** | R-rACC | -0.038 | 0.047 | 0.205 |
| **L-Medio_Dorsal** | L-MGN | -0.038 | 0.047 | 0.207 |
| **R-parstr** | R-PAC | -0.038 | 0.047 | 0.208 |
| **L-IPC** | R-PAC | -0.038 | 0.047 | 0.209 |
| **R-Pallidum** | R-PAC | -0.038 | 0.047 | 0.21 |
| **L-MGN** | L-Pallidum | -0.037 | 0.047 | 0.212 |
| **L-bankssts** | L-MGN | -0.037 | 0.047 | 0.213 |
| **L-CN-SOC** | R-bankssts | -0.036 | 0.047 | 0.218 |
| **L-CN-SOC** | R-precuneus | -0.036 | 0.047 | 0.219 |
| **L-Hippo_Tail** | R-PAC | -0.036 | 0.047 | 0.221 |
| **L-PAC** | R-rMFC | -0.035 | 0.047 | 0.223 |
| **L-MGN** | R-Medio_Dorsal | -0.035 | 0.047 | 0.225 |
| **L-cMFC** | R-PAC | -0.035 | 0.047 | 0.225 |
| **L-SFC** | R-PAC | -0.035 | 0.047 | 0.225 |
| **R-MGN** | R-postcentral | -0.035 | 0.047 | 0.225 |
| **L-rMFC** | R-MGN | -0.035 | 0.047 | 0.226 |
| **L-CN-SOC** | R-SPC | -0.035 | 0.047 | 0.227 |
| **R-precentral** | R-CN-SOC | -0.035 | 0.047 | 0.228 |
| **R-MGN** | R-precuneus | -0.034 | 0.047 | 0.229 |
| **L-MGN** | L-parso | -0.034 | 0.047 | 0.23 |
| **L-PAC** | L-postcentral | -0.034 | 0.047 | 0.231 |
| **R-Medio_Dorsal** | R-PAC | -0.034 | 0.047 | 0.232 |
| **L-bankssts** | R-PAC | -0.033 | 0.047 | 0.236 |
| **L-cACC** | L-MGN | -0.033 | 0.047 | 0.237 |
| **L-MGN** | R-Pallidum | -0.033 | 0.047 | 0.237 |
| **L-IC** | R-pars | -0.033 | 0.047 | 0.238 |
| **L-MGN** | L-Putamen | -0.033 | 0.047 | 0.24 |
| **R-IC** | R-rMFC | -0.033 | 0.047 | 0.241 |
| **L-IC** | R-postcentral | -0.033 | 0.047 | 0.242 |
| **R-MGN** | R-SPC | -0.032 | 0.047 | 0.244 |
| **L-paracentral** | L-CN-SOC | -0.032 | 0.047 | 0.245 |
| **L-MGN** | R-MTC | -0.031 | 0.047 | 0.25 |
| **L-Anterior** | L-MGN | -0.031 | 0.047 | 0.253 |
| **L-cACC** | R-PAC | -0.03 | 0.047 | 0.257 |
| **R-bankssts** | R-MGN | -0.03 | 0.047 | 0.258 |
| **R-IC** | R-paracentral | -0.03 | 0.047 | 0.258 |
| **L-rMFC** | L-CN-SOC | -0.03 | 0.047 | 0.259 |
| **L-LOC** | L-CN-SOC | -0.03 | 0.047 | 0.26 |
| **L-Putamen** | R-PAC | -0.03 | 0.047 | 0.26 |
| **L-MGN** | R-IPC | -0.029 | 0.047 | 0.261 |
| **R-cACC** | R-MGN | -0.03 | 0.047 | 0.261 |
| **L-Hippo_Pres** | R-PAC | -0.03 | 0.047 | 0.263 |
| **L-IC** | R-Putamen | -0.029 | 0.047 | 0.263 |
| **R-Hippo_CA3** | R-PAC | -0.029 | 0.047 | 0.264 |
| **L-MGN** | L-rACC | -0.029 | 0.047 | 0.266 |
| **R-Hippo_Tail** | R-PAC | -0.029 | 0.047 | 0.266 |
| **L-CN-SOC** | R-rACC | -0.028 | 0.047 | 0.266 |
| **R-paracentral** | R-CN-SOC | -0.029 | 0.047 | 0.268 |
| **L-MGN** | L-SFC | -0.028 | 0.047 | 0.27 |
| **L-Pallidum** | R-MGN | -0.028 | 0.047 | 0.273 |
| **L-MGN** | R-Anterior | -0.028 | 0.047 | 0.275 |
| **L-CN-SOC** | R-parstr | -0.027 | 0.047 | 0.275 |
| **L-IC** | R-precuneus | -0.027 | 0.047 | 0.276 |
| **L-MGN** | R-Hippo | -0.027 | 0.047 | 0.278 |
| **L-MGN** | R-Isth | -0.027 | 0.047 | 0.279 |
| **L-MGN** | L-CN-SOC | -0.026 | 0.047 | 0.282 |
| **R-cuneus** | R-PAC | -0.026 | 0.047 | 0.282 |
| **R-rMFC** | R-CN-SOC | -0.027 | 0.047 | 0.283 |
| **L-CN-SOC** | R-cACC | -0.026 | 0.047 | 0.289 |
| **L-bankssts** | R-MGN | -0.025 | 0.047 | 0.291 |
| **L-Isth** | L-MGN | -0.025 | 0.047 | 0.292 |
| **L-CN-SOC** | R-LOC | -0.025 | 0.047 | 0.294 |
| **L-PAC** | L-SPC | -0.025 | 0.047 | 0.295 |
| **L-MGN** | R-Hippo_CA4 | -0.025 | 0.047 | 0.295 |
| **L-cMFC** | L-MGN | -0.025 | 0.047 | 0.296 |
| **L-Hippo_CA4** | R-PAC | -0.024 | 0.047 | 0.297 |
| **L-parstr** | R-PAC | -0.025 | 0.047 | 0.297 |
| **R-MGN** | R-rACC | -0.024 | 0.047 | 0.297 |
| **R-IC** | R-SFC | -0.025 | 0.047 | 0.297 |
| **L-PAC** | R-precentral | -0.024 | 0.047 | 0.299 |
| **L-MGN** | R-V_Anterior | -0.024 | 0.047 | 0.3 |
| **L-Medio_Dorsal** | L-CN-SOC | -0.024 | 0.047 | 0.303 |
| **L-IC** | L-LOC | -0.024 | 0.047 | 0.303 |
| **L-CN-SOC** | R-IPC | -0.024 | 0.047 | 0.304 |
| **L-precentral** | R-IC | -0.024 | 0.047 | 0.305 |
| **L-LOC** | R-MGN | -0.024 | 0.047 | 0.305 |
| **R-MGN** | R-parstr | -0.024 | 0.047 | 0.306 |
| **L-SPC** | R-IC | -0.023 | 0.047 | 0.308 |
| **L-CN-SOC** | R-Isth | -0.023 | 0.047 | 0.308 |
| **R-IC** | R-parso | -0.023 | 0.047 | 0.308 |
| **L-IC** | R-SPC | -0.023 | 0.047 | 0.308 |
| **L-Pallidum** | L-CN-SOC | -0.023 | 0.047 | 0.309 |
| **L-MGN** | L-precuneus | -0.023 | 0.047 | 0.309 |
| **L-VLD** | R-PAC | -0.023 | 0.047 | 0.311 |
| **L-cuneus** | R-PAC | -0.022 | 0.047 | 0.315 |
| **R-MGN** | R-Pallidum | -0.022 | 0.047 | 0.315 |
| **R-V_Anterior** | R-PAC | -0.022 | 0.047 | 0.316 |
| **L-paracentral** | R-MGN | -0.022 | 0.047 | 0.318 |
| **R-PCC** | R-PAC | -0.021 | 0.047 | 0.318 |
| **R-CLLPM_Pulvinar** | R-PAC | -0.021 | 0.047 | 0.321 |
| **L-IC** | L-rMFC | -0.021 | 0.047 | 0.322 |
| **L-Medio_Dorsal** | R-MGN | -0.021 | 0.047 | 0.322 |
| **L-Pulvinar** | R-PAC | -0.021 | 0.047 | 0.322 |
| **L-Isth** | L-CN-SOC | -0.021 | 0.047 | 0.323 |
| **L-MGN** | L-PAC | -0.021 | 0.047 | 0.323 |
| **L-MGN** | R-Hippo_CA1 | -0.021 | 0.047 | 0.325 |
| **L-bankssts** | L-CN-SOC | -0.021 | 0.047 | 0.326 |
| **L-MGN** | L-parstr | -0.021 | 0.047 | 0.326 |
| **L-IC** | R-bankssts | -0.021 | 0.047 | 0.326 |
| **L-rACC** | L-CN-SOC | -0.02 | 0.047 | 0.327 |
| **R-IC** | R-PAC | -0.021 | 0.047 | 0.327 |
| **R-parso** | R-CN-SOC | -0.021 | 0.047 | 0.329 |
| **R-IC** | R-postcentral | -0.02 | 0.047 | 0.329 |
| **L-IC** | R-LOC | -0.02 | 0.047 | 0.33 |
| **L-PAC** | R-pars | -0.02 | 0.047 | 0.332 |
| **L-Putamen** | L-CN-SOC | -0.02 | 0.047 | 0.334 |
| **L-precuneus** | L-CN-SOC | -0.02 | 0.047 | 0.335 |
| **L-CN-SOC** | R-MTC | -0.019 | 0.047 | 0.335 |
| **L-IC** | L-paracentral | -0.02 | 0.047 | 0.337 |
| **L-MGN** | R-PCC | -0.019 | 0.047 | 0.338 |
| **L-SFC** | L-CN-SOC | -0.019 | 0.047 | 0.339 |
| **R-LOC** | R-MGN | -0.019 | 0.047 | 0.339 |
| **L-PAC** | R-precuneus | -0.019 | 0.047 | 0.339 |
| **L-IC** | L-MGN | -0.019 | 0.047 | 0.34 |
| **L-cACC** | R-MGN | -0.019 | 0.047 | 0.341 |
| **L-parso** | R-PAC | -0.018 | 0.047 | 0.345 |
| **R-Anterior** | R-MGN | -0.018 | 0.047 | 0.348 |
| **L-PAC** | R-paracentral | -0.018 | 0.047 | 0.348 |
| **L-IC** | R-cACC | -0.018 | 0.047 | 0.35 |
| **L-CN-SOC** | R-Medio_Dorsal | -0.018 | 0.047 | 0.352 |
| **L-PCC** | R-PAC | -0.017 | 0.047 | 0.352 |
| **L-parso** | L-CN-SOC | -0.017 | 0.047 | 0.354 |
| **L-precentral** | R-CN-SOC | -0.017 | 0.047 | 0.355 |
| **L-cACC** | L-CN-SOC | -0.017 | 0.047 | 0.356 |
| **R-CN-SOC** | R-PAC | -0.017 | 0.047 | 0.357 |
| **L-parso** | R-MGN | -0.017 | 0.047 | 0.358 |
| **L-PAC** | R-parso | -0.017 | 0.047 | 0.359 |
| **R-IPC** | R-MGN | -0.016 | 0.047 | 0.361 |
| **R-SFC** | R-CN-SOC | -0.016 | 0.047 | 0.362 |
| **L-CN-SOC** | R-Hippo | -0.017 | 0.047 | 0.362 |
| **L-rACC** | R-MGN | -0.016 | 0.047 | 0.365 |
| **L-IPC** | L-MGN | -0.016 | 0.047 | 0.366 |
| **L-Anterior** | R-MGN | -0.016 | 0.047 | 0.366 |
| **R-IC** | R-pars | -0.016 | 0.047 | 0.366 |
| **L-IC** | R-rACC | -0.015 | 0.047 | 0.367 |
| **L-PAC** | R-SFC | -0.016 | 0.047 | 0.367 |
| **L-precuneus** | R-MGN | -0.016 | 0.047 | 0.368 |
| **L-IC** | L-precuneus | -0.015 | 0.047 | 0.369 |
| **L-SPC** | R-CN-SOC | -0.016 | 0.047 | 0.369 |
| **L-MGN** | R-MGN | -0.015 | 0.047 | 0.371 |
| **L-CN-SOC** | R-MGN | -0.015 | 0.047 | 0.371 |
| **L-PAC** | R-SPC | -0.015 | 0.047 | 0.372 |
| **L-CLLPM_Pulvinar** | L-MGN | -0.015 | 0.047 | 0.373 |
| **L-Hippo_CA1** | R-PAC | -0.015 | 0.047 | 0.373 |
| **L-parstr** | L-CN-SOC | -0.015 | 0.047 | 0.374 |
| **R-Isth** | R-MGN | -0.015 | 0.047 | 0.376 |
| **L-IC** | L-Isth | -0.015 | 0.047 | 0.378 |
| **L-CN-SOC** | R-Pallidum | -0.014 | 0.047 | 0.378 |
| **R-Medio_Dorsal** | R-MGN | -0.014 | 0.047 | 0.379 |
| **R-postcentral** | R-CN-SOC | -0.014 | 0.047 | 0.38 |
| **L-MGN** | L-VLD | -0.014 | 0.047 | 0.382 |
| **L-CN-SOC** | R-Hippo_CA4 | -0.014 | 0.047 | 0.382 |
| **L-IC** | R-Isth | -0.014 | 0.047 | 0.382 |
| **L-PAC** | R-LOC | -0.014 | 0.047 | 0.382 |
| **L-Isth** | R-MGN | -0.014 | 0.047 | 0.382 |
| **L-CLLPM_Pulvinar** | L-CN-SOC | -0.014 | 0.047 | 0.383 |
| **R-MGN** | R-MTC | -0.014 | 0.047 | 0.383 |
| **L-MGN** | R-Caudate | -0.014 | 0.047 | 0.384 |
| **L-SFC** | R-MGN | -0.014 | 0.047 | 0.384 |
| **L-Hippo** | L-CN-SOC | -0.013 | 0.047 | 0.385 |
| **L-Hippo_Mol_layer_HP** | L-CN-SOC | -0.013 | 0.047 | 0.386 |
| **L-LOC** | L-PAC | -0.013 | 0.047 | 0.387 |
| **L-CN-SOC** | R-Hippo_CA1 | -0.013 | 0.047 | 0.387 |
| **R-Hippo_CA1** | R-MGN | -0.013 | 0.047 | 0.387 |
| **L-PAC** | R-Hippo | -0.013 | 0.047 | 0.389 |
| **L-IC** | R-IPC | -0.013 | 0.047 | 0.39 |
| **R-Caudate** | R-PAC | -0.013 | 0.047 | 0.392 |
| **R-VLD** | R-PAC | -0.012 | 0.047 | 0.392 |
| **R-pars** | R-CN-SOC | -0.012 | 0.047 | 0.395 |
| **L-Putamen** | R-MGN | -0.012 | 0.047 | 0.4 |
| **L-cMFC** | R-MGN | -0.012 | 0.047 | 0.401 |
| **R-Hippo_CA4** | R-MGN | -0.011 | 0.047 | 0.401 |
| **L-Isth** | L-PAC | -0.011 | 0.047 | 0.403 |
| **L-IC** | L-rACC | -0.011 | 0.047 | 0.407 |
| **R-pericalc** | R-PAC | -0.011 | 0.047 | 0.407 |
| **R-IC** | R-Putamen | -0.011 | 0.047 | 0.408 |
| **L-PAC** | L-CN-SOC | -0.01 | 0.047 | 0.41 |
| **L-MGN** | R-Hippo_Tail | -0.011 | 0.047 | 0.41 |
| **L-IC** | L-SFC | -0.011 | 0.047 | 0.412 |
| **L-IC** | L-Pallidum | -0.01 | 0.047 | 0.417 |
| **L-PAC** | R-Hippo_CA4 | -0.01 | 0.047 | 0.418 |
| **L-parstr** | R-MGN | -0.01 | 0.047 | 0.419 |
| **L-Hippo** | L-MGN | -0.009 | 0.047 | 0.42 |
| **R-IC** | R-SPC | -0.009 | 0.047 | 0.423 |
| **L-IC** | R-parstr | -0.009 | 0.047 | 0.424 |
| **L-MGN** | R-VLD | -0.009 | 0.047 | 0.425 |
| **L-MGN** | R-cuneus | -0.009 | 0.047 | 0.426 |
| **L-Hippo_Tail** | L-MGN | -0.009 | 0.047 | 0.427 |
| **L-MGN** | R-Hippo_GCDG | -0.009 | 0.047 | 0.427 |
| **L-IC** | R-MTC | -0.008 | 0.047 | 0.428 |
| **L-IPC** | L-CN-SOC | -0.008 | 0.047 | 0.429 |
| **L-cACC** | L-IC | -0.008 | 0.047 | 0.43 |
| **L-IC** | L-Medio_Dorsal | -0.008 | 0.047 | 0.43 |
| **L-cMFC** | L-CN-SOC | -0.008 | 0.047 | 0.431 |
| **L-Hippo** | L-PAC | -0.008 | 0.047 | 0.431 |
| **L-IC** | R-MGN | -0.008 | 0.047 | 0.431 |
| **L-bankssts** | L-IC | -0.008 | 0.047 | 0.432 |
| **L-CLLPM_Pulvinar** | L-IC | -0.008 | 0.047 | 0.432 |
| **L-PAC** | R-Isth | -0.008 | 0.047 | 0.433 |
| **L-Hippo_Mol_layer_HP** | L-MGN | -0.008 | 0.047 | 0.434 |
| **L-MGN** | L-V_Anterior | -0.008 | 0.047 | 0.434 |
| **L-PAC** | R-Putamen | -0.007 | 0.047 | 0.436 |
| **L-Hippo_CA3** | R-PAC | -0.007 | 0.047 | 0.437 |
| **L-CN-SOC** | L-VLD | -0.007 | 0.047 | 0.438 |
| **R-Hippo** | R-MGN | -0.007 | 0.047 | 0.439 |
| **R-IC** | R-precuneus | -0.007 | 0.047 | 0.439 |
| **L-V_Anterior** | R-PAC | -0.007 | 0.047 | 0.44 |
| **L-MGN** | L-PCC | -0.007 | 0.047 | 0.441 |
| **L-LOC** | R-IC | -0.007 | 0.047 | 0.442 |
| **L-PAC** | R-IPC | -0.007 | 0.047 | 0.442 |
| **R-MGN** | R-V_Anterior | -0.007 | 0.047 | 0.442 |
| **R-CN-SOC** | R-SPC | -0.006 | 0.047 | 0.443 |
| **L-PAC** | L-precuneus | -0.006 | 0.047 | 0.444 |
| **R-Putamen** | R-CN-SOC | -0.006 | 0.047 | 0.445 |
| **L-Hippo_Tail** | L-CN-SOC | -0.006 | 0.047 | 0.447 |
| **L-PAC** | R-rACC | -0.006 | 0.047 | 0.447 |
| **R-bankssts** | R-CN-SOC | -0.006 | 0.047 | 0.448 |
| **L-IC** | R-Hippo | -0.006 | 0.047 | 0.45 |
| **L-PAC** | L-rACC | -0.006 | 0.047 | 0.451 |
| **L-CLLPM_Pulvinar** | R-MGN | -0.005 | 0.047 | 0.452 |
| **L-MGN** | R-CLLPM_Pulvinar | -0.005 | 0.047 | 0.453 |
| **L-PAC** | R-Hippo_CA1 | -0.005 | 0.047 | 0.454 |
| **L-cuneus** | L-MGN | -0.005 | 0.047 | 0.455 |
| **L-CN-SOC** | R-PCC | -0.005 | 0.047 | 0.455 |
| **L-Hippo** | R-MGN | -0.005 | 0.047 | 0.458 |
| **R-cMFC** | R-PAC | -0.005 | 0.047 | 0.458 |
| **L-IC** | L-Putamen | -0.005 | 0.047 | 0.459 |
| **L-IPC** | R-MGN | -0.005 | 0.047 | 0.46 |
| **L-CN-SOC** | R-Hippo_Tail | -0.004 | 0.047 | 0.461 |
| **L-PAC** | R-MGN | -0.004 | 0.047 | 0.461 |
| **L-Anterior** | L-CN-SOC | -0.004 | 0.047 | 0.464 |
| **R-bankssts** | R-IC | -0.004 | 0.047 | 0.464 |
| **L-CN-SOC** | R-Anterior | -0.004 | 0.047 | 0.466 |
| **R-lingual** | R-PAC | -0.004 | 0.047 | 0.466 |
| **R-IC** | R-LOC | -0.004 | 0.047 | 0.467 |
| **L-IC** | R-Pallidum | -0.004 | 0.047 | 0.468 |
| **L-cMFC** | L-IC | -0.003 | 0.047 | 0.474 |
| **L-IC** | R-Hippo_CA1 | -0.003 | 0.047 | 0.474 |
| **L-CN-SOC** | R-V_Anterior | -0.003 | 0.047 | 0.475 |
| **L-rMFC** | R-IC | -0.003 | 0.047 | 0.476 |
| **L-PAC** | L-rMFC | -0.002 | 0.047 | 0.477 |
| **L-IC** | R-Hippo_CA4 | -0.003 | 0.047 | 0.477 |
| **L-Hippo_Mol_layer_HP** | L-PAC | -0.003 | 0.047 | 0.478 |
| **L-PAC** | R-postcentral | -0.003 | 0.047 | 0.478 |
| **L-paracentral** | R-IC | -0.002 | 0.047 | 0.479 |
| **L-IC** | L-parso | -0.002 | 0.047 | 0.483 |
| **R-cACC** | R-IC | -0.002 | 0.047 | 0.483 |
| **L-IC** | L-CN-SOC | -0.002 | 0.047 | 0.486 |
| **L-Hippo** | L-IC | -0.001 | 0.047 | 0.489 |
| **L-PAC** | L-precentral | -0.001 | 0.047 | 0.492 |
| **L-MGN** | R-Hippo_CA3 | -0.001 | 0.047 | 0.494 |
| **L-MGN** | R-IC | 0 | 0.047 | 0.496 |
| **L-VLD** | R-MGN | 0 | 0.047 | 0.497 |
| **L-IC** | L-IPC | 0 | 0.047 | 0.498 |
| **L-Hippo_Mol_layer_HP** | R-MGN | 0 | 0.047 | 0.498 |
| **L-paracentral** | R-CN-SOC | 0 | 0.047 | 0.499 |
| **L-CN-SOC** | R-Hippo_GCDG | 0 | 0.047 | 0.5 |
| **R-IC** | R-rACC | 0 | 0.047 | 0.5 |
| **L-Hippo_Mol_layer_HP** | L-IC | 0 | 0.047 | 0.502 |
| **L-CLLPM_Pulvinar** | L-PAC | 0 | 0.047 | 0.502 |
| **L-lingual** | R-PAC | 0.001 | 0.047 | 0.503 |
| **R-MGN** | R-PCC | 0 | 0.047 | 0.504 |
| **R-precuneus** | R-CN-SOC | 0.001 | 0.047 | 0.506 |
| **L-PCC** | L-CN-SOC | 0.001 | 0.047 | 0.507 |
| **L-CN-SOC** | R-cuneus | 0.001 | 0.047 | 0.507 |
| **R-Caudate** | R-MGN | 0.001 | 0.047 | 0.511 |
| **L-IC** | L-parstr | 0.002 | 0.047 | 0.516 |
| **L-Anterior** | L-PAC | 0.002 | 0.047 | 0.518 |
| **L-cuneus** | L-CN-SOC | 0.002 | 0.047 | 0.519 |
| **L-MGN** | R-cMFC | 0.002 | 0.047 | 0.521 |
| **L-CN-SOC** | R-CLLPM_Pulvinar | 0.003 | 0.047 | 0.522 |
| **L-LOC** | R-CN-SOC | 0.003 | 0.047 | 0.524 |
| **L-PAC** | L-Pallidum | 0.003 | 0.047 | 0.53 |
| **L-PAC** | R-PAC | 0.004 | 0.047 | 0.533 |
| **R-cuneus** | R-MGN | 0.004 | 0.047 | 0.534 |
| **R-IC** | R-Isth | 0.004 | 0.047 | 0.537 |
| **L-Isth** | R-IC | 0.004 | 0.047 | 0.538 |
| **L-IC** | L-VLD | 0.005 | 0.047 | 0.539 |
| L-IC | R-Hippo_Tail | 0.004 | 0.047 | 0.539 |
| L-PAC | R-Hippo_GCDG | 0.005 | 0.047 | 0.541 |
| L-Caudate | R-PAC | 0.005 | 0.047 | 0.542 |
| R-rACC | R-CN-SOC | 0.005 | 0.047 | 0.543 |
| L-Hippo_Tail | R-MGN | 0.005 | 0.047 | 0.543 |
| L-Hippo_Tail | L-IC | 0.005 | 0.047 | 0.544 |
| R-parstr | R-CN-SOC | 0.005 | 0.047 | 0.544 |
| L-Hippo_Pres | L-CN-SOC | 0.005 | 0.047 | 0.545 |
| L-rMFC | R-CN-SOC | 0.006 | 0.047 | 0.545 |
| L-cuneus | R-MGN | 0.005 | 0.047 | 0.547 |
| L-bankssts | R-IC | 0.006 | 0.047 | 0.548 |
| L-rACC | R-IC | 0.006 | 0.047 | 0.549 |
| L-IC | R-Medio_Dorsal | 0.006 | 0.047 | 0.55 |
| L-IPC | L-PAC | 0.006 | 0.047 | 0.551 |
| L-MGN | L-Pulvinar | 0.006 | 0.047 | 0.551 |
| R-IC | R-IPC | 0.006 | 0.047 | 0.552 |
| L-Caudate | L-MGN | 0.006 | 0.047 | 0.553 |
| R-cACC | R-CN-SOC | 0.006 | 0.047 | 0.553 |
| L-PAC | R-Pallidum | 0.006 | 0.047 | 0.554 |
| L-Anterior | L-IC | 0.006 | 0.047 | 0.555 |
| L-SFC | R-IC | 0.007 | 0.047 | 0.556 |
| L-pericalc | R-PAC | 0.006 | 0.047 | 0.556 |
| L-CN-SOC | R-VLD | 0.007 | 0.047 | 0.558 |
| L-PAC | L-paracentral | 0.007 | 0.047 | 0.56 |
| L-IC | R-PCC | 0.007 | 0.047 | 0.561 |
| L-PAC | R-bankssts | 0.007 | 0.047 | 0.562 |
| R-MTC | R-CN-SOC | 0.007 | 0.047 | 0.563 |
| L-Hippo_CA4 | L-CN-SOC | 0.008 | 0.047 | 0.564 |
| L-Hippo_CA4 | L-MGN | 0.007 | 0.047 | 0.564 |
| L-precuneus | R-IC | 0.007 | 0.047 | 0.564 |
| R-Hippo_GCDG | R-MGN | 0.007 | 0.047 | 0.565 |
| L-IC | L-PAC | 0.008 | 0.047 | 0.566 |
| L-cACC | R-IC | 0.008 | 0.047 | 0.566 |
| L-Pallidum | R-IC | 0.008 | 0.047 | 0.566 |
| R-Hippo_Tail | R-MGN | 0.008 | 0.047 | 0.566 |
| L-MGN | R-CN-SOC | 0.008 | 0.047 | 0.567 |
| L-CN-SOC | R-IC | 0.008 | 0.047 | 0.568 |
| L-MGN | R-pericalc | 0.008 | 0.047 | 0.569 |
| L-V_Anterior | R-MGN | 0.008 | 0.047 | 0.571 |
| L-CLLPM_Pulvinar | R-IC | 0.008 | 0.047 | 0.572 |
| R-IC | R-parstr | 0.008 | 0.047 | 0.572 |
| L-IC | R-Hippo_GCDG | 0.009 | 0.047 | 0.573 |
| **L-cMFC** | L-PAC | 0.009 | 0.047 | 0.575 |
| **L-PAC** | R-Anterior | 0.009 | 0.047 | 0.575 |
| **L-cuneus** | L-IC | 0.009 | 0.047 | 0.576 |
| **L-PAC** | R-cACC | 0.009 | 0.047 | 0.576 |
| **L-Medio_Dorsal** | R-IC | 0.009 | 0.047 | 0.577 |
| **L-PCC** | R-MGN | 0.009 | 0.047 | 0.579 |
| **R-MGN** | R-VLD | 0.009 | 0.047 | 0.579 |
| **L-IC** | L-PCC | 0.009 | 0.047 | 0.581 |
| **L-IC** | R-cuneus | 0.01 | 0.047 | 0.582 |
| **L-Hippo_CA1** | L-MGN | 0.01 | 0.047 | 0.584 |
| **L-PAC** | R-MTC | 0.01 | 0.047 | 0.584 |
| **L-IC** | R-Anterior | 0.01 | 0.047 | 0.585 |
| **R-LOC** | R-CN-SOC | 0.01 | 0.047 | 0.586 |
| **L-MGN** | R-lingual | 0.01 | 0.047 | 0.587 |
| **L-bankssts** | R-CN-SOC | 0.011 | 0.047 | 0.589 |
| **L-Medio_Dorsal** | L-PAC | 0.011 | 0.047 | 0.59 |
| **L-PAC** | L-SFC | 0.011 | 0.047 | 0.593 |
| **L-CN-SOC** | R-Caudate | 0.011 | 0.047 | 0.593 |
| **R-IC** | R-MTC | 0.011 | 0.047 | 0.595 |
| **L-IC** | R-V_Anterior | 0.011 | 0.047 | 0.595 |
| **L-cMFC** | R-IC | 0.012 | 0.047 | 0.597 |
| **R-Isth** | R-CN-SOC | 0.012 | 0.047 | 0.599 |
| **L-Hippo_Pres** | L-PAC | 0.012 | 0.047 | 0.6 |
| **L-rACC** | R-CN-SOC | 0.012 | 0.047 | 0.601 |
| **R-IC** | R-Pallidum | 0.012 | 0.047 | 0.602 |
| **L-Hippo_CA1** | L-CN-SOC | 0.012 | 0.047 | 0.604 |
| **L-Medio_Dorsal** | R-CN-SOC | 0.013 | 0.047 | 0.607 |
| **R-IPC** | R-CN-SOC | 0.013 | 0.047 | 0.608 |
| **L-Isth** | R-CN-SOC | 0.013 | 0.047 | 0.61 |
| **R-cMFC** | R-MGN | 0.013 | 0.047 | 0.61 |
| **L-Pallidum** | R-CN-SOC | 0.013 | 0.047 | 0.611 |
| **L-CN-SOC** | R-Hippo_CA3 | 0.013 | 0.047 | 0.612 |
| **L-CN-SOC** | L-V_Anterior | 0.013 | 0.047 | 0.613 |
| **L-PAC** | R-parstr | 0.013 | 0.047 | 0.613 |
| **L-Hippo_Tail** | L-PAC | 0.014 | 0.047 | 0.618 |
| **L-PAC** | R-Hippo_CA3 | 0.014 | 0.047 | 0.619 |
| **R-CLLPM_Pulvinar** | R-MGN | 0.015 | 0.047 | 0.623 |
| **R-Hippo_CA1** | R-IC | 0.015 | 0.047 | 0.624 |
| **L-cACC** | R-CN-SOC | 0.015 | 0.047 | 0.625 |
| **L-Hippo_CA4** | R-MGN | 0.015 | 0.047 | 0.626 |
| **L-bankssts** | L-PAC | 0.015 | 0.047 | 0.627 |
| **L-PAC** | R-Hippo_Tail | 0.015 | 0.047 | 0.631 |
| R-Hippo_CA4 | R-IC | 0.016 | 0.047 | 0.633 |
| L-IC | R-CLLPM_Pulvinar | 0.016 | 0.047 | 0.636 |
| R-Hippo | R-IC | 0.016 | 0.047 | 0.636 |
| R-Hippo_CA1 | R-CN-SOC | 0.017 | 0.047 | 0.64 |
| R-MGN | R-CN-SOC | 0.017 | 0.047 | 0.64 |
| L-Putamen | R-IC | 0.017 | 0.047 | 0.64 |
| L-CN-SOC | R-cMFC | 0.017 | 0.047 | 0.641 |
| L-Hippo_CA1 | R-MGN | 0.017 | 0.047 | 0.644 |
| R-MGN | R-pericalc | 0.017 | 0.047 | 0.644 |
| L-Pulvinar | R-MGN | 0.017 | 0.047 | 0.646 |
| L-SFC | R-CN-SOC | 0.018 | 0.047 | 0.647 |
| R-Hippo_CA4 | R-CN-SOC | 0.018 | 0.047 | 0.647 |
| L-IC | R-VLD | 0.018 | 0.047 | 0.647 |
| R-IC | R-MGN | 0.018 | 0.047 | 0.648 |
| L-cuneus | L-PAC | 0.018 | 0.047 | 0.649 |
| L-Hippo | R-IC | 0.018 | 0.047 | 0.65 |
| L-PAC | L-VLD | 0.018 | 0.047 | 0.652 |
| L-CLLPM_Pulvinar | R-CN-SOC | 0.018 | 0.047 | 0.653 |
| L-parso | R-IC | 0.018 | 0.047 | 0.653 |
| R-Hippo | R-CN-SOC | 0.018 | 0.047 | 0.654 |
| R-Hippo_CA3 | R-MGN | 0.018 | 0.047 | 0.654 |
| L-cACC | L-PAC | 0.019 | 0.047 | 0.657 |
| L-CN-SOC | R-lingual | 0.019 | 0.047 | 0.658 |
| L-Putamen | R-CN-SOC | 0.019 | 0.047 | 0.659 |
| R-IC | R-Medio_Dorsal | 0.019 | 0.047 | 0.659 |
| L-Hippo_CA4 | L-PAC | 0.019 | 0.047 | 0.661 |
| L-IC | R-Caudate | 0.019 | 0.047 | 0.661 |
| L-parso | R-CN-SOC | 0.019 | 0.047 | 0.662 |
| L-Caudate | R-MGN | 0.019 | 0.047 | 0.662 |
| L-precuneus | R-CN-SOC | 0.02 | 0.047 | 0.664 |
| L-Hippo_Pres | R-MGN | 0.019 | 0.047 | 0.664 |
| L-IPC | R-IC | 0.02 | 0.047 | 0.665 |
| L-PAC | R-cuneus | 0.02 | 0.047 | 0.668 |
| L-VLD | R-IC | 0.02 | 0.047 | 0.668 |
| L-Pulvinar | L-CN-SOC | 0.02 | 0.047 | 0.669 |
| L-Hippo_Mol_layer_HP | R-IC | 0.021 | 0.047 | 0.67 |
| L-IC | R-Hippo_CA3 | 0.021 | 0.047 | 0.672 |
| L-PAC | R-Medio_Dorsal | 0.021 | 0.047 | 0.672 |
| L-CN-SOC | R-pericalc | 0.02 | 0.047 | 0.673 |
| L-Hippo_CA4 | L-IC | 0.021 | 0.047 | 0.676 |
| R-Medio_Dorsal | R-CN-SOC | 0.022 | 0.047 | 0.677 |
| R-Pallidum | R-CN-SOC | 0.021 | 0.047 | 0.679 |
| L-Hippo_Pres | L-IC | 0.022 | 0.047 | 0.68 |
| L-parstr | R-CN-SOC | 0.022 | 0.047 | 0.681 |
| L-PAC | R-CN-SOC | 0.022 | 0.047 | 0.682 |
| L-parstr | R-IC | 0.022 | 0.047 | 0.682 |
| L-PAC | L-Pulvinar | 0.022 | 0.047 | 0.683 |
| L-lingual | L-MGN | 0.022 | 0.047 | 0.685 |
| L-PAC | L-parstr | 0.023 | 0.047 | 0.688 |
| L-Hippo_Pres | L-MGN | 0.023 | 0.047 | 0.691 |
| L-IC | R-cMFC | 0.023 | 0.047 | 0.691 |
| L-Hippo | R-CN-SOC | 0.023 | 0.047 | 0.692 |
| L-PAC | R-IC | 0.023 | 0.047 | 0.693 |
| R-IC | R-PCC | 0.024 | 0.047 | 0.701 |
| L-Hippo_Tail | R-IC | 0.025 | 0.047 | 0.704 |
| R-IC | R-V_Anterior | 0.025 | 0.047 | 0.708 |
| L-PAC | L-Putamen | 0.026 | 0.047 | 0.712 |
| L-IC | L-V_Anterior | 0.026 | 0.047 | 0.714 |
| L-VLD | R-CN-SOC | 0.026 | 0.047 | 0.715 |
| L-Hippo_Mol_layer_HP | R-CN-SOC | 0.027 | 0.047 | 0.719 |
| R-Anterior | R-IC | 0.027 | 0.047 | 0.719 |
| L-PAC | R-V_Anterior | 0.027 | 0.047 | 0.72 |
| L-IC | L-Pulvinar | 0.027 | 0.047 | 0.721 |
| L-cuneus | R-IC | 0.028 | 0.047 | 0.723 |
| R-Hippo_GCDG | R-IC | 0.027 | 0.047 | 0.723 |
| L-Hippo_CA3 | L-MGN | 0.028 | 0.047 | 0.724 |
| L-lingual | L-CN-SOC | 0.028 | 0.047 | 0.726 |
| L-cMFC | R-CN-SOC | 0.028 | 0.047 | 0.727 |
| L-Hippo_CA1 | L-PAC | 0.029 | 0.047 | 0.728 |
| L-PAC | R-CLLPM_Pulvinar | 0.028 | 0.047 | 0.728 |
| L-Hippo_Tail | R-CN-SOC | 0.029 | 0.047 | 0.73 |
| R-Hippo_Tail | R-CN-SOC | 0.028 | 0.047 | 0.73 |
| R-cuneus | R-IC | 0.028 | 0.047 | 0.73 |
| L-IC | R-pericalc | 0.029 | 0.047 | 0.732 |
| R-PCC | R-CN-SOC | 0.029 | 0.047 | 0.733 |
| L-Anterior | R-IC | 0.029 | 0.047 | 0.735 |
| R-Hippo_Tail | R-IC | 0.029 | 0.047 | 0.736 |
| R-IC | R-VLD | 0.029 | 0.047 | 0.737 |
| L-PAC | R-PCC | 0.029 | 0.047 | 0.738 |
| L-PAC | R-VLD | 0.03 | 0.047 | 0.738 |
| L-MGN | L-pericalc | 0.03 | 0.047 | 0.74 |
| L-PCC | R-IC | 0.03 | 0.047 | 0.74 |
| R-CLLPM_Pulvinar | R-IC | 0.03 | 0.047 | 0.74 |
| L-Caudate | L-CN-SOC | 0.03 | 0.047 | 0.742 |
| **L-PAC** | L-parso | 0.031 | 0.047 | 0.743 |
| **L-Hippo_CA1** | L-IC | 0.03 | 0.047 | 0.744 |
| **L-IC** | R-lingual | 0.03 | 0.047 | 0.744 |
| **L-IPC** | R-CN-SOC | 0.031 | 0.047 | 0.748 |
| **L-Hippo_CA3** | L-CN-SOC | 0.032 | 0.047 | 0.751 |
| **L-Anterior** | R-CN-SOC | 0.032 | 0.047 | 0.753 |
| **R-lingual** | R-MGN | 0.032 | 0.047 | 0.754 |
| **R-Anterior** | R-CN-SOC | 0.032 | 0.047 | 0.755 |
| **L-PAC** | R-Caudate | 0.032 | 0.047 | 0.759 |
| **L-pericalc** | L-CN-SOC | 0.033 | 0.047 | 0.765 |
| **R-Caudate** | R-IC | 0.034 | 0.047 | 0.766 |
| **L-IC** | L-lingual | 0.033 | 0.047 | 0.767 |
| **R-cuneus** | R-CN-SOC | 0.033 | 0.047 | 0.767 |
| **R-Hippo_GCDG** | R-CN-SOC | 0.034 | 0.047 | 0.767 |
| **L-PAC** | L-PCC | 0.034 | 0.047 | 0.77 |
| **R-CLLPM_Pulvinar** | R-CN-SOC | 0.034 | 0.047 | 0.77 |
| **L-pericalc** | R-MGN | 0.034 | 0.047 | 0.77 |
| **L-Hippo_CA3** | L-PAC | 0.035 | 0.047 | 0.773 |
| **R-CN-SOC** | R-V_Anterior | 0.035 | 0.047 | 0.773 |
| **L-Hippo_CA3** | R-MGN | 0.034 | 0.047 | 0.774 |
| **L-IC** | R-CN-SOC | 0.035 | 0.047 | 0.776 |
| **L-PCC** | R-CN-SOC | 0.037 | 0.047 | 0.786 |
| **L-cuneus** | R-CN-SOC | 0.038 | 0.047 | 0.792 |
| **L-lingual** | R-MGN | 0.037 | 0.047 | 0.792 |
| **R-cMFC** | R-IC | 0.04 | 0.047 | 0.806 |
| **L-PAC** | L-V_Anterior | 0.04 | 0.047 | 0.807 |
| **L-PAC** | R-pericalc | 0.04 | 0.047 | 0.808 |
| **L-Caudate** | L-IC | 0.04 | 0.047 | 0.809 |
| **R-IC** | R-pericalc | 0.041 | 0.047 | 0.813 |
| **L-PAC** | R-cMFC | 0.041 | 0.047 | 0.814 |
| **L-V_Anterior** | R-IC | 0.042 | 0.047 | 0.814 |
| **R-CN-SOC** | R-VLD | 0.041 | 0.047 | 0.814 |
| **L-Hippo_CA3** | L-IC | 0.042 | 0.047 | 0.817 |
| **L-Hippo_CA4** | R-IC | 0.042 | 0.047 | 0.817 |
| **R-Hippo_CA3** | R-IC | 0.042 | 0.047 | 0.817 |
| **L-IC** | L-pericalc | 0.042 | 0.047 | 0.819 |
| **L-Pulvinar** | R-IC | 0.042 | 0.047 | 0.82 |
| **L-Hippo_Pres** | R-CN-SOC | 0.043 | 0.047 | 0.823 |
| **R-IC** | R-CN-SOC | 0.044 | 0.047 | 0.828 |
| **L-Hippo_Pres** | R-IC | 0.044 | 0.047 | 0.828 |
| **R-Caudate** | R-CN-SOC | 0.045 | 0.047 | 0.835 |
| **L-PAC** | R-lingual | 0.046 | 0.047 | 0.839 |
| **R-Hippo_CA3** | R-CN-SOC | 0.047 | 0.047 | 0.842 |
| **L-Hippo_CA4** | R-CN-SOC | 0.047 | 0.047 | 0.844 |
| **L-Caudate** | L-PAC | 0.047 | 0.047 | 0.846 |
| **L-Hippo_CA1** | R-CN-SOC | 0.048 | 0.047 | 0.846 |
| **R-IC** | R-lingual | 0.047 | 0.047 | 0.846 |
| **L-Hippo_CA1** | R-IC | 0.048 | 0.047 | 0.849 |
| **L-V_Anterior** | R-CN-SOC | 0.049 | 0.047 | 0.851 |
| **L-lingual** | L-PAC | 0.049 | 0.047 | 0.855 |
| **L-lingual** | R-IC | 0.049 | 0.047 | 0.857 |
| **R-cMFC** | R-CN-SOC | 0.051 | 0.047 | 0.861 |
| **R-lingual** | R-CN-SOC | 0.051 | 0.047 | 0.862 |
| **L-Pulvinar** | R-CN-SOC | 0.052 | 0.047 | 0.869 |
| **R-pericalc** | R-CN-SOC | 0.052 | 0.047 | 0.869 |
| **L-PAC** | L-pericalc | 0.053 | 0.047 | 0.873 |
| **L-CN-SOC** | R-CN-SOC | 0.055 | 0.047 | 0.882 |
| **L-Caudate** | R-IC | 0.057 | 0.047 | 0.894 |
| **L-pericalc** | R-IC | 0.059 | 0.047 | 0.898 |
| **L-lingual** | R-CN-SOC | 0.06 | 0.047 | 0.9 |
| **L-Hippo_CA3** | R-IC | 0.061 | 0.047 | 0.906 |
| **L-pericalc** | R-CN-SOC | 0.066 | 0.047 | 0.918 |
| **L-Caudate** | R-CN-SOC | 0.066 | 0.047 | 0.922 |
| **L-Hippo_CA3** | R-CN-SOC | 0.069 | 0.047 | 0.929 |
| **L-IC** | R-IC | 0.08 | 0.048 | 0.954 |

## Appendix E: RS-fMRI-based functional network

|  | degree |  |  |  | strength |  |  |  | transitivity |  |  |  | Nodal efficiency |  |  |  | Local efficiency |  |  |
| --- | --- | --- | --- | --- | --- | --- | --- | --- | --- | --- | --- | --- | --- | --- | --- | --- | --- | --- | --- |
| ROI | mean | SD | P+ |  | mean | SD | P+ |  | mean | SD | P+ |  | mean | SD | P+ |  | mean | SD | P+ |
| L Anterior | 0,526 | 4,510 | 0,550 |  | 0,070 | 2,191 | 0,486 |  | 0,005 | 0,061 | 0,534 |  | 0,059 | 0,079 | 0,780 |  | 0,070 | 0,062 | 0,872 |
| L bankssts | 0,112 | 4,372 | 0,513 |  | 0,050 | 2,177 | 0,492 |  | 0,005 | 0,061 | 0,536 |  | 0,058 | 0,078 | 0,778 |  | 0,053 | 0,060 | 0,815 |
| L cACC | 0,071 | 4,371 | 0,509 |  | 0,035 | 2,177 | 0,494 |  | 0,006 | 0,061 | 0,538 |  | 0,053 | 0,079 | 0,758 |  | 0,048 | 0,060 | 0,790 |
| L Caudate | 0,378 | 4,415 | 0,536 |  | 0,015 | 2,178 | 0,498 |  | 0,005 | 0,061 | 0,537 |  | 0,063 | 0,078 | 0,796 |  | 0,053 | 0,060 | 0,814 |
| L CLLPM_Pulvinar | 0,011 | 4,379 | 0,502 |  | 0,105 | 2,187 | 0,484 |  | 0,005 | 0,061 | 0,535 |  | 0,063 | 0,079 | 0,792 |  | 0,055 | 0,060 | 0,820 |
| L cMFC | 0,103 | 4,368 | 0,512 |  | 0,028 | 2,178 | 0,494 |  | 0,006 | 0,061 | 0,540 |  | 0,062 | 0,079 | 0,787 |  | 0,051 | 0,060 | 0,805 |
| L cuneus | 0,125 | 4,372 | 0,513 |  | 0,033 | 2,177 | 0,495 |  | 0,006 | 0,061 | 0,542 |  | 0,055 | 0,079 | 0,765 |  | 0,053 | 0,060 | 0,815 |
| L Hippo | 0,216 | 4,518 | 0,486 |  | 0,059 | 2,181 | 0,491 |  | 0,006 | 0,061 | 0,541 |  | 0,057 | 0,078 | 0,774 |  | 0,063 | 0,061 | 0,854 |
| L Hippo_CA1 | 0,208 | 4,528 | 0,488 |  | 0,060 | 2,180 | 0,489 |  | 0,006 | 0,061 | 0,539 |  | 0,041 | 0,085 | 0,708 |  | 0,051 | 0,064 | 0,793 |
| L Hippo_CA3 | 0,239 | 4,492 | 0,523 |  | 0,025 | 2,180 | 0,497 |  | 0,005 | 0,061 | 0,532 |  | 0,048 | 0,080 | 0,738 |  | 0,062 | 0,063 | 0,839 |
| L Hippo_CA4 | 0,007 | 4,418 | 0,504 |  | 0,035 | 2,178 | 0,495 |  | 0,005 | 0,061 | 0,534 |  | 0,056 | 0,079 | 0,770 |  | 0,058 | 0,060 | 0,834 |
| L Hippo_Mol_layer_HP | 0,012 | 4,401 | 0,501 |  | 0,052 | 2,179 | 0,490 |  | 0,005 | 0,061 | 0,536 |  | 0,060 | 0,079 | 0,783 |  | 0,059 | 0,061 | 0,837 |
| L Hippo_Pres | 0,176 | 4,413 | 0,518 |  | 0,018 | 2,177 | 0,497 |  | 0,005 | 0,061 | 0,536 |  | 0,056 | 0,078 | 0,769 |  | 0,063 | 0,061 | 0,848 |
| L Hippo_Tail | 0,048 | 4,371 | 0,506 |  | 0,043 | 2,178 | 0,493 |  | 0,005 | 0,061 | 0,535 |  | 0,051 | 0,079 | 0,749 |  | 0,050 | 0,062 | 0,794 |
| L IC | 1,677 | 5,025 | 0,627 |  | 0,041 | 2,216 | 0,494 |  | 0,005 | 0,061 | 0,538 |  | 0,046 | 0,082 | 0,727 |  | 0,087 | 0,066 | 0,909 |
| L IPC | 0,226 | 4,392 | 0,524 |  | 0,017 | 2,180 | 0,497 |  | 0,005 | 0,061 | 0,536 |  | 0,064 | 0,078 | 0,797 |  | 0,054 | 0,060 | 0,818 |
| L Isth | 0,017 | 4,368 | 0,504 |  | 0,034 | 2,178 | 0,494 |  | 0,005 | 0,061 | 0,536 |  | 0,053 | 0,078 | 0,758 |  | 0,055 | 0,060 | 0,823 |
| L lingual | 0,258 | 4,396 | 0,525 |  | 0,022 | 2,178 | 0,497 |  | 0,006 | 0,061 | 0,538 |  | 0,049 | 0,080 | 0,740 |  | 0,056 | 0,061 | 0,826 |
| L LOC | 0,195 | 4,383 | 0,518 |  | 0,036 | 2,179 | 0,494 |  | 0,005 | 0,061 | 0,532 |  | 0,053 | 0,078 | 0,759 |  | 0,054 | 0,060 | 0,817 |
| L Medio_Dorsal | 0,013 | 4,395 | 0,503 |  | 0,088 | 2,181 | 0,485 |  | 0,006 | 0,061 | 0,541 |  | 0,064 | 0,081 | 0,793 |  | 0,053 | 0,060 | 0,814 |
| L MGN | 0,063 | 4,395 | 0,510 |  | 0,044 | 2,186 | 0,493 |  | 0,005 | 0,061 | 0,537 |  | 0,064 | 0,081 | 0,793 |  | 0,055 | 0,060 | 0,824 |
| L PAC | 0,012 | 4,413 | 0,502 |  | 0,026 | 2,177 | 0,496 |  | 0,005 | 0,061 | 0,536 |  | 0,049 | 0,081 | 0,737 |  | 0,060 | 0,061 | 0,842 |
| L Pallidum | 0,247 | 4,381 | 0,526 |  | 0,024 | 2,181 | 0,499 |  | 0,005 | 0,061 | 0,533 |  | 0,050 | 0,081 | 0,740 |  | 0,058 | 0,060 | 0,835 |
| L paracentral | 0,170 | 4,375 | 0,518 |  | 0,031 | 2,177 | 0,496 |  | 0,006 | 0,061 | 0,537 |  | 0,055 | 0,078 | 0,765 |  | 0,051 | 0,060 | 0,804 |
| L parso | 0,085 | 4,375 | 0,509 |  | 0,029 | 2,178 | 0,494 |  | 0,005 | 0,061 | 0,537 |  | 0,056 | 0,082 | 0,761 |  | 0,052 | 0,060 | 0,813 |
| L parstr | 0,189 | 4,376 | 0,520 |  | 0,026 | 2,176 | 0,497 |  | 0,006 | 0,061 | 0,538 |  | 0,059 | 0,078 | 0,779 |  | 0,052 | 0,060 | 0,811 |
| L PCC | 0,085 | 4,412 | 0,496 |  | 0,011 | 2,181 | 0,498 |  | 0,005 | 0,061 | 0,533 |  | 0,064 | 0,079 | 0,797 |  | 0,052 | 0,060 | 0,811 |
| L pericalc | 0,149 | 4,375 | 0,515 |  | 0,039 | 2,175 | 0,495 |  | 0,006 | 0,061 | 0,539 |  | 0,056 | 0,079 | 0,768 |  | 0,054 | 0,060 | 0,822 |
| L postcentral | 0,321 | 4,410 | 0,530 |  | 0,011 | 2,179 | 0,499 |  | 0,006 | 0,061 | 0,537 |  | 0,072 | 0,081 | 0,820 |  | 0,054 | 0,060 | 0,820 |
| L precentral | 0,115 | 4,377 | 0,513 |  | 0,040 | 2,181 | 0,492 |  | 0,005 | 0,061 | 0,536 |  | 0,082 | 0,088 | 0,834 |  | 0,054 | 0,060 | 0,816 |
| L precuneus | 0,076 | 4,368 | 0,509 |  | 0,016 | 2,182 | 0,498 |  | 0,005 | 0,061 | 0,536 |  | 0,052 | 0,079 | 0,753 |  | 0,053 | 0,060 | 0,814 |
| L Pulvinar | 4,273 | 5,917 | 0,762 |  | 0,046 | 2,231 | 0,509 |  | 0,005 | 0,061 | 0,531 |  | 0,032 | 0,129 | 0,607 |  | 0,127 | 0,073 | 0,960 |
| L Putamen | 0,003 | 4,381 | 0,501 |  | 0,044 | 2,179 | 0,493 |  | 0,005 | 0,061 | 0,536 |  | 0,030 | 0,090 | 0,659 |  | 0,054 | 0,060 | 0,818 |
| L rACC | 0,084 | 4,369 | 0,509 |  | 0,024 | 2,176 | 0,497 |  | 0,006 | 0,061 | 0,539 |  | 0,049 | 0,080 | 0,738 |  | 0,052 | 0,060 | 0,809 |
| L rMFC | 0,041 | 4,368 | 0,506 |  | 0,024 | 2,178 | 0,496 |  | 0,006 | 0,061 | 0,539 |  | 0,051 | 0,081 | 0,744 |  | 0,048 | 0,061 | 0,792 |
| L SFC | 0,014 | 4,391 | 0,504 |  | 0,027 | 2,182 | 0,493 |  | 0,006 | 0,061 | 0,540 |  | 0,049 | 0,079 | 0,741 |  | 0,046 | 0,060 | 0,781 |
| L CN/SOC | 0,601 | 4,904 | 0,549 |  | 0,038 | 2,225 | 0,508 |  | 0,006 | 0,061 | 0,538 |  | 0,063 | 0,079 | 0,790 |  | 0,080 | 0,068 | 0,883 |
| L SPC | 0,086 | 4,377 | 0,511 |  | 0,059 | 2,180 | 0,489 |  | 0,005 | 0,061 | 0,537 |  | 0,062 | 0,078 | 0,792 |  | 0,053 | 0,060 | 0,815 |
| L V_Anterior | 0,169 | 4,378 | 0,519 |  | 0,044 | 2,179 | 0,492 |  | 0,006 | 0,061 | 0,538 |  | 0,062 | 0,078 | 0,792 |  | 0,054 | 0,060 | 0,821 |
| L VLD | 0,095 | 4,370 | 0,510 |  | 0,052 | 2,178 | 0,492 |  | 0,005 | 0,061 | 0,537 |  | 0,064 | 0,080 | 0,795 |  | 0,056 | 0,060 | 0,827 |
| R Anterior | 0,280 | 4,386 | 0,527 |  | 0,042 | 2,178 | 0,493 |  | 0,005 | 0,061 | 0,536 |  | 0,071 | 0,081 | 0,814 |  | 0,052 | 0,060 | 0,809 |
| R bankssts | 0,146 | 4,371 | 0,516 |  | 0,042 | 2,177 | 0,494 |  | 0,005 | 0,061 | 0,531 |  | 0,068 | 0,079 | 0,808 |  | 0,054 | 0,060 | 0,819 |
| R cACC | 0,067 | 4,371 | 0,508 |  | 0,041 | 2,177 | 0,494 |  | 0,005 | 0,061 | 0,536 |  | 0,058 | 0,078 | 0,777 |  | 0,050 | 0,060 | 0,802 |
| R Caudate | 0,134 | 4,368 | 0,515 |  | 0,037 | 2,177 | 0,494 |  | 0,005 | 0,061 | 0,531 |  | 0,063 | 0,078 | 0,795 |  | 0,051 | 0,060 | 0,808 |
| R CLLPM_Pulvinar | 0,147 | 4,371 | 0,516 |  | 0,039 | 2,176 | 0,494 |  | 0,004 | 0,061 | 0,529 |  | 0,067 | 0,080 | 0,804 |  | 0,049 | 0,060 | 0,797 |
| R cMFC | 0,015 | 4,378 | 0,501 |  | 0,034 | 2,178 | 0,494 |  | 0,005 | 0,061 | 0,533 |  | 0,064 | 0,079 | 0,795 |  | 0,049 | 0,060 | 0,797 |
| R cuneus | 0,149 | 4,373 | 0,515 |  | 0,045 | 2,176 | 0,493 |  | 0,004 | 0,061 | 0,528 |  | 0,058 | 0,078 | 0,777 |  | 0,057 | 0,060 | 0,832 |
| R Hippo | 0,155 | 4,449 | 0,519 |  | 0,054 | 2,184 | 0,491 |  | 0,005 | 0,061 | 0,531 |  | 0,071 | 0,080 | 0,815 |  | 0,057 | 0,060 | 0,831 |
| R Hippo_CA1 | 0,228 | 4,473 | 0,484 |  | 0,075 | 2,184 | 0,488 |  | 0,006 | 0,061 | 0,538 |  | 0,055 | 0,078 | 0,766 |  | 0,057 | 0,060 | 0,828 |
| R Hippo_CA3 | 1,058 | 4,566 | 0,591 |  | 0,010 | 2,179 | 0,500 |  | 0,004 | 0,061 | 0,530 |  | 0,070 | 0,080 | 0,813 |  | 0,058 | 0,060 | 0,831 |
| R Hippo_CA4 | 0,016 | 4,403 | 0,502 |  | 0,047 | 2,178 | 0,493 |  | 0,006 | 0,061 | 0,540 |  | 0,051 | 0,079 | 0,745 |  | 0,060 | 0,061 | 0,840 |
| R Hippo_GCDG | 0,022 | 4,415 | 0,502 |  | 0,051 | 2,177 | 0,491 |  | 0,005 | 0,061 | 0,533 |  | 0,058 | 0,078 | 0,779 |  | 0,059 | 0,060 | 0,836 |
| R Hippo_Tail | 0,012 | 4,377 | 0,505 |  | 0,047 | 2,177 | 0,493 |  | 0,006 | 0,061 | 0,541 |  | 0,056 | 0,078 | 0,770 |  | 0,054 | 0,060 | 0,818 |
| R IC | 0,098 | 4,528 | 0,514 |  | 0,086 | 2,202 | 0,486 |  | 0,006 | 0,061 | 0,541 |  | 0,005 | 0,111 | 0,520 |  | 0,125 | 0,071 | 0,962 |
| R IPC | 0,093 | 4,368 | 0,511 |  | 0,045 | 2,178 | 0,492 |  | 0,006 | 0,061 | 0,536 |  | 0,084 | 0,087 | 0,841 |  | 0,053 | 0,060 | 0,814 |
| R Isth | 0,087 | 4,369 | 0,511 |  | 0,029 | 2,176 | 0,495 |  | 0,005 | 0,061 | 0,532 |  | 0,065 | 0,078 | 0,803 |  | 0,050 | 0,060 | 0,803 |
| R lingual | 0,129 | 4,377 | 0,516 |  | 0,039 | 2,177 | 0,493 |  | 0,006 | 0,061 | 0,538 |  | 0,072 | 0,082 | 0,820 |  | 0,049 | 0,060 | 0,799 |
| R LOC | 0,169 | 4,372 | 0,519 |  | 0,030 | 2,176 | 0,495 |  | 0,005 | 0,061 | 0,534 |  | 0,051 | 0,079 | 0,750 |  | 0,050 | 0,060 | 0,802 |
| R Medio_Dorsal | 0,196 | 4,421 | 0,485 |  | 0,080 | 2,179 | 0,487 |  | 0,006 | 0,061 | 0,538 |  | 0,054 | 0,078 | 0,760 |  | 0,056 | 0,060 | 0,828 |
| R MGN | 0,041 | 4,412 | 0,501 |  | 0,109 | 2,193 | 0,480 |  | 0,006 | 0,061 | 0,538 |  | 0,076 | 0,093 | 0,802 |  | 0,073 | 0,062 | 0,880 |
| R MTC | 0,200 | 4,389 | 0,520 |  | 0,006 | 2,181 | 0,499 |  | 0,006 | 0,061 | 0,537 |  | 0,056 | 0,082 | 0,760 |  | 0,052 | 0,060 | 0,808 |
| R PAC | 0,091 | 4,377 | 0,512 |  | 0,023 | 2,180 | 0,497 |  | 0,006 | 0,061 | 0,538 |  | 0,036 | 0,137 | 0,466 |  | 0,058 | 0,061 | 0,830 |
| R Pallidum | 0,377 | 4,395 | 0,537 |  | 0,014 | 2,178 | 0,499 |  | 0,005 | 0,061 | 0,529 |  | 0,066 | 0,079 | 0,799 |  | 0,050 | 0,060 | 0,799 |
| R paracentral | 0,108 | 4,368 | 0,513 |  | 0,055 | 2,178 | 0,490 |  | 0,005 | 0,061 | 0,536 |  | 0,070 | 0,080 | 0,816 |  | 0,051 | 0,060 | 0,809 |
| R pars | 0,194 | 4,378 | 0,520 |  | 0,015 | 2,181 | 0,497 |  | 0,005 | 0,061 | 0,536 |  | 0,064 | 0,079 | 0,795 |  | 0,050 | 0,060 | 0,801 |
| R parso | 0,037 | 4,381 | 0,499 |  | 0,043 | 2,181 | 0,491 |  | 0,005 | 0,061 | 0,535 |  | 0,058 | 0,077 | 0,778 |  | 0,050 | 0,060 | 0,799 |
| R parstr | 0,128 | 4,376 | 0,514 |  | 0,005 | 2,180 | 0,500 |  | 0,006 | 0,061 | 0,538 |  | 0,057 | 0,080 | 0,770 |  | 0,049 | 0,060 | 0,795 |
| R PCC | 0,015 | 4,373 | 0,504 |  | 0,046 | 2,179 | 0,491 |  | 0,005 | 0,061 | 0,535 |  | 0,065 | 0,078 | 0,804 |  | 0,053 | 0,060 | 0,815 |
| R pericalc | 0,074 | 4,368 | 0,508 |  | 0,057 | 2,179 | 0,491 |  | 0,005 | 0,061 | 0,537 |  | 0,054 | 0,079 | 0,763 |  | 0,053 | 0,060 | 0,812 |
| R postcentral | 0,214 | 4,371 | 0,522 |  | 0,027 | 2,179 | 0,497 |  | 0,005 | 0,061 | 0,534 |  | 0,069 | 0,079 | 0,813 |  | 0,054 | 0,060 | 0,820 |
| R precentral | 0,153 | 4,373 | 0,515 |  | 0,045 | 2,178 | 0,492 |  | 0,006 | 0,061 | 0,538 |  | 0,068 | 0,079 | 0,810 |  | 0,053 | 0,060 | 0,814 |
| R precuneus | 0,186 | 4,369 | 0,518 |  | 0,023 | 2,178 | 0,496 |  | 0,005 | 0,061 | 0,535 |  | 0,076 | 0,083 | 0,828 |  | 0,055 | 0,060 | 0,824 |
| R Putamen | 0,181 | 4,370 | 0,519 |  | 0,037 | 2,178 | 0,495 |  | 0,005 | 0,061 | 0,530 |  | 0,069 | 0,080 | 0,810 |  | 0,051 | 0,060 | 0,806 |
| R rACC | 0,114 | 4,376 | 0,513 |  | 0,036 | 2,176 | 0,494 |  | 0,006 | 0,061 | 0,538 |  | 0,074 | 0,082 | 0,824 |  | 0,050 | 0,060 | 0,803 |
| R rMFC | 0,139 | 4,374 | 0,514 |  | 0,027 | 2,180 | 0,494 |  | 0,006 | 0,061 | 0,539 |  | 0,077 | 0,083 | 0,831 |  | 0,049 | 0,060 | 0,797 |
| R SFC | 0,140 | 4,383 | 0,517 |  | 0,012 | 2,183 | 0,498 |  | 0,005 | 0,061 | 0,533 |  | 0,059 | 0,078 | 0,779 |  | 0,050 | 0,060 | 0,801 |
| **R CN/SOC** | 1,533 | 5,884 | 0,601 |  | 0,089 | 2,224 | 0,485 |  | 0,005 | 0,061 | 0,531 |  | 0,059 | 0,080 | 0,777 |  | 0,127 | 0,073 | 0,963 |
| **R SPC** | 0,079 | 4,375 | 0,512 |  | 0,046 | 2,179 | 0,492 |  | 0,005 | 0,061 | 0,536 |  | 0,065 | 0,079 | 0,800 |  | 0,052 | 0,060 | 0,814 |
| **R V_Anterior** | 0,080 | 4,366 | 0,510 |  | 0,046 | 2,178 | 0,491 |  | 0,005 | 0,061 | 0,532 |  | 0,057 | 0,078 | 0,772 |  | 0,052 | 0,060 | 0,810 |
| **R VLD** | 0,114 | 4,370 | 0,513 |  | 0,043 | 2,177 | 0,493 |  | 0,006 | 0,061 | 0,538 |  | 0,056 | 0,079 | 0,768 |  | 0,053 | 0,060 | 0,814 |

## Notes

### Competing Interest Statement

The authors have declared no competing interest.

